# The porphyran degradation system is complete, phylogenetically and geographically diverse across the gut microbiota of East Asian populations

**DOI:** 10.1101/2023.03.30.534863

**Authors:** Laure Ségurel, Thirumalai Selvi Ulaganathan, Sophie Mathieu, Mélanie Touvrey, Laurent Poulet, Sophie Drouillard, Miroslaw Cygler, William Helbert

## Abstract

The human gut microbiota can acquire new catabolic functions by integrating genetic material coming from the environment, for example from food-associated bacteria. The most illustrative example is the acquisition by the human gut microbiota of Asian populations of genes coming from marine bacteria living on the surface of red algae that are incorporated into their diet when eating maki-sushi. To better understand the function and evolution of this set of algal genes corresponding to a polysaccharide utilization locus (PUL) dedicated to the degradation of porphyran, the main polysaccharide of the red algae *Porphyra sp*., we characterized it biochemically, assessed its genetic diversity and investigated its geographical distribution in large public worldwide datasets. We first demonstrated that both methylated and unmethylated fractions are catabolized without the help of external enzymes. By scanning the genomic data of more than 10,000 cultivated isolates, we then found that the porphyran PUL organization is conserved in 22 different *Bacteroides* strains coming from at least 8 species, highlighting multiple lateral transfers within the gut microbiota. We then analyzed the metagenomic data of more than 14,000 individuals coming from 32 countries worldwide and showed that the porphyran PUL exists only in East Asia (Japan, China, Korea), but not anywhere else. Finally, we identified three major PUL haplotypes which frequency differ between countries. This geographic structure is likely the reflect of the rate of bacterial horizontal transmission between individuals.

**Background:** The human gut microbiota can acquire new catabolic functions by integrating genetic material coming from the environment, for example from food-associated bacteria. The most illustrative example is the acquisition by the gut microbiota of Asian populations of genes coming from marine bacteria living on the surface of red algae that are incorporated into their diet when eating maki-sushi. To better understand the function and evolution of this set of algal genes corresponding to a polysaccharide utilization locus (PUL) dedicated to the degradation of porphyran, the main polysaccharide of the red algae *Porphyra sp*., we characterized it biochemically, assessed its genetic diversity and investigated its geographical distribution in large public worldwide datasets.

**Results:** We first demonstrated that both methylated and unmethylated fractions of porphyran are catabolized by the porphyran PUL without the help of external enzymes. By scanning the genomic data of more than 10,000 cultivated isolates, we then found that the porphyran PUL organization is conserved in 22 different *Bacteroides* strains coming from at least 8 species, highlighting multiple lateral transfers within the gut microbiota. We then analyzed the metagenomic data of more than 14,000 individuals coming from 32 countries worldwide and confirmed that the porphyran PUL exists only in East Asia (Japan, China, Korea). We identified three major porphyran PUL haplotypes which frequency differ between countries.

**Conclusion:** The encoded genes of the PUL porphyran can autonomously catabolized all the complex porphyran structure. The PUL is further encoded by a variety of bacterial species, and its genetic diversity is geographically structured, likely reflecting the rate of bacterial horizontal transmission between individuals.

## Introduction

The impact of diet on the composition of the bacterial communities inhabiting the human gut is now well documented [1,2]. For example, the ratio of Firmicutes to Bacteroidetes decreases with calorie-restricted or high-fiber diets, and the abundance of *Prevotella* is associated with the intake of carbohydrates while that of *Bacteroides* is associated with animal protein and fat [3,4,5]. In parallel, the gut microbiota can also acquire new functional capacities by horizontal transfer of genetic material between species of the gut microbiota but also from species found in the food ingested by the host [6,7,8]. One of the most illustrative example of how the food participates in shaping the human gut microbiota, at the molecular level, is the discovery of the horizontal gene transfer of a set of enzymes involved in the degradation of porphyran, the cell wall polysaccharide of the red algae *Porphyra sp.* used to prepare the maki-sushi, from a marine species living at the surface of the algae to the gut bacteria *Bacteroides plebeius* (now *Phocaeicola plebeius*) found in the Japanese gut microbiota [9]. Since this discovery, other algal polysaccharides degradation systems (i.e. agarose, alginate, carrageenan), likely acquired by lateral transfer from marine organisms, have also been studied [9,10,11,12,13].

Porphyran is an agar-type polysaccharide made of two disaccharides repetition: agarobiose and porphyranobiose decorated up to 64% by methyl groups [14,15] (Figure 1A). In *B. plebieus*, the genes encoding the enzymes degrading porphyran are co-localized and co-regulated in a so-called “polysaccharide utilizing loci” (PUL). The porphyran PUL identified in *B. plebieus* contains genes encoding two β-porphyranases (BpGH16B, Bacple_01689; BpGH86A, Bacple_01693) and two β-agarases (BpGH16A; Bacple_01670; BpGH86B, Bacple_01694) [9,16] (Figure 1B). The oligosaccharides resulting from the degradation of porphyran by these enzymes are then degraded further by exo-acting enzymes including sulfatase (BpS1_11, Bacple_01701), α-L-galactosidase (BpGH29, Bacple_01702) and β-D-galactosidase (BpGH2C, Bacple_01706) produced by *B. plebeius* or another agarolytic strain such as *Bacteroides uniformis* [13] (Figure 1B). The presence of a gene (BpGH50, Bacple_01683) encoding an enzyme grouped in the GH50 family known to contain agarases, was previously studied but its substrate specificity was not demonstrated [17]. Altogether, the many enzymes of the porphyran PUL (i.e. glycoside hydrolases and sulfatases) have been characterized, explaining the degradation process of the agarose components and the non-methylated fractions of porphyran by *B. plebieus* [9,16,18] (Figure 1B). However, the enzymes involved in the degradation of abundant fraction of the methylated porphyran have not been discovered yet.

**Figure 1:**
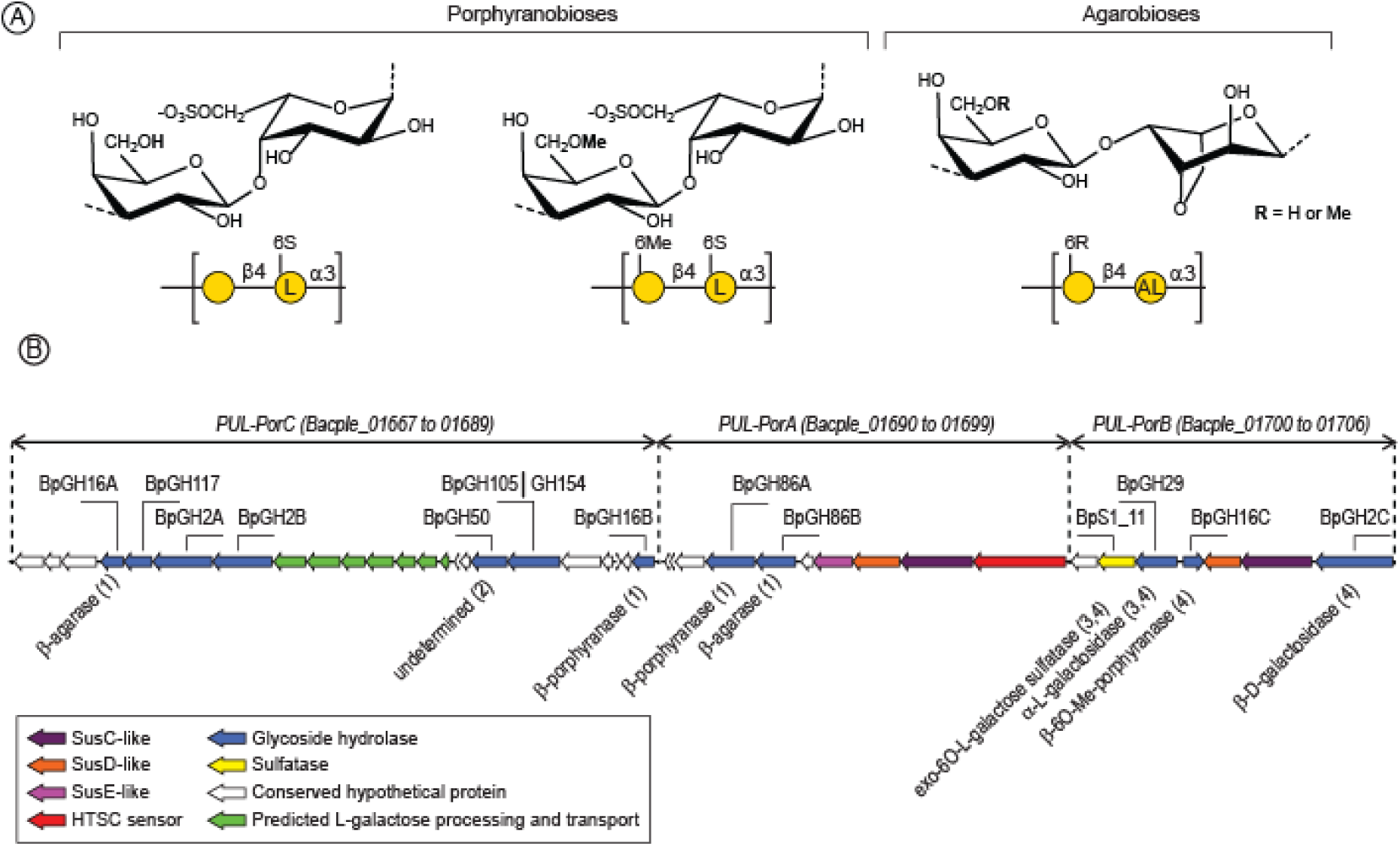
**A)** Chemical structure of the repetition moieties of porphyran. The sulfated disaccharides - porphyranobioses – can present methyl group at the position 6 of the D-galactose giving methylated and none-methylated porphyran components. Agarobiose units, obtained by desulfation/cyclization of the L-galactose residue during the biosynthesis, are methylated accordingly. **B)** Organization of the *B. plebieus* porphyran PUL. The PUL is divided in three segments (*PUL-PorA*, *-Por B* and *-PorC*) based on previous transcriptomic analyses. The cluster Bacple_01692 to Bacple_01699 genes – named *PUL-PorA*– was moderately up-regulated when *B. plebieus* was grown in the presence of porphyran. This contrasts with the two neighboring clusters of genes: Bacple_01668 to Bacple_01689 (*PUL-PorC*) and Bacple_01700 to Bacple_01706 (*PUL-PorB*), which were highly up-regulated (10-fold more than *PUL-PorA*) [16]. The enzyme functions were determined in (1) [16], (2) [17], [18] and (4) this study.

Even though the fronds of *Porphyra sp*. are nowadays mostly consumed by Asian populations for their taste and nutritional value [19], the most ancient proof that humans harvested and stored *Porphyra sp.* was found in the archaeological site of Monte Verde (Chile) occupied by humans 12,300 years ago [20]. The traditional use of *Porphyra spp.* by the first nations living along the Pacific coast of North America was also documented by ethnobotanists [21,22,23,24]. In South America, pre-Inca and Inca populations also incorporated red algae in their diet [25]. The very ancient know-how and use of *Porphyra sp.*, preserved for millennia along the Pacific Ocean’s coast, suggests that several lateral transfer events of genetic material dedicated to porphyran degradation from marine bacteria to gut bacteria might have occurred in ancient human populations. Furthermore, Europeans populations (from Spain to Scotland to Lithuania) have recently been found to have consumed seaweed from the Neolithic transition thousands of years ago up to the early Middle Ages [26], raising the question of whether these horizontally-transferred genes are found in different places across the globe. Originally, the porphyran PUL was identified in individuals from Japan, and has since been shown to be present also in China, but not in Denmark, Spain, USA, Tanzania or Brazil (survey of a dataset of 2,440 individuals) [13]. However, more data (covering more than 14,000 individuals from 32 different countries) is now available in the literature to tackle that question in more detail.

The number of human gut species carrying this metabolic capacity, allowing to make hypotheses about the origin of the algal degradation system in the human gut and the timing of these transfers, is not fully resolved either. A previous study of *in vitro* cultivation of stool samples of 240 donors in the presence of porphyran indeed showed that four Bacteroides species, as well as *Faecalicatena contorta* (from the Firmicutes phylum), were able to grow on porphyran [13], but a large-scale survey of existing gut bacterial genomes has not been performed yet.

We thus aimed to continue the detailed analysis of the porphyran PUL of *B. plebeius* by biochemically studying the uncharacterized glycoside hydrolases BpGH16C (Bacple_01703) and BpGH2C (Bacple_01706) and by completing the structural data of the sulfatase BpS1_11. We further took advantage of the increasingly large genomic datasets available in the literature to survey the genomes of more than 10,000 gut isolates, as well as the metagenomes of 14,000 worldwide individuals, in order i) to evaluate the number of gut bacterial species carrying the porphyran PUL, ii) to identify the number of countries where the porphyran PUL is present, and iii) to characterize the genetic diversity of porphyran PUL and its geographical differentiation.

## Results

### Biochemical and molecular characterization of the endo-acting 6O-methyl-porphyranase BpGH16C (Bacple_01703)

The gene Bacple_01703 encodes a glycoside hydrolase of the GH16 family (BpGH16C), which includes characterized agarases in sub-families GH16_15-16, and β-porphyranases in sub-families GH16_11-12 [27]. BpGH16C was classified in the GH16_14 sub-family that contains one characterized β-agarase (*Vibrio sp*. strain PO-303, BAF62129.1) [28] and one endo-6O-methyl-β-porphyranase (*Wenyingzhuangia fucanilytica*, WP_068825734.1) [29]. We assayed BpGH16C on various marine polysaccharides including carrageenan, agarose and porphyran. Only porphyran was degraded, leading to the production of a series of oligosaccharides characteristic of endo-acting glycoside hydrolase (Figure 2A). The structural characterization by NMR of the end-products purified by chromatography [30] revealed the occurrence of a methyl group on the β-linked D-galactose residue (Figure 2B), demonstrating that BpGH16C is an endo-6O-methyl-β-porphyranase able to accommodate the methyl group of porphyran in its active site. For comparison, we have examined the substrate specificities of one another predicted GH16_14 endo-6O-methyl-β-porphyranase from *Paraglaciecola atlantica* T6c (Patl_0824, ABG39352.1) presenting 51.1% identity with the BpGH16C and one GH16_12 β-porphyranase (Patl_0805, ABG39333.1) homologous to BpGH16B. Analyses of the degradation products confirmed the different porphyranase specificities (Figure 2A).

**Figure 2:**
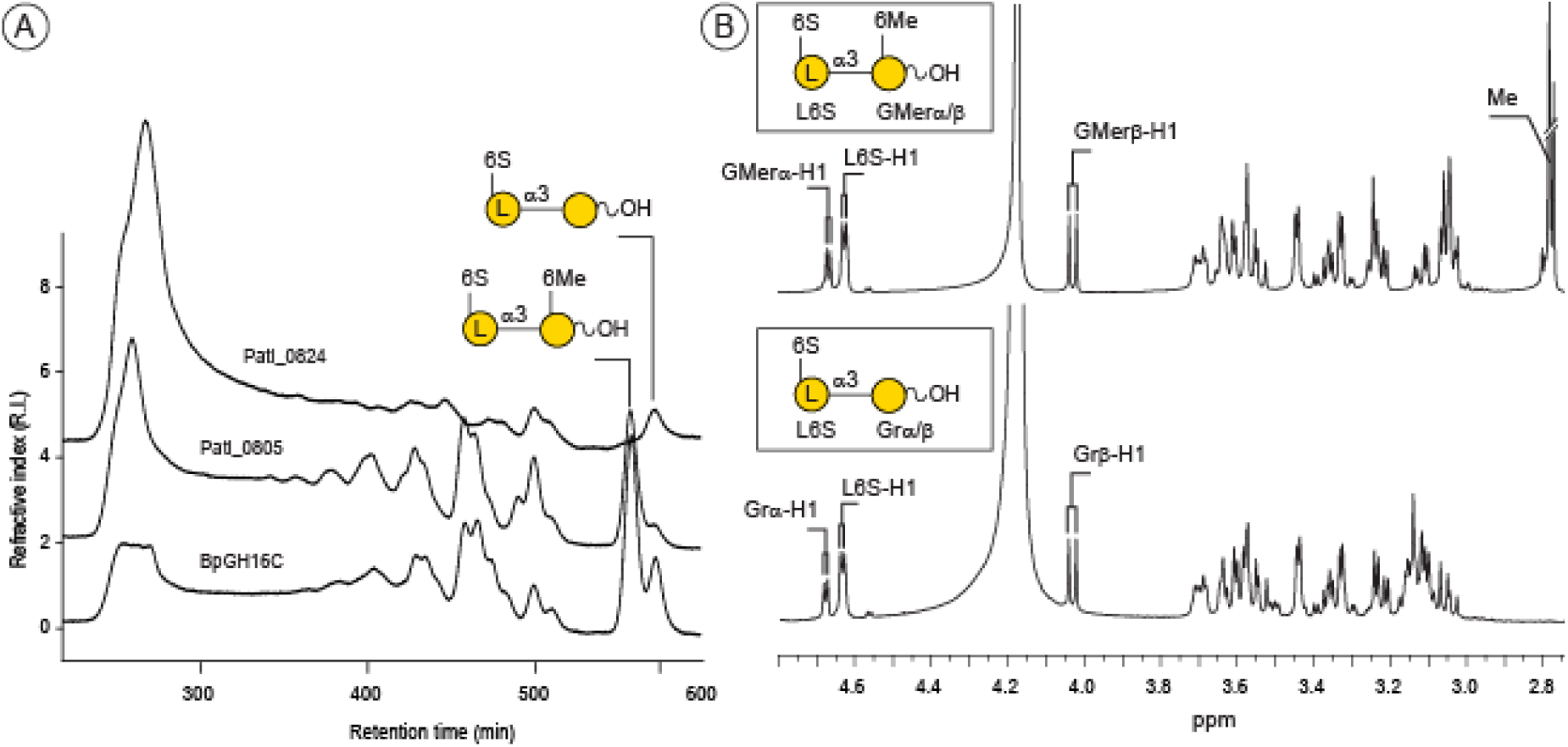
**A)** Size exclusion chromatography of the degradation products of porphyran incubated with the methyl-6O-β-porphyranase BpGH16C. The degradation profile was compared with the two *P. atlantica* T6c porphyranases grouped in the GH16_12 (Patl_0824) and GH16_14 (Patl_0805) sub-families. **B)** ^1^H NMR of the disaccharides end-products of BpGH16C.

We determined the crystal structure of BpGH16C at 1.9 Å resolution (PDB ID 8EP4) (Figure 3A). The enzyme has a β-sandwich jelly-roll fold with two stacking antiparallel β-sheets typical of proteins from GH16 family [27] and like in most GH16, a Ca^2+^ is bound on the convex side of the β-sandwich [31]. One molecule of Hepes from the crystallization buffer bound in the center of the cleft, with the 2-hydroxyethyl end buried into the molecule interior and the sulfate group exposed on protein surface (Figure 3A). To get a better understanding of substrate binding and catalysis, we crystallized the E145L inactive mutant with the tetrasaccharide L-α-6O-sulfate-Gal-(1→3)-D-β-Gal-(1→4)-L-α-6O-sulfate-Gal-(1→3)-D-a/β-Gal and determined its structure at 1.8 Å resolution (PDB ID 8EW1). An electron density was present within the groove in all the three independent molecules in the asymmetric unit fitted a D-β-Gal-(1→4)-L-α-6O-sulfate-Gal-(1→3)-D-β-Gal trisaccharide (Figure 3B). The electron density for the D-Gal on the nonreducing end is weaker than for the first two residues and no density is observed for the fourth residue indicating its mobility in the crystal. The residues correspond to the −1, −2 and −3 positions according to the standard nomenclature [32].

**Figure 3:**
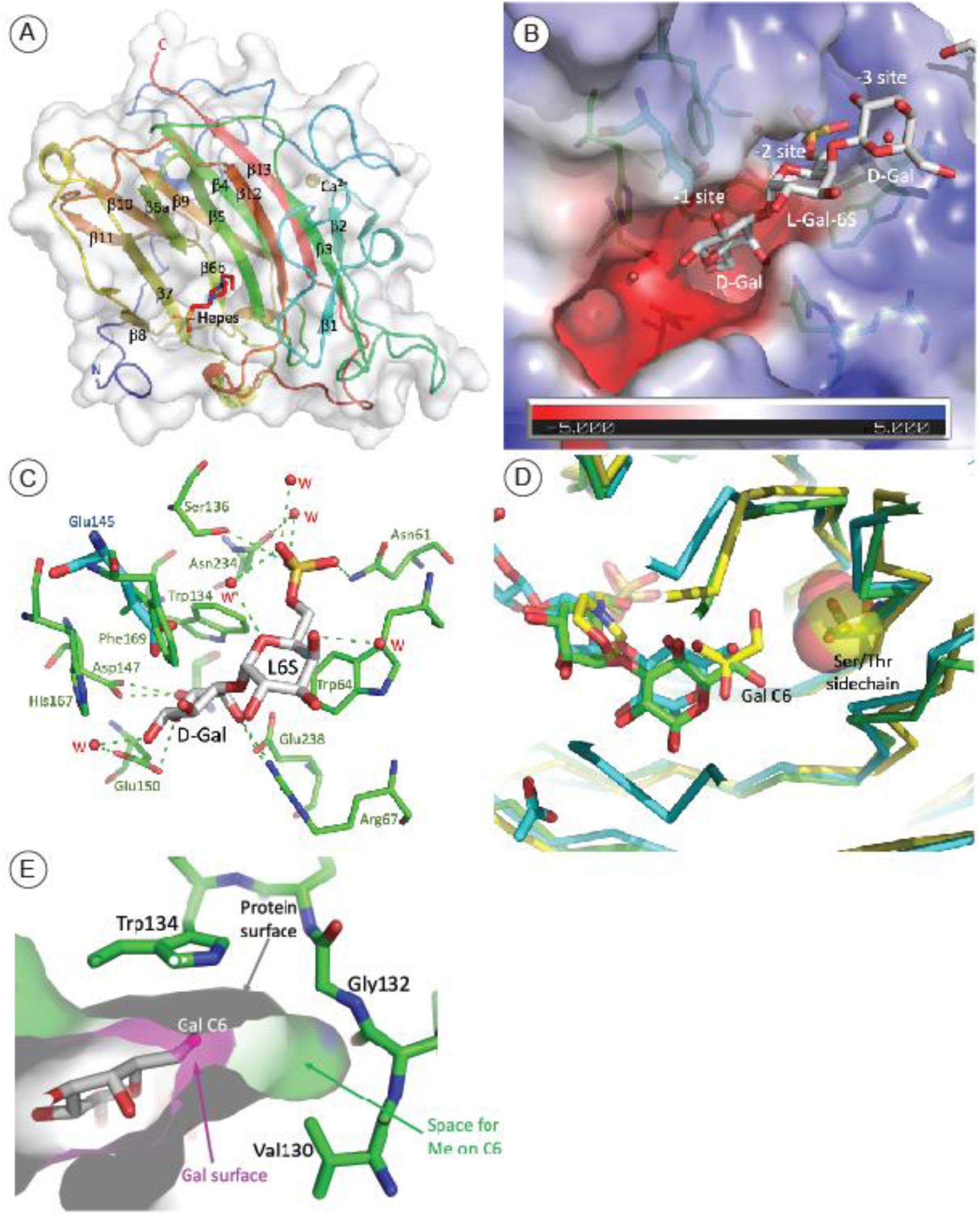
**A)** Crystal structure of BpGH16C (Bacple_01703). Chain A of asymmetric unit is shown in cartoon representation and colored in spectrum from N to C terminal. The HEPES molecule in the active site is shown as stick representation. The N- and C-terminals are marked, and the strands are labeled in their order in the sequence. **B)** The substrate binding site in the mutant BpGH16C-E145L complexed with the D-Gal-β4-6-sulfate-L-Gal disaccharide end-product. The semitransparent surface of the protein is colored by the electrostatic potential. The residues forming the site are shown in stick representation. The sulfate group from L-galactose 6 sulfate is docked into a positively charged pocket. **C)** The hydrogen bonds between the disaccharide in substrate binding site residues. Several H-bonds are bridged by water molecules (W). Glu145 (in blue and thicker bonds) from the native structure is superimposed on the Leu145 in the mutants. **D)** Structure comparison of BpGH16C (green), PorA (PDB id-3ILF) (magenta) and PorB (PDB id-3JUU) (yellow) showing the active site region. The single mutation of Ser129/Thr137 (PorA/PorB) (sidechains shown as spheres) to Gly132 in BpGH16C provides space for accommodating a methyl group at C6 on L-galactose residue. **E)** A cross-section of the surface representation of the Gal sugar and the cavity in the binding site near Gly132. There is free space within the cavity sufficient to accommodate the methyl group of a methylated porphyran.

The trisaccharide sits edge-on in the groove with C5 substituent of the ring in the −1 site directed toward the bottom of the cleft while O1 and O2 hydroxyls point toward the solvent. Trp134 stacks against hydrophobic side of the −1 residue while Arg67 and Glu238 hydrogen bond to the O4 hydroxyl. Residue in the −2 subsite is stacked between Trp64 and the edges of Trp143 and Phe169. Finally, the sulfate group of the −2 residue is hydrogen bonded to Asn61, Ser136 and Asn234 through a bridging water molecule. The D-Gal on the non-reducing end (- 3 site) has somewhat different orientation in molecules A/B and C and makes only one bridging hydrogen bond through a bridging water (Figure 3C). The structure of the complex with a trisaccharide adds also to our understanding of the catalytic mechanism. Glu150 is indeed the closest to the O1 hydroxyl that formed the bond to the next sugar eliminated by hydrolysis, while Glu145 approaches from the opposite side of the ring relative the O1 hydroxyl (Figure 3C) and is likely helping in proper orientation of the substrate. Asp147 is directed to the bridging O1 and might be helping in catalysis.

Structural alignment of BpGH16C with three other GH16 porphyranases: porphyranase A (PDB ID 3ILF) and porphyranase B from *Zobellia galactanivorans* (PDB ID 3JUU) [9] and another porphyranase from *B. plebeius* (PDB ID 4AWD) [16] shows that the Gly132 is replaced by either a Ser or a Thr, making this pocket too small to accommodate an additional methyl group on the C6-O (Figure 3D). Sequence alignment of β-porphyranases with BpGH16C and the two other characterized 6O-methyl-β-porphyranases: Patl_0824 of (*P. atlantica* T6c, this study) and AXE80_06940 (*W. fucanilytica*) [29] confirmed that the replacement of the Ser/Thr by Gly was correlated to the accommodation of the methyl group in the −1 sub-site of the catalytic site (Figure S1). The BpGH16C is the first crystallized 6-OMe-porphyranase that tolerates the 6-methyl group on the galactose ring. Indeed, there is an extra space in the active site at the end of the pocket that accommodates C6-OH, with its bottom formed by Gly132 (Figure 3E).

### Complete degradation of oligo-methyl-porphyrans

Following the same strategy reported by Robb and co-workers [18], we have incubated the methylated oligosaccharides with the BpS1_11 sulfatase, the BpGH29 α-L-galactosidasee and the predicted BpGH2C β-D-galactosidase. Purified methylated and non-methylated oligo-porphyrans were incubated with the BpS1_11 sulfatase and analyses of the products by ^1^H NMR showed a strong chemical shift (about 0.1 ppm) of the signal attributed to the proton at position 6 of the galactose located at the non-reducing end, demonstrating the removal of the sulfate ester group independently of the degree of methylation of the oligosaccharides (Figure S2).

Crystal structure of the BuS1_11 sulfatase from *B. uniformis* complexed with a di-saccharides [18] and the *B. plebeius* sulfatase BpS1_11 on its own (this study, PDB ID 7SNJ, 1.64 Å resolution) and complexed with a tetrasaccharide (this study, PDB ID 7SNO, 2.1 Å resolution) revealed that there is room in the active site to accommodate oligo-methyl-porphyrans (Figure S3). The porphyran sulfatase BpS1_11 belongs to the formyl-glycine dependent sulfatases, requiring maturation of a cysteine or a serine into the formyl-glycine residue as an active catalytic residue. A large N-terminal domain of BS1_11 contains active site residues and forms main part of the substrate binding site. A smaller C-terminal domain completes the substrate binding site.

To map the details of substrate binding, we determined the structure of H214N inactive mutant with L-α-6O-sulfate-Gal-(1→3)-D-β-Gal-(1→4)-L-α-6O-sulfate-Gal-(1→3)-D-a/β-Gal tetrasaccharide substrate (PDB ID 7SNO). The electron density map clearly showed the entire tetrasaccharide bound to BpS1_11 (H214N). Following the recently proposed nomenclature [33], the tetarasaccharide binds to the 0, +1, +2 and +3 positions (Figure S3). The 6S-L-Gal in position 0, which the sulfate is removed, has all three hydroxyl groups hydrogen bonded to the protein side chains: O2 to Arg271, O3 to Asp215 and Arg271, O3 to Asp345 and Arg347. The 6-sulfate group is facing Ser83 (formyl-glycine in mature enzyme) and one of its oxygens is a ligand to the Ca^2+^ ion. D-Gal in +1 position hydrogen bonds C6OH to NE2 of His133 and its O2 is bonded to the O2 of L-Gal (position 0) through a bridging water molecule. The +2 6S-L-Gal makes only one contact, between the 6-sulfate oxygen and His428. The +3 D-Gal extends far from the binding site and makes no contacts with the protein. However, in the crystal, this sugar hydrogen bonds from O1 and O4 as well as O5 within the ring to the guanidino group of Arg68 from a symmetry-related molecule. These interactions stabilize the +2 and +3 sugars in the crystal. Importantly, the 6-O group of +1 D-Gal is not tightly constraint by the enzyme and there is sufficient space for the additional methyl group present in the 6-O-Me-D-Gal (Figure S3).

We confirmed the exo-based cycle of porphyran depolymerization [18] using tetra-saccharides as starting substrate (Figure S4). After desulfation of the tetra-saccharides, BpGH29 exo-α-L-galactosidase was active on the methylated and non-methylated substrate demonstrating that the methyl group did not hinder the activity of the enzyme. Independently of Robb and co-workers [18], we have determined the structure of BpGH29 (PDB ID 7SNK). The superposition with their structure (PDB ID 7LJJ) shows root-mean-squares deviation of 0.6 Å, indicating that the structures that were crystallized under different conditions are virtually identical, and confirms that ordering of the loop 408-426 and small rearrangement of other surrounding loops result from the substrate binding to the active site.

Finally, we observed that the methylated or non-methylated D-galactose residue located at the non-reducing end of the trisaccharide was cleaved by the BpGH2C β-D-galactosidase (Figure S4). Altogether, the enzymology and crystallography experiments conducted in this work revealed the pathway leading to the degradation of the methylated fraction of porphyran. Our observations combined with previous investigations demonstrate that all the methylated and non-methylated fractions of porphyran are digested by the enzymes encoded by porphyran PUL of *B. plebeius*. Therefore, this PUL can catabolize autonomously the chemically complex porphyran, without the help of enzyme(s) external to this PUL.

### Phylogenetic distribution of the porphyran utilizing loci and gene organization

*B. plebeius* DSM 17135 was the first gut bacteria shown to carry the porphyran PUL. Recently, some strains of *B. fragilis*, *B. ovatus*, *B. thetaiotaomicron* and *B. xylanisolvens* were further shown to be able to grow on porphyran and to carry a >96% identical sequence to the *B. plebeius* porphyran PUL [13]. In order to investigate in more depth whether other human gut bacteria also carry the porphyran PUL, we used BLASTn to identify homologs of the entire coding sequences of *PUL-PorA* (17,980 bp), *PUL-PorB* (12,888 bp) and *PUL-PorC* (19,198 bp) of *B. plebeius* DSM 17135 among the 10,969 genomes of bacteria isolated from the human gut and catalogued by Almeida and co-workers [34]. We found that the PULs were present in the genomes of 22 *Phocaeicola*/*Bacteroides* strains (corresponding to at least 8 different identified species (Table S1). All these strains came from isolates from the gut microbiome of East Asian individuals, and the same species/strains isolated from other human populations worldwide had no homologs of porphyran PULs. The identified PULs were present in different species, including *B. dorei*, *B. eggerthii*, *B. ovatus*, *B. plebeius*, *B. stercoris*, *B. uniformis*, *B. vulgatus* and *B. xylanisolvens,* but were absent in other common *Bacteroides* sister species from the human gut such as *B. intestinalis*.

While the *PUL-PorB* organization as observed in *B. plebieus* DSM 17135 was conserved in all the other strains we have identified, the gene organization of *PUL-PorA* was only conserved in 17/22 strains (Figure 4). In five other strains (*B. plebeius* strain AM09-36, *B. stercoris* strain AF05-4, *B. uniformis* AF21-53, *B. uniformis* AF26-10BH and *B. uniformis* strain AM43-9), the homolog of the *B. plebieus* β-agarase (GH86, Bacple_01694) appeared to have recombined with a DNA fragment composed of two genes predicted to encode metabolic enzymes: 2-dehydro-3-deoxygluconokinase and 2-dehydro-3-deoxyphosphogluconate aldolase (Figure S5). For these last four strains, a deletion of the genes encoding the GH50 (Bacple_01683) and GH105|GH154 (Bacple_01684) located in *PUL-PorC* was also observed, highlighting the remodeling of the PUL by insertion and deletion of DNA segments (Figure S5). Note that parts of *PUL-PorA* and *PUL-PorC* were incompletely sequenced in *B. plebeius* strain AM09-36 and *B. eggerthii* strain AM42-16.

**Figure 4:**
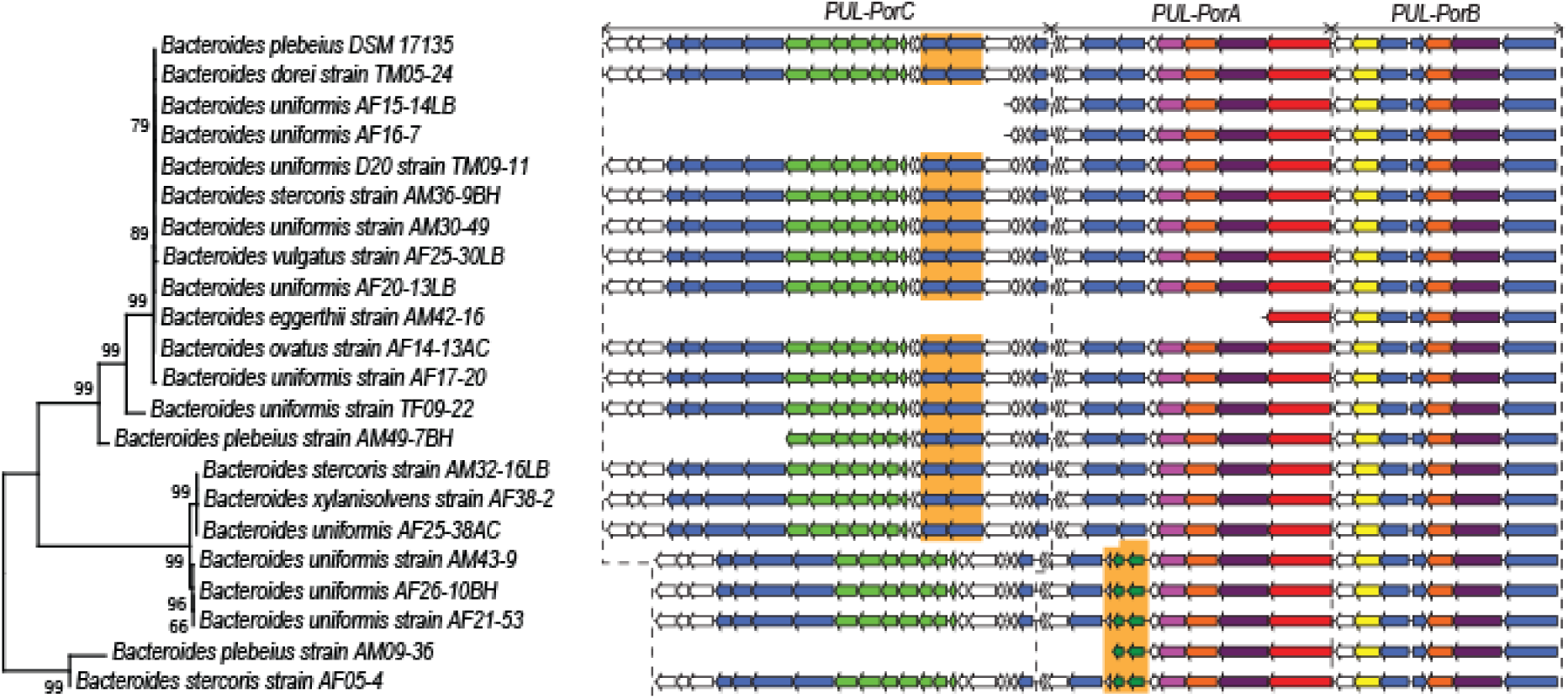
Gene organization of the porphyran degradation system of *B. plebieus* compared with other identified human gut bacteria. Phylogenetic tree was calculated using concaneted gene sequences of PUL-PorB. PUL organizations of each strains were indicated based on the available sequencing data.

To explore further the phylogenetic diversity of *PUL-PorB*, we scanned the whole catalog of Almeida et al [34], including not only the previously mentioned 10,969 gut bacterial isolates, but also 278,263 gut bacterial metagenomes-assembled genomes (MAGs) coming from 13,587 worldwide individuals, which we completed with 29,082 MAGs coming from 842 individuals from under-represented Asian countries (Japan, India, Korea) [35]. We used BLASTp to identify homologs of the six known proteins from *PUL-PorB* and found overall 242 strains having a full *PUL-PorB* sequence, which were reduced to 130 non-redundant ones after removing strains from the same bacterial species having 100% identity (Table S2). This analysis revealed four additional *Phocaeicola*/*Bacteroides* species that acquired the porphyran PUL (*B. coprocola*, *B. stercorirosoris*, *B. caccae, B. fragilis*) but also two new genera: *Tyzzerella sp*. and *Holdemanella sp*. which belong to the Bacillota phylum.

### Genetic diversity of *PUL-PorB*

The neighbor-joining tree of these 130 non-redundant strains, calculated with the six concatenated protein sequences of *PUL-PorB* (Figure 5A), revealed four clades with high bootstrap values, allowing us to classify the strains in four groups: GI, GII, GIII and GIIIrec.

**Figure 5:**
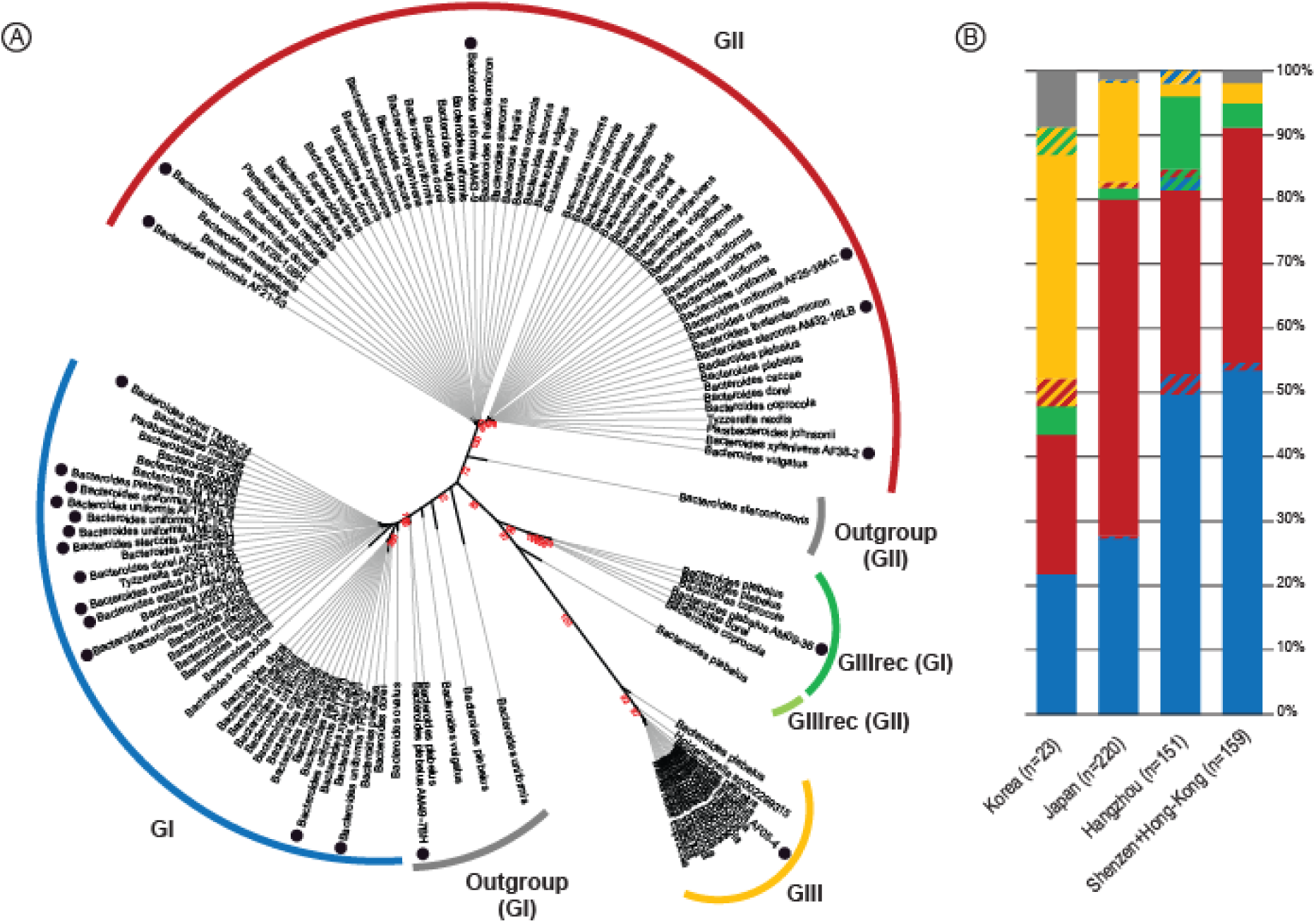
**A)** Phylogenetic tree calculated with the concatenated protein sequences of *PUL-PorB* recorded in non-redundant human gut bacteria including isolated strains (●). The observed clades were used to create groups of homologous porphyran PUL *PUL-PorB* (GI, GII, GIII and GIIIrec) and divergent *PUL-PorB* (Outgroup). **B)** Distribution of the different group of the *PUL-PorB* in human gut metagenome assembly of South east and North East Chinese, Korea and Japanese populations.

The percentage of identity of each *PUL-PorB* protein was compared with the corresponding sequence of *B. plebeius* DSM 17135 (Table S2). Phylogenetic trees were calculated with each protein dataset (not concatenated) to validate their classification. In the group GI, which includes the *B. plebeius* DSM 17135 reference, most protein sequences presented 99.5-100% identity with the reference. Similarly, for the members of the group GII, the protein sequences were 99-100% identical to each other, but presented 96-100% identity with the *B. plebeius* reference. At the root of the GI clade (Outgroup GI), the concatenated sequences of *PUL-PorB* of 5 strains presented lower identity (98-100%), reflecting some recombination events. For example, the isolated *B. plebeius* strain AM49-7BH suggested that its divergence with GI members resulted from a recombination event that occurred between SusD, SusC and GH2 genes from GI and Sulf and GH29 genes more related to GII. The recombination site was identified in the gene coding for GH16 (Figure S6).

The clade GIII encompasses members that presented 92-98% identity with the *B. plebeius* reference DSM 17135. The *PUL-PorB* sequence of the GIIIrec members seems to result from a recombination event between GI and GIII (Table S2), since the SusD (bacple_01704), SusC (bacple_01705) and GH2 (bacple_01706) sequences presented 99-100% identity with GI, while the S1_11 sulfatase (bacple_01701) and GH29 (bacple_01702) were nearly identical to GIII sequences. Again, the recombination site was identified in the gene coding for GH16 (Figure S6).

### Geographic distribution of *PUL-PorB* in assembled metagenomes

We then explored the worldwide geographic prevalence of the porphyran degradation system in diverse human gut microbiota by probing the six *PUL-PorB* genes against assembled human gut metagenomic datasets (Table S3). We first scanned 20 manually downloaded projects covering a total of 3,556 biosamples from 10 countries for which gut metagenomes were obtained and assembled. Only fully sequenced genes were retrieved, and individuals were considered positive if they carried at least one fully sequenced gene of *PUL-PorB* out of the 6 probed ones. We found that in East Asia, 29% of individuals were positive (585 out of the 2,016 tested) – corresponding to 33% of the Chinese individuals, 25% of the Japanese individuals and 26% of the Korean individuals, while we found only six positive individuals among the 1237 North American gut metagenomes (0.5% of the individuals) and none among the other geographic areas, including 117 individuals from South Asia (India, Bangladesh), 36 from South America (Peru), 139 from Africa (Tanzania, Madagascar) and 11 from Europe (Italy) (Table S3).

We then conducted a similar analysis with the catalog of 278,263 MAGs [34] coming from 13,587 individuals living in 30 different countries (13 from Europe, 7 from Asia, 6 from America, 2 from Oceania and 2 from Africa). We found that 19% of individuals from Japan (6 out of 31) were positive (similar to our previous analysis above), while only 4% of Chinese (117 out of 3134) were positive. All other countries had no hits or had prevalence lower than 1% (from higher to lower prevalence: 1/110 positive individual in Mongolia, 1/238 in Fiji, 2/759 in Sweden, 7/3036 in USA, 2/954 in Israel, and 1/1083 in Denmark). Investigating why the prevalence is so low in China in this catalog as compared to our previous analysis based on the original studies (4% versus 36%), made us realize that the filters applied by Almeida and co-workers [34] were very stringent. Indeed, to have a high-quality catalog, they removed the contigs that had less than 50% completeness and a quality score less than 50. Consequently, the number of metagenomes per individual are much lower (about ten times) than in the original projects. We can thus conclude that the dataset compiled by Almeida et al [34] is not suited to obtain the PUL prevalence values.

Overall, consistently with previous investigations, the occurrence of algal polysaccharides degradation systems seems to be restricted to East Asia (Japan, China, Korea) and not be present in other Asian countries (Kazakhstan, Mongolia, India, Bangladesh, Singapore) or elsewhere in the world.

The 591 positive individuals from the original 20 metagenome projects (not the filtered catalog) were then characterized as belonging to GI, GII, GIII and/or GIIIrec based on their *PUL-PorB* gene sequence (Table S4). For 95.4% of the 585 positive East Asian individuals, the genes detected were attributed to a single group; however, few individuals (4.6%) presented *PUL-PorB* genes that belonged to two different groups. In order to obtain large sets, the individuals were grouped by geographic location (Figure 5B). We distinguished populations living in North-East China (Hangzhou), South-East China (Shenzen and Hong-Kong), Korea and Japan. In both set of Chinese individuals, the group GI was present in more than half of the population (57% and 55%, respectively) followed by the group GII (33% and 39%, respectively). In contrast, in Japan, GII was the major group, found in twice higher proportion than GI (55% and 28%, respectively). GIII and GIIIrec were minor groups in these populations, with GIII found in higher proportion in Japan (16%) and GIIIrec in higher proportion in China (14%, 4% and 0% in Hangzhou, Shenzen and Hong-Kong, respectively). In Korea, even though much fewer individuals are analyzed (n=23), the picture seems to be different from both China and Japan, with GIII being the major group (43%) and GI, GII being at similar frequencies (22% and 26%, respectively).

### Analysis of short read datasets

The analyses of the assembled metagenomes revealed that 20% to 30% of individuals carry the *PUL-PorB* in East Asia (Japan, China, Korea), but that this PUL is mostly absent in individuals from other parts of the world (<1%). In addition, 95.2% of the individuals carry only one version – one group – of the *PUL-PorB*. However, these observations may include some bias inherent to the bioinformatics processes involved in the building up of the contigs. Therefore, to analyze the data devoid of such biases, we have selected a 54 nucleotides sequence allowing to probe the different *PUL-PorB* groups directly on raw sequencing data (short reads). The probe was part of the Bacple_01703 (GH16) gene encoding the methyl-porphyranase, which presented distinct mutations specific to the GI, GII, GIII and GIIIrec groups (Figure S7). This method allowed us to explore new metagenomic datasets which were not assembled, thus expending the number of sampling sites, as well as the number individuals tested, with now a total of 4,617 individuals (Table S3).

As observed previously, we confirmed by this complementary approach that *PUL-PorB* is only found in East Asian coastal populations (China, Japan, Korea), but not in America, Africa or other parts of Asia (Mongolia, India, Malaysia), where the prevalence is null to extremely low (3% at most in Mongolia) (Table S3, Figure 6). However, we noted that the prevalence in Kuala Lumpur was found to be 16% (but 1% in rural Malaysia), likely reflecting migration fluxes from China. These results allowed us to confirm indirectly the specificity of our probe showing that it does not result in false positives. Across Asia, the prevalences obtained are quite higher than the ones obtained with the assembled datasets (50% in China, 72% in Japan and 89% in Korea), with also a certain variance across datasets in Japan (15-94%) and in China (20-70%).

**Figure 6:**
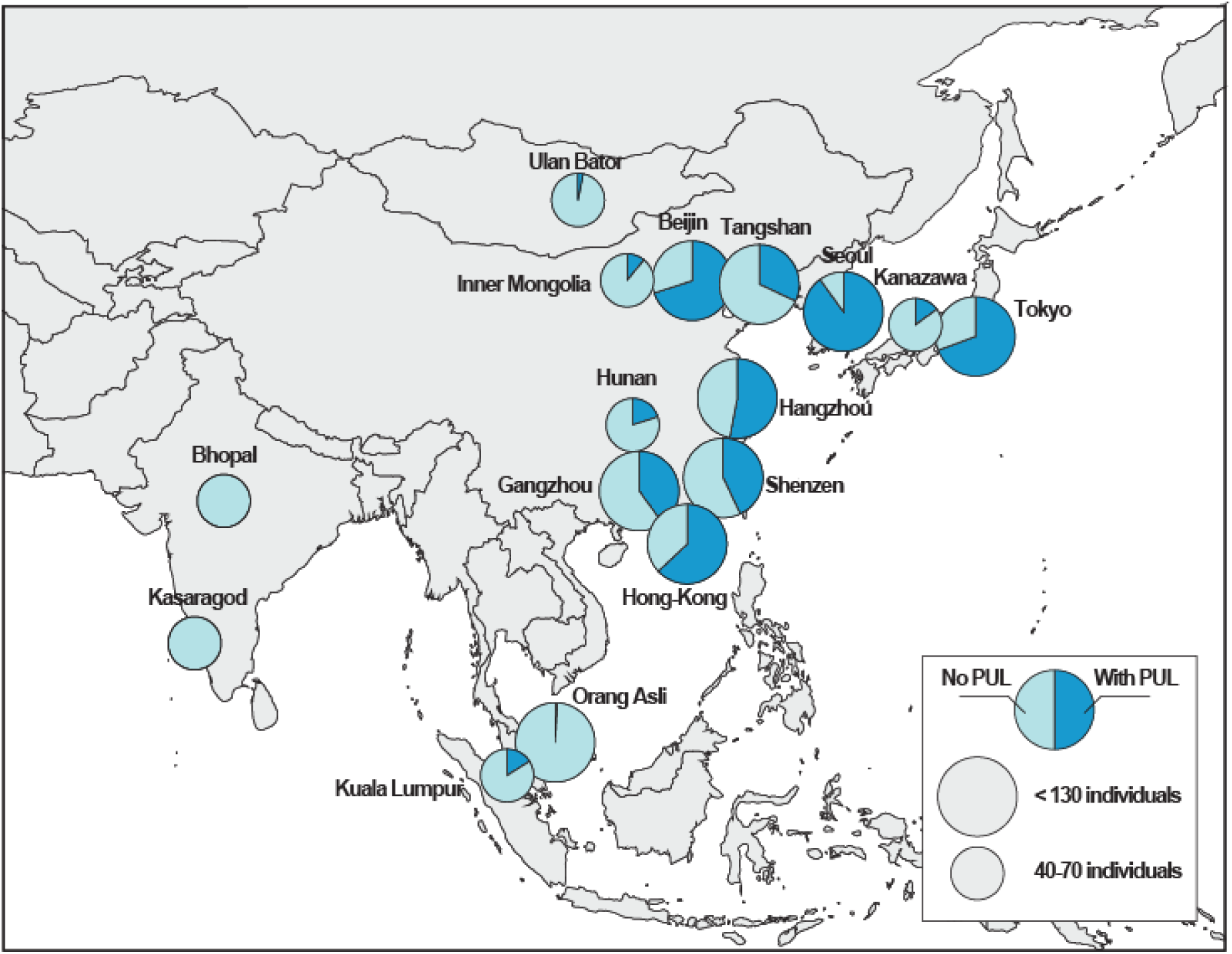
Proportion of individuals with short reads mapping to the 54-nucleotides probes specific to the *PUL-PorB* in Asia (PUL-positive individuals), grouped by city. Some geographic coordinates have been adjusted so that each point is visible.

Because of the variability in sequencing depths across datasets, we first tested whether the prevalence of *PUL-PorB* could be influenced by coverage (with the idea that more false negatives could be observed in datasets with too little coverage). However, we observed that there is no significant effect of coverage on *PUL-PorB* prevalence (Pearson correlation coefficient r=0.14, p-val=0.499).

We further found that there is no effect of age (chi-squared test, p-val=0.086), even though metadata on age was available only for two Chinese projects (PRJNA422434 and PRJNA356225); we did not find a significant effect of sex either Shenzen (PRJNA422434: chi-squared test, p-val=0.158).

We then estimated the abundance of bacteria carrying *PUL-PorB* by looking at the proportion of reads blasting against it in positive individual. We found that it varies between 10^-11^ to 10^-7^ between individuals (with absolute counts being between 1 and 3099), with a median of 10^-9^. Interestingly, Chinese have 3-4 times lower abundances of *PUL-PorB* than Koreans or Japanese (0.7×10-9 versus 2.6×10-9 and 3.2×10-9, respectively, ANOVA p-val<2.2e-16, Figure S8).

Because of the variability in sequencing depths across individuals, we then excluded individuals with too few (less than 10) reads to characterize their *PUL-PorB* genetic diversity. From 2138 positive individuals, we thus retained 1171 individuals having at least 10 reads. The histograms presented in Figure S9 show the number of hits for these individuals and their attribution to the GI, GII, GIII and GIIIrec groups. We found that 60% of individuals carried short reads attributed to a single group, and an additional 21% of individuals had a dominant group, defined as having a prevalence of the major group above 80%. Thus, we estimated that overall, 81% of individuals carried only one or a dominant group. Quite notably, more individuals carried a single or dominant group in China and Japan (84%) as compared to Korea (53%), where half of the individuals carried co-dominant groups. To verify that this is not due to the about five times higher coverage in Korean samples, we looked at the 52 Korean individuals with less than 100 reads (corresponding to individuals with a median number of positive reads of 42, thus quite similar to the median in Japanese of 47 and that in Chinese of 29), and we have found a similarly low number of individuals with one or a dominant group (39%) in this subset.

Looking at the prevalence of individuals being positive for each group, we confirmed that GI were predominant in Chinese populations (52% of individuals, versus 33% in Koreans and 32% in Japanese), while GII was most common in Japanese populations (62%, versus 53% in Koreans and 46% in Chinese) (Figure 7). Finally, the GIII and GIIIrec groups were more prevalent in Koreans (60%) as compared to Japanese (25%) and Chinese (14%).

**Figure 7:**
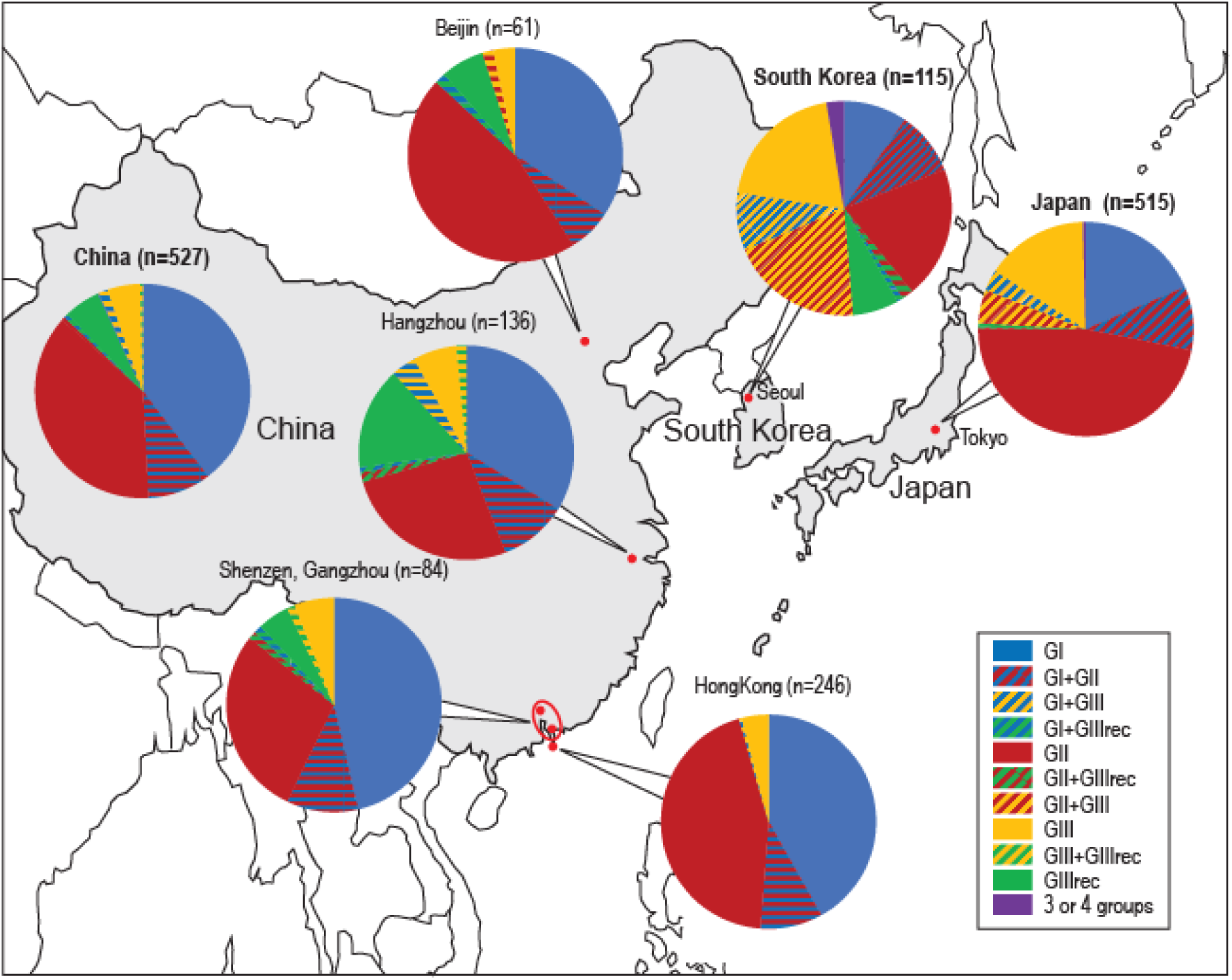
Distribution of the various porphyran PUL among Chinese, Japanese and Korean populations. The pie charts derive from the histogram presented in Figure S9 obtained from the analyses of short read sequencing of Chinese, Japanese and Korean metagenomics data.

Including only categories with large enough numbers (more than 10 individuals overall), we found that the six populations were overall significantly different (X-squared= 324.41, p-value<2.2×10^-16^) (Figure 7). This difference was mostly driven by significant differences between countries (all comparisons between countries: q-value<1e-04), as well as between Hong-Kong and Hangzhou, and Hong-Kong and Shenzen/Gangzhou (q-value=0.027 and 0.044, respectively). We then tested separately which group was significantly different in prevalence between populations and found that all were significantly different (proportion test, q-value<5e-06), but GIII and GIIIrec presented the stronger difference (proportion test, q-value=1.1×10^-15^).

## Discussion

Biochemical and crystallographic investigations showed that the porphyran PUL encodes tailored enzymes dedicated to the complete degradation of the polysaccharide, including the methylated fraction, giving galactopyranoside and 6-O-methyl-galactopyranoside as end-products. We showed that the 6-O-methyl-porphyranase (BpGH16C) presents an active site able to accommodate the methyl group, in contrast with the active sites of the previously investigated β-porphyranases. Similarly, the exo-6O-L-galactose porphyran sulfatase Bp S1_11 can accommodate methylated oligosaccharides in its active site. The demethylation enzyme of 6-O-methyl-galactopyranoside was located in agarose PUL of several aerobic bacteria belonging to the Gammaproteobacteria, Bacteroidetes and Planctomycetes phyla. In these bacteria, the reaction was obtained with specific monooxygenase enzyme system, including ferredoxin, ferredoxin reductase and P450 monooxygenase [36]. However, demethylation of galactose residue in anaerobic bacteria has not been elucidated yet and no encoding gene surrounding the porphyran utilization loci could be suspected to be involved in the demethylation pathway.

Thanks to a large catalog of assembled gut bacterial genomes [34, 35] including both isolated strains and metagenome-assembled genomes (MAGs), we were able to interrogate a large number of sequences (about 300,000) and show that more than a hundred different strains carry the porphyran PUL, corresponding to 12 *Bacteroides* species as well as two other genera from the Bacillota phylum (e.g. *Tyzzerella sp*. and *Holdemanella sp*.). The gene organization of the porphyran PUL is furthermore quite conserved, with a minority of rearrangements events (i.e. loss or duplication of genes). This likely indicates a unique common ancestor of these PUL from the environment, and then the occurrence of multiple transfers of this system across closely related gut bacterial species by horizontal gene transfer. Importantly, within a given bacterial species, strains with or without the porphyran PUL coexist. Consequently, the taxonomic composition of an individual is not a good predictor of its metabolic capacity.

As already shown, these transfers seem to happen preferentially between phylogenetically closely related bacteria [37, 38], as the strains carrying the PUL were mostly found in the genera *Bacteroides*. While it would be interesting to know more about the timing of the original transfer of the porphyran PUL from the marine to the gut bacteria, the environmental bacteria from which the transfer occurred have not been identified (and does not seem to be present in current databases). Indeed, searching marine bacterial genomes and metagenomic data (e.g. Tara Ocean) failed to identify a porphyran PUL similar or distantly related to those observed in the human gut. Therefore, it is presently not possible to establish when and how the porphyran PUL has evolved from its marine ancestor.

Because harvesting, drying and cooking *Porphyra sp*. were conserved practices in numerous populations living not only along the Pacific but also in Europe, and because these ancestral traditions seem to follow the dispersion line of *Homo sapiens* from Asia to the South of the American continent, we wondered if the porphyran degradation system may be present in a larger set of populations than the Chinese and Japanese where it was previously identified. We again took advantage of the large catalog of MAGs from Almeida et al [34], which we completed with another dataset from Japanese, Korean and Indian populations [35] to evaluate the geographic prevalence of the porphyran PUL degradation system. This allowed us to precisely define the distribution area of this system, which appears to be limited to coastal East Asian populations (Japan, China, Korea), as well as some continental Chinese populations and few individuals from urban Malaysia (potentially due to Chinese immigration). However, it is not found in neighboring countries such as Mongolia, rural Malaysia or India (nor Peru or Italy), suggesting, as expected, that strains carrying the PUL are not selectively retained in populations that do not include algae in their diet. Further studies including coastal countries in South America would be interesting to definitively assess whether the PUL is present there.

While the catalog from Almeida and co-workers [34], a compilation of many datasets covering more than 13,000 individuals, was very useful to be able to scan the gut microbiome at a large geographic scale, the prevalence obtained on that catalog seems to be biased downwards (4% in China versus 33% on the same manually downloaded datasets). Indeed, for the assembled genomes to be of high quality, stringent quality filters were applied to the catalog.

We thus could not exploit that catalog to obtain reliable prevalence data, which we instead estimated in the original projects. This analysis revealed a prevalence of the porphyran PUL of about 30% in East Asia. This number is itself quite different from the one obtained when analyzing directly the short-read data (unassembled), where we observed a prevalence of 50% in China, 72% in Japan and 89% in Korea. The most plausible explanation is that this genomic region is quite hard to assemble, because it is found on different bacterial backgrounds. Bacterial long-reads data, which start to emerge to look at bacterial structural variation [39], might help us better understand this discrepancy between the short-reads data and the assembled datasets. In any case, there seems to be a higher proportion of individuals carrying the PUL in Korea and Japan than in China, as well as three times more reads blasting against the PUL (reflecting the abundance of PUL-positive bacterial strains) in positive individuals from Korea and Japan compared to China. This might indicate that PUL-positive bacterial strains thrive more in the gut of Koreans and Japanese, probably because of a higher amount of porphyran in their diet, and thus, their gut. Some discrepancies remain to be explained, though, notably the large difference between the prevalence of the porphyran PUL in two datasets from Japan, one from Kanazawa (15%) and one from Tokyo (76%).

While we did not detect a significant effect of age on the prevalence of PUL-carrying bacteria when merging the data from the only two datasets with age information, we still identified a trend toward a lower prevalence in individuals over 40 years old, which was significant in the dataset from Shenzen alone (PRJNA422434, p-val=0.023). This effect deserves to be further investigated to see whether it is replicated in other cohorts and to test whether this relate to different dietary habits between younger and older persons.

Investigation of the genetic diversity within *PUL-PorB* showed a certain amount of genetic structure across individuals which allowed us to define three clear clusters of sequences, as well as a recombinant group between two of them: GI, GII, GIII and GIIIrec. These groups are not associated with particular bacterial species, but rather are found in different proportions across countries. Notably, GI was found in higher frequency in China, GII in Japan and GIII in Korea. Such significant differences in frequency between countries are what we expect to observe in a model where all populations are not entirely inter-connected. Conversely, the higher similarity between different populations in China might indicate that these populations are more connected in term of migration, and thus in term of exchanges of bacteria between individuals by horizontal transmission. However, even though differences in frequency are observed, all *PUL-PorB* groups (GI, GII, GIII, GIIIrec) are observed in all populations, showing that these strains do circulate at a quite large geographical scale. In general, this system is a very unique one, where we can track the migration of individuals through the diffusion of the PUL. It could be interesting to use it as a model to estimate the rate of horizontal transmission of gut bacteria between individuals at greater or lesser geographic and cultural distances.

Interestingly, analyses of both the MAGs and the short reads revealed that most individuals (80%) carry a single PUL variant, or a dominating one (even though less so in Korea – 53% of individuals), giving the impression of an apparent haploidy of the PUL. This observation has also been made for 80% of gut bacterial species, where one strain is dominant [40], so this might be a property of how bacteria establish and evolve in this peculiar environment.

In conclusion, the porphyran PUL encodes a portfolio of enzymes catalyzing the complete degradation of the methylated and unmethylated fractions of the polysaccharide without the help of external enzymes. While the architecture of the human gut microbiota across populations has mostly been studied and functionally interpreted at the genera or species level, we see here with the example of the porphyran degradation system that the important functional information is at the strain, or even the genetic level. Indeed, the PUL system is carried by various species and strains of the *Bacteroides* genus, and the GI/GII/GIII groups are not associated with particular strains, so the inference of these groups based on taxonomy would not be possible. In this study, we demonstrated that geographically distant human populations (at the scale of East Asia) present different prevalence of PUL-carrying bacteria, and among positive individuals, these countries have similar groups, but at significantly different frequencies. Overall, the current structure of the investigated populations is likely the result of lateral transfer, recombination, as well as migration events, which reflect an evolution history of the gut microbiota and therefore, of its host.

## Material and methods

### Purification of porphyran

6 g of dried *Porphyra columbina* were suspended in 120 ml de distilled H_2_O and autoclaved for 30 min at 120°C. After 14 h at room temperature, the suspension was centrifuged at 8000 rpm during 30 min at 4°C. The supernatant was added to 120 ml of pure ethanol (50% v/v EtOH final concentration) and the solution was maintained at 4°C for 2h. After centrifugation (8000 rpm, 30 min, 4°C), the porphyran, present in the supernatant, was precipitated by addition of 130 ml of pure ethanol (67% v/v EtOH final concentration). After centrifugation the porphyran pellet was dissolved in distilled water, and dialysis for 3 days against distilled water using dialysis membrane with a cut-off 1000 Da (pre-wetted Spectra/Por® 6 dialysis tubing – Sprectrumlabs). The polysaccharide was lyophilized and the purity was verified by ^1^H-NMR. The yield of purification was about 15-20 % w/w dried algae.

### Heterologous expression of *Bacteroides plebeius* DSM 17135 PUL-PorB enzymes

The genes encoding the predicted glycoside hydrolases (BpGH29, Bacple_01702; BpGH16C, Bacple_01703; BpGH2C, Bacple_01706) and sulfatase (BpS1_11, Bacple_01701) from *B. plebeius* DSM 17135 were cloned using genomic DNA as template in the pET-28 or pFO4 expression plasmid [41] without their signal peptides identified with SignalP [42]. The expression strains harboring the recombinant expression plasmids were grown in Luria Bertani (LB) medium supplemented with 50 µg/ml kanamycin (pET28a plasmid) or 100 µg/ml ampicillin (pFO4 plasmid) until the OD_600nm_ reached 0.6 in a shaking incubator working at 180 rpm and 37°C. After the addition of isopropyl-β-D-thiogalactopyranoside (IPTG), the temperature was cooled down at 20°C and maintained overnight.

Cultures (200 mL) were centrifuged at 6000 g for 15 min and the bacterial pellet was suspended in 10 ml of buffer A (20 mM Tris pH 8, 500 mM NaCl, 20 mM Imidazole). The cells were lysed using a cell disrupter (Constant system Ltd). Insoluble fractions were removed by centrifugation at 20000 g during 30 min at 4°C and the supernatant was loaded on a 1 mL HisTrap^TM^ HP column (GE Healthcare) connected to a NGC chromatography system (BioRad). The proteins were eluted with a imidazole gradient from 20 to 300 mM gradient of imidazole. Pure enzymes were obtained after a size exclusion chromatography using a HiLoad 16/600 Superdex 75 pg column in buffer B (10 mM Tris-HCl pH 8, 50 mM NaCl).

#### Enzymatic assays

Enzymatic degradations were monitored by analytical gel permeation chromatography using Superdex S200 10/300 and Superdex peptide 10/300 (GE Healthcare) columns mounted in series and connected to a high-performance liquid chromatography (HPLC) Ultimate 3000 system (Thermo Fisher). The injection volume was 20 µL and the elution was performed at 0.4 mL.min-1 in 0.1 M NaCl. Oligosaccharides were detected by differential refractometry (Iota 2 differential refractive index detector, Precision Instruments).

The oligosaccharide products were purified by semi-preparative gel permeation chromatography using three HiLoad® 26/600 Superdex® 30 pg (GE Healthcare) columns mounted in series and connected to a semi-preparative size-exclusion chromatography system which consisted of a Knauer pump (pump model 100), a refractive detector (iota2 Precision instrument) and a fraction collector (Foxy R1) mounted in series. The elution was conducted at a flow rate of 1.2 mL.min^-1^ at room temperature using 100 mM (NH_4_)_2_CO_3_ as eluent. The collected fractions were freeze-dried prior NMR and mass spectrometry analyses.

Samples were exchanged twice with D_2_O and were transferred to a 5 mm NMR tube. ^1^H NMR spectra were recorded at 323 K using an Advance III 400 MHz spectrometer (Bruker). Chemical shifts are expressed in ppm in reference to water. The HOD signal was not suppressed.

#### NMR

^1^H NMR spectra were recorded with a Bruker Avance 400 spectrometer operating at a frequency of 400.13 MHz. Samples were solubilized in D_2_O at a temperature of 293 K for the oligosaccharides and 353 K for the polysaccharide. Residual signal of the solvent was used as internal standard: HOD at 4.85 ppm at 293 K and 4.35 ppm at 343 K. Proton spectra were recorded with a 4006 Hz spectral width, 32,768 data points, 4.089 s acquisition times, 0.1 s relaxation delays and 16 scans.

#### Crystallization, data collection and structure solutions

The homogenous fractions of proteins obtained from gel permeation chromatography were concentrated and crystallization experiments were attempted. Initial crystals were obtained by screening against wide range of commercial screens and the hits were optimized by hanging drop diffusion method. The drop containing 1 μl of protein and 1 μl of reservoir solution was incubated over 1 ml of reservoir solution and crystal growth was monitored regularly. The conditions of the best diffracting crystals of the three proteins were listed: BpS1_11 (Bacple_01701) was crystallized at 23 mg/ml in 40% Peg200, 0.1 M Sodium Citrate buffer pH 5.5 and 30% MPD; BpGH29 (Bacple_01702) was crystallized at 18 mg/ml in 20% Peg 8K and 0.1 M KH2PO4; BpGH16C (Bacple_01703) was crystallized at 34 mg/ml in 16% Peg 8K, 0.1 M Hepes pH 7.5 and 0.2M Calcium acetate. For diffraction experiments, the crystals were cryo protected in reservoir solution containing 20% MPD (BpS1_11), 25% glycerol (BpGH29), 30% ethylene glycol (BpGH16C) and flash frozen in liquid nitrogen. The diffraction data of all the crystals were collected at the 08ID beamline at the Canadian Light Source [43].

The X-ray diffraction data were processed using XDS [44]. The structures were solved by molecular replacement using the program Phaser in Phenix package [45]. For BpS1_11, the structure solution was obtained by molecular replacement using the search model an arylsulfatase from *Pseudomonas aeruginosa* (PDB ID 1HDH) [46]. For BpGH29, the structure solution was obtained using the α-L-fucosidase from *Fusarium graminearum* as a search model (PDB ID 4NI3) [47]. BpGH16C was solved using porphyranase B from *Zobellia galactanivorans* (PDB ID 3JUU) [9] as a model. All the structures were refined with the Phenix software [48] and manual rebuilding and solvent placement was conducted with the COOT program [49]. The stereochemistry of all the models were validated with MolProbity [50].

To obtain the complex structure, putative active site mutants of BpS1_11 (BpS1_11(H214N)) and BpGH16C (BpGH16C(E145L)) were made using the Quickchange site-directed mutagenesis protocol and using KOD polymerase. Briefly, the plasmid containing the gene of interest was amplified with the primer pairs carrying the mutation using KOD polymerase. After PCR, the template plasmid was digested with 1 µl of DpnI enzyme for an hour at 37°C. Five µl of PCR product was then transformed into 50µl of chemically competent *E. coli* DH5α cells. The clones carrying the desired mutation were confirmed by sequencing and proceeded for crystallization experiments.

Based on the structural comparison with other sulfatases, the His 214 residue in BpS1_11 was mutated to Asn (H214N). The mutant protein BpS1_11(H214N) was expressed and purified following the same protocol used for wild type enzyme. Purified Bacple_01701(H214N) was crystallized from solution containing 40% PEG 200, 0.1M sodium citrate buffer pH 5.5 and 30% MPD. The putative active site mutant of BpGH16C(E145L) was purified following the same protocol used for wild type enzyme. BpGH16C(E145L) was crystallized from solution containing 20% PEG 8K, 0.1 M Hepes pH 7.5 and 0.2 M calcium acetate. BpS1_11(H214N) and BpGH16C(E145L) crystals were soaked in the tetrasaccharide solution for an hour before the diffraction experiments. The soaked crystals were flash frozen in liquid nitrogen and diffraction data was collected at the 08ID beamline in the Canadian light source [43]. The diffraction data was processed with XDS [44]. One round of rigid body refinement was carried out using Phenix refinement program [48]. Manual rebuilding and substrate placement were done using coot [49]. The geometry was validated using MolProbity program [50].

#### Human gut metagenomes analysis

More than 10,000 non redundant genomes of bacteria isolated from the human gut were downloaded from the unified catalog of Almeida et al. [34]. Bacterial strains harboring the porphyran PUL were identified after BLASTn of PUL-*PorA*,-*B* and -*C* sequences against the set of genome. Genes of the identified porphyran PUL variant were retrieved from bacterial genome repositories. Genes and concatenated genes were aligned using Muscle and distant tree were built using MEGA6 [51] allowing to distinguish GI, GII, GIII and GIIIrec groups.

Assembled bacterial isolates and metagenomes (Table 2) were downloaded and analyzed locally. Six genes of the porphyran *PUL-PorB* were BLASTn and aligned with Muscle using MEGA6. Only coding genes with full length were kept and compared with those of the reference bacteria. We kept only hits with >90% identity and 100% coverage. Distant trees were built allowing to group the genes in GI, GII, GIII or GIIIrec with bacterial isolate. To identify GIIIrec in contig from metagenomes, we required to observe genes matching both GI and GIII on the same contig. When only part of the genes was available, we considered the major group (i.e., GI or GIII) as the most probable one, creating a slight bias downward for GIIIrec. When multiple contigs in the same individual were found to be positive and they corresponded to different groups, we considered the individual as multi-groups (e.g., GI+GII). When we could not assign a contig to one of the 4 groups defined earlier, we considered the individual to be unresolved (outgroup).

Analyses of short reads data set (Table S3) were conducted using 54 nt probes (Figure S7). The probes were used to search at Sequence Read Archive (SRA) data at NCBI.

