## Supplementary_tables_figures for "The porphyran degradation system is complete, phylogenetically and geographically diverse across the gut microbiota of East Asian populations"

**Table S1**: List of the bacterial isolated strains which assembled genome contained genes of the porphyran degradation system. *Bacteroides* and *Phocaeicola* are basonyms. The name of the strains were reported as reported in databank.

| GenBank assembly | Assembly level | Biosamples | Strains |
|---|---|---|---|
| GCA_003466465.1 | Scaffold | SAMN09736981 | *Phocaeicola dorei* TM05-24 |
| GCA_003467285.1 | Scaffold | SAMN09736544 | *Bacteroides eggerthii* AM42-16 |
| GCA_003465275.1 | Scaffold | SAMN09734245 | *Bacteroides ovatus* AF14-13AC |
| GCA_000187895.1 | Scaffold | SAMN00000028 | *Bacteroides plebeius* DSM 17135 |
| GCA_003472665.1 | Scaffold | SAMN09734757 | *Phocaeicola plebeius* AM09-36 |
| GCA_003463985.1 | Scaffold | SAMN09736616 | *Phocaeicola plebeius* AM49-7BH |
| GCA_003465625.1 | Scaffold | SAMN09734188 | *Bacteroides stercoris* AF05-4 |
| GCA_003468685.1 | Scaffold | SAMN09736438 | *Bacteroides stercoris* AM32-16LB |
| GCA_003468355.1 | Scaffold | SAMN09736487 | *Bacteroides stercoris* AM36-9BH |
| GCA_003460085.1 | Scaffold | SAMN09734330 | *Bacteroides uniformis* AF17-20 |
| GCA_003459325.1 | Scaffold | SAMN09734434 | *Bacteroides uniformis* AF21-53 |
| GCA_003468865.1 | Scaffold | SAMN09736420 | *Bacteroides uniformis* AM30-49 |
| GCA_003466885.1 | Scaffold | SAMN09736574 | *Bacteroides uniformis* AM43-9 |
| GCA_003437605.1 | Scaffold | SAMN09736925 | *Bacteroides uniformis* TF09-22 |
| GCA_003458385.1 | Scaffold | SAMN09734500 | *Phocaeicola vulgatus* AF25-30LB |
| GCA_003474645.1 | Scaffold | SAMN09734674 | *Bacteroides xylanisolvens* AF38-2 |
| GCA_003603275.1 | Scaffold | SAMN09734277 | *Bacteroides uniformis*. AF15-14LB |
| GCA_003603315.1 | Scaffold | SAMN09734319 | *Bacteroides uniformis*. AF16-7 |
| GCA_003603065.1 | Scaffold | SAMN09734409 | *Bacteroides uniformis* AF20-13LB |
| GCA_003602995.1 | Scaffold | SAMN09734502 | *Bacteroides uniformis* AF25-38AC |
| GCA_003602945.1 | Scaffold | SAMN09734509 | *Bacteroides uniformis* AF26-10BH |
| GCA_003436255.1 | Scaffold | SAMN09737012 | *Bacteroides uniformis* D20 TM09-11 |

**Table S2**: List of the 130 non-redundant bacterial strains containing the complete *PUL-PorB*. The percentage of identity of the protein sequences of the *PUL-PorB* were compared with those of the reference *B. plebeius* DSM 17135. The *PUL-PorB* group of each individuals / isolate was determined based on the phylogenetic tree calculated on concatenated

| Ref catalog | Strain | MAG/ISO | PUL-PorB Group | Bacple_01701 | Bacple_01702 | Bacple_01703 | Bacple_01704 | Bacple_01705 | Bacple_01706 |
|---|---|---|---|---|---|---|---|---|---|
| **GUT_GENOME096174** | **Bacteroides plebeius DSM 17135** | **ISO** | GI | 100 | 100 | 100 | 100 | 100 | 100 |
| GUT_GENOME154239 | Bacteroides cellulosilyticus | MAG | GI | 100 | 100 | 100 | 100 | 100 | 99.8 |
| GUT_GENOME099338 | Bacteroides coprocola | MAG | GI | 100 | 100 | 100 | 100 | 100 | 100 |
| KIJ_genome 1302 | Bacteroides coprocola | MAG | GI | 100 | 99.5 | 100 | 100 | 100 | 99.8 |
| GUT_GENOME001074 | Bacteroides dorei AF25-30LB | ISO | GI | 100 | 100 | 100 | 100 | 100 | 99.8 |
| GUT_GENOME032129 | Bacteroides dorei | MAG | GI | 100 | 100 | 100 | 100 | 99.9 | 99.8 |
| GUT_GENOME239769 | Bacteroides dorei TM05-24 | ISO | GI | 100 | 100 | 100 | 100 | 99.9 | 100 |
| GUT_GENOME037305 | Bacteroides dorei | MAG | GI | 100 | 99.5 | 100 | 100 | 100 | 99.9 |
| GUT_GENOME154531 | Bacteroides dorei | MAG | GI | 100 | 100 | 100 | 100 | 100 | 100 |
| GUT_GENOME001766 | Bacteroides eggerthii AM42-16 | ISO | GI | 100 | 100 | 100 | 100 | 100 | 99.8 |
| KIJ_genome 5720 | Bacteroides eggerthii | MAG | GI | 100 | 99.5 | 100 | 100 | 100 | 99.9 |
| GUT_GENOME155110 | Bacteroides eggerthii | MAG | GI | 100 | 100 | 100 | 100 | 100 | 100 |
| KIJ_genome 13110 | Bacteroides finegoldii | MAG | GI | 100 | 99.5 | 100 | 100 | 99.9 | 99.9 |
| GUT_GENOME098104 | Bacteroides finegoldii | MAG | GI | 100 | 99.5 | 100 | 100 | 100 | 99.9 |
| GUT_GENOME249472 | Bacteroides finegoldii | MAG | GI | 100 | 100 | 100 | 100 | 100 | 100 |
| GUT_GENOME101376 | Bacteroides massiliensis | MAG | GI | 100 | 99.5 | 100 | 100 | 100 | 99.9 |
| GUT_GENOME000819 | Bacteroides ovatus AF14-13AC | ISO | GI | 100 | 100 | 100 | 100 | 100 | 99.9 |
| GUT_GENOME030187 | Bacteroides ovatus | MAG | GI | 100 | 100 | 100 | 100 | 100 | 99.8 |
| GUT_GENOME152759 | Bacteroides ovatus | MAG | GI | 100 | 99.5 | 100 | 100 | 100 | 99.9 |
| KIJ_genome 25909 | Bacteroides plebeius | MAG | GI | 100 | 100 | 100 | 100 | 100 | 99.9 |
| GUT_GENOME034154 | Bacteroides plebeius | MAG | GI | 100 | 99.5 | 100 | 100 | 100 | 99.8 |
| GUT_GENOME250348 | Bacteroides plebeius | MAG | GI | 100 | 99.5 | 100 | 100 | 100 | 99.9 |
| GUT_GENOME153158 | Bacteroides plebeius | MAG | GI | 100 | 100 | 100 | 100 | 99.9 | 100 |
| GUT_GENOME001642 | Bacteroides uniformis AM30-49 | ISO | GI | 100 | 100 | 100 | 100 | 100 | 100 |
| GUT_GENOME000904 | Bacteroides uniformis AF17-20 | ISO | GI | 100 | 99.5 | 100 | 100 | 100 | 99.9 |
| KIJ_genome 736 | Bacteroides uniformis | MAG | GI | 100 | 99.5 | 100 | 100 | 99.9 | 99.9 |
| GUT_GENOME155200 | Bacteroides uniformis | MAG | GI | 100 | 100 | 100 | 100 | 100 | 99.8 |
| GUT_GENOME000851 | Bacteroides uniformis AF15-14LB | ISO | GI | 100 | 100 | 100 | 100 | 100 | 100 |
| GUT_GENOME000893 | Bacteroides uniformis AF16-7 | ISO | GI | 100 | 100 | 100 | 100 | 100 | 100 |
| GUT_GENOME000983 | Bacteroides uniformis AF20-13LB | ISO | GI | 100 | 100 | 100 | 100 | 100 | 99.8 |
| GUT_GENOME170222 | Bacteroides uniformis | MAG | GI | 100 | 99.3 | 100 | 100 | 100 | 99.9 |
| GUT_GENOME239800 | Bacteroides uniformis TM09-11 | ISO | GI | 100 | 100 | 100 | 100 | 100 | 100 |
| GUT_GENOME001710 | Bacteroides stercoris AM36-9BH | ISO | GI | 100 | 100 | 100 | 100 | 100 | 100 |
| KIJ_genome 10066 | Bacteroides stercoris | MAG | GI | 100 | 100 | 100 | 100 | 100 | 99.9 |
| KIJ_genome 11367 | Bacteroides vulgatus | MAG | GI | 100 | 100 | 100 | 100 | 100 | 99.9 |
| GUT_GENOME269537 | Bacteroides xylanisolvens | MAG | GI | 100 | 100 | 100 | 100 | 100 | 100 |
| GUT_GENOME178534 | Bacteroides xylanisolvens | MAG | GI | 100 | 99.5 | 100 | 100 | 100 | 99.9 |
| KIJ_genome 23951 | Parabacteroides merdae | MAG | GI | 100 | 100 | 100 | 100 | 99.9 | 100 |
| GUT_GENOME027847 | Bacteroides dorei | MAG | GI | 100 | 99.3 | 100 | 100 | 100 | 99.8 |
| KIJ_genome 12929 | Tyzzerella sp000411335 | MAG | GI | 73.2 | 100 | 100 | 100 | 99.9 | 99.9 |
| KIJ_genome 24121 | Bacteroides coprocola | MAG | GI | 98.3 | 100 | 100 | 99.4 | 99.8 | 99.6 |
| GUT_GENOME178548 | Bacteroides ovatus | MAG | GI | 99.4 | 100 | 100 | 100 | 100 | 99.6 |
| GUT_GENOME239713 | Bacteroides uniformis TF09-22 | ISO | GI | 99.4 | 99 | 99.7 | 100 | 98.8 | 99.8 |
| KIJ_genome 22831 | Bacteroides plebeius | MAG | Outgroup GI | 99.6 | 99.5 | 99.7 | 99.8 | 99.9 | 99.9 |
| GUT_GENOME001839 | Bacteroides plebeius AM49-7BH | ISO | Outgroup GI | 99.8 | 99.3 | 99.7 | 99.8 | 99.9 | 99.9 |
| KIJ_genome 28353 | Bacteroides vulgatus | MAG | Outgroup GI | 99.4 | 99 | 98 | 100 | 100 | 99.9 |
| KIJ_genome 22636 | Bacteroides uniformis | MAG | Outgroup GI | 100 | 98.8 | 96.7 | 98.9 | 99.3 | 99.5 |
| KIJ_genome 7975 | Bacteroides plebeius | MAG | Outgroup GI | 100 | 100 | 100 | 98.2 | 98.2 | 100 |
| GUT_GENOME229839 | Bacteroides stercorirosoris | MAG | Outgroup GII | 99.8 | 99 | 96.7 | 97.4 | 98.2 | 99.5 |
| GUT_GENOME034168 | Bacteroides caccae | MAG | GII | 99.8 | 99 | 96.7 | 97.4 | 98.2 | 98.7 |
| GUT_GENOME149815 | Bacteroides caccae | MAG | GII | 99.4 | 99 | 96.7 | 97.6 | 98.2 | 98.7 |
| KIJ_genome 15612 | Bacteroides coprocola | MAG | GII | 99.6 | 99 | 96.7 | 97.6 | 98.2 | 98.8 |
| KIJ_genome 27310 | Bacteroides coprocola | MAG | GII | 99.4 | 99 | 96.7 | 97.4 | 98.2 | 98.7 |
| KIJ_genome 20890 | Bacteroides dorei | MAG | GII | 99.8 | 98.8 | 96.7 | 97.4 | 98.2 | 98.8 |
| KIJ_genome 2914 | Bacteroides dorei | MAG | GII | 99.8 | 99 | 96.7 | 97.6 | 98.1 | 98.8 |
| KIJ_genome 22256 | Bacteroides dorei | MAG | GII | 99.4 | 99 | 96.3 | 97.6 | 98.2 | 98.7 |
| GUT_GENOME151759 | Bacteroides dorei | MAG | GII | 99.4 | 99 | 96.3 | 97.6 | 98.2 | 98.7 |
| GUT_GENOME027742 | Bacteroides dorei | MAG | GII | 99.4 | 99 | 96.7 | 97.4 | 98.2 | 98.7 |
| GUT_GENOME027786 | Bacteroides dorei | MAG | GII | 99.8 | 99 | 96.7 | 97.4 | 98.2 | 98.8 |
| GUT_GENOME028138 | Bacteroides dorei | MAG | GII | 99.8 | 98.8 | 96.7 | 97.4 | 98.2 | 98.8 |
| GUT_GENOME046102 | Bacteroides finegoldii | MAG | GII | 99.4 | 99 | 96.7 | 97.6 | 98.2 | 98.7 |
| GUT_GENOME080948 | Bacteroides fragilis | MAG | GII | 99.8 | 99 | 96.7 | 97.6 | 98.2 | 98.8 |

| | | | | | | | | | |
|---|---|---|---|---|---|---|---|---|---|
| GUT_GENOME228260 | Bacteroides fragilis | MAG | GII | 99.8 | 99 | 96.3 | 97.6 | 98.2 | 98.7 |
| KIJ_genome 14857 | Bacteroides ilei | MAG | GII | 99.8 | 99 | 96.7 | 97.4 | 98.2 | 98.8 |
| KIJ_genome 16412 | Bacteroides massiliensis | MAG | GII | 99.8 | 99 | 96.7 | 97.4 | 98.2 | 98.8 |
| GUT_GENOME148452 | Bacteroides massiliensis | MAG | GII | 99.4 | 99 | 96.3 | 97.6 | 98.2 | 98.7 |
| KIJ_genome 17159 | Bacteroides plebeius | MAG | GII | 99.8 | 99 | 96.7 | 97.4 | 98.2 | 98.8 |
| KIJ_genome 24998 | Bacteroides plebeius | MAG | GII | 99.8 | 99 | 96.7 | 98.1 | 98.2 | 98.8 |
| KIJ_genome 157 | Bacteroides plebeius | MAG | GII | 99.4 | 99 | 96.7 | 97.4 | 98.2 | 98.7 |
| KIJ_genome 26449 | Bacteroides plebeius | MAG | GII | 99.4 | 99 | 96.3 | 97.6 | 98.2 | 98.7 |
| GUT_GENOME148536 | Bacteroides plebeius | MAG | GII | 99.4 | 99 | 96.7 | 97.4 | 98.2 | 98.7 |
| GUT_GENOME001661 | Bacteroides stercoris AM32-16LB | ISO | GII | 99.4 | 99 | 96.7 | 97.4 | 98.2 | 98.7 |
| GUT_GENOME027726 | Bacteroides stercoris | MAG | GII | 99.8 | 99 | 96.7 | 97.6 | 98.1 | 98.8 |
| GUT_GENOME098019 | Bacteroides stercoris | MAG | GII | 99.8 | 99 | 96.7 | 97.6 | 98.2 | 98.8 |
| GUT_GENOME246849 | Bacteroides stercoris | MAG | GII | 99.8 | 99 | 96.7 | 97.4 | 98.2 | 98.8 |
| KIJ_genome 11227 | Bacteroides thetaiotaomicron | MAG | GII | 99.8 | 99 | 96.7 | 97.6 | 98.2 | 98.8 |
| GUT_GENOME003489 | Bacteroides thetaiotaomicron | MAG | GII | 99.8 | 99 | 96.7 | 97.4 | 98.2 | 98.8 |
| GUT_GENOME101535 | Bacteroides thetaiotaomicron | MAG | GII | 99.4 | 99 | 96.7 | 97.4 | 98.2 | 98.7 |
| KIJ_genome 26710 | Bacteroides uniformis | MAG | GII | 99.8 | 99 | 96.7 | 97.4 | 98.2 | 98.8 |
| KIJ_genome 6460 | Bacteroides uniformis | MAG | GII | 99.8 | 99 | 96.7 | 97.4 | 98.2 | 98.8 |
| KIJ_genome 9193 | Bacteroides uniformis | MAG | GII | 99.4 | 99 | 96.7 | 97.4 | 98.2 | 98.7 |
| KIJ_genome 4327 | Bacteroides uniformis | MAG | GII | 99.4 | 99 | 96.3 | 97.6 | 98.2 | 98.7 |
| KIJ_genome 4881 | Bacteroides uniformis | MAG | GII | 99.4 | 99 | 96.3 | 97.6 | 98.2 | 98.7 |
| GUT_GENOME001008 | Bacteroides uniformis AF21-53 | ISO | GII | 99.8 | 99 | 96.7 | 97.4 | 98.2 | 98.8 |
| GUT_GENOME001076 | Bacteroides uniformis AF25-38AC | ISO | GII | 99.4 | 99 | 96.7 | 97.4 | 98.2 | 98.7 |
| GUT_GENOME001081 | Bacteroides uniformis AF26-10BH | ISO | GII | 99.8 | 99 | 96.7 | 97.4 | 98.2 | 98.8 |
| GUT_GENOME001798 | Bacteroides uniformis AM43-9 | ISO | GII | 99.8 | 99 | 96.7 | 97.6 | 98.2 | 98.8 |
| GUT_GENOME016334 | Bacteroides uniformis | MAG | GII | 99.4 | 99 | 96.7 | 97.4 | 98.2 | 98.7 |
| GUT_GENOME033675 | Bacteroides uniformis | MAG | GII | 99.4 | 99 | 96.3 | 97.6 | 98.2 | 98.6 |
| GUT_GENOME147943 | Bacteroides uniformis | MAG | GII | 99.4 | 99 | 96.7 | 97.6 | 98.2 | 98.7 |
| GUT_GENOME150364 | Bacteroides uniformis | MAG | GII | 99.4 | 99 | 96.7 | 97.4 | 98.2 | 98.7 |
| GUT_GENOME151214 | Bacteroides uniformis | MAG | GII | 99.8 | 99 | 96.7 | 97.6 | 98.2 | 98.8 |
| GUT_GENOME178496 | Bacteroides uniformis | MAG | GII | 99.4 | 99 | 96.7 | 97.4 | 98.2 | 98.7 |
| GUT_GENOME227045 | Bacteroides uniformis | MAG | GII | 99.4 | 99 | 96.7 | 97.4 | 98.2 | 98.7 |
| KIJ_genome 1091 | Bacteroides vulgatus | MAG | GII | 99.8 | 99 | 96.7 | 97.4 | 98.2 | 98.8 |
| KIJ_genome 12389 | Bacteroides vulgatus | MAG | GII | 99.8 | 99 | 96.7 | 97.6 | 98.2 | 98.8 |
| KIJ_genome 15885 | Bacteroides vulgatus | MAG | GII | 99.8 | 99 | 96.7 | 97.2 | 98.1 | 98.8 |
| KIJ_genome 17209 | Bacteroides vulgatus | MAG | GII | 99.8 | 99 | 96.3 | 97.6 | 98.1 | 98.8 |
| KIJ_genome 10033 | Bacteroides vulgatus | MAG | GII | 99.4 | 99 | 96.7 | 97.2 | 98.2 | 98.7 |
| KIJ_genome 20177 | Bacteroides vulgatus | MAG | GII | 99.4 | 99 | 96.3 | 97.6 | 98.2 | 98.7 |
| KIJ_genome 22411 | Bacteroides xylanisolvens | MAG | GII | 99.4 | 99 | 96.3 | 97.6 | 98.2 | 98.7 |
| GUT_GENOME001249 | Bacteroides xylanisolvens AF38-2 | ISO | GII | 99.4 | 99 | 96.7 | 97.2 | 98.2 | 98.7 |
| GUT_GENOME153813 | Bacteroides xylanisolvens | MAG | GII | 99.8 | 99 | 96.7 | 97.4 | 98.2 | 98.8 |
| KIJ_genome 22412 | Bacteroides xylanisolvens | MAG | GII | 99.8 | 99 | 96.7 | 97.4 | 98.1 | 98.8 |
| KIJ_genome 15171 | Parabacteroides johnsonii | MAG | GII | 99.4 | 99 | 96.7 | 97.2 | 98.2 | 98.7 |
| KIJ_genome 17018 | Parabacteroides merdae | MAG | GII | 99.8 | 99 | 96.7 | 97.4 | 98.2 | 98.8 |
| GUT_GENOME099480 | Tyzzerella nexilis | MAG | GII | 99.4 | 99 | 96.7 | 97.2 | 98.2 | 98.7 |
| GUT_GENOME085027 | Bacteroides coprocola | MAG | GIIIrec (GI) | 98.3 | 97.6 | 95.7 | 99.8 | 99.9 | 99.9 |
| GUT_GENOME033354 | Bacteroides dorei | MAG | GIIIrec (GI) | 98.3 | 97.6 | 95.7 | 99.8 | 99.9 | 99.9 |
| GUT_GENOME001331 | Bacteroides plebeius AM09-36 | ISO | GIIIrec (GI) | 98.3 | 97.6 | 95.7 | 99.8 | 99.9 | 99.9 |
| GUT_GENOME017313 | Bacteroides coprocola | MAG | GIIIrec (GI) | 98.3 | 98 | 93 | 99.8 | 99.9 | 99.9 |
| GUT_GENOME150750 | Bacteroides plebeius | MAG | GIIIrec (GI) | 98.3 | 98 | 93 | 99.8 | 99.9 | 99.9 |
| GUT_GENOME249569 | Bacteroides plebeius | MAG | GIIIrec (GI) | 98.3 | 98 | 93 | 98.7 | 99.1 | 99.9 |
| KIJ_genome 9348 | Bacteroides plebeius | MAG | GIIIrec (GII) | 98.3 | 97.6 | 92 | 97.4 | 98.2 | 98.8 |
| KIJ_genome 101 | Bacteroides coprocola | MAG | GIII | 98.3 | 97.5 | 92 | 96.3 | 97.9 | 96 |
| KIJ_genome 26984 | Bacteroides coprocola | MAG | GIII | 98.3 | 97.7 | 92 | 96.3 | 97.9 | 96 |
| KIJ_genome 15823 | Bacteroides coprocola | MAG | GIII | 98.3 | 97.7 | 92.3 | 96.1 | 97.9 | 95.9 |
| GUT_GENOME032023 | Bacteroides coprocola | MAG | GIII | 98.3 | 97.7 | 92 | 96.3 | 97.9 | 96 |
| GUT_GENOME044522 | Bacteroides coprocola | MAG | GIII | 98.3 | 97.6 | 92.3 | 96.3 | 97.9 | 96 |
| KIJ_genome 7396 | Bacteroides plebeius | MAG | GIII | 98.5 | 98.1 | 92.3 | 97.1 | 97.9 | 96 |
| KIJ_genome 15369 | Bacteroides plebeius | MAG | GIII | 98.3 | 97.6 | 92.3 | 96.3 | 98 | 96 |
| KIJ_genome 9284 | Bacteroides plebeius | MAG | GIII | 98.3 | 97.6 | 92.3 | 96.1 | 97.9 | 96 |
| KIJ_genome 24917 | Bacteroides plebeius | MAG | GIII | 98.3 | 97.7 | 92.3 | 96.3 | 97.8 | 96 |
| KIJ_genome 19050 | Bacteroides plebeius | MAG | GIII | 98.3 | 97.6 | 92 | 96.3 | 97.9 | 96 |
| KIJ_genome 22909 | Bacteroides plebeius | MAG | GIII | 98.3 | 97.6 | 92 | 96.1 | 97.9 | 96 |
| KIJ_genome 26757 | Bacteroides plebeius | MAG | GIII | 98.3 | 97.6 | 92 | 96.3 | 97.9 | 96 |
| GUT_GENOME000762 | Bacteroides stercoris AF05-4 | ISO | GIII | 98.3 | 97.6 | 92.3 | 96.3 | 97.9 | 96 |
| KIJ_genome 20422 | Bacteroides vulgatus | MAG | GIII | 98.3 | 97.6 | 92.3 | 96.3 | 97.9 | 96 |
| KIJ_genome 24180 | Bacteroides vulgatus | MAG | GIII | 98.3 | 97.5 | 92 | 96.3 | 97.9 | 96 |
| KIJ_genome 24692 | Holdemanella sp002299315 | MAG | GIII | 98.3 | 97.7 | 92.3 | 97.2 | 97.9 | 96.2 |

Clade GI
Clade GII
Clade GIII
Outgroup

**Table S3:** List of assembly projects probed with the genes encoding *PUL-PorB* and list of SRA projects probed with the 50 nucleotides probes.

| Location of sampling | | Project (assembly) | Bio-samples | PUL porB | | Project (SRA) | Bio-samples | 50 n probes | |
|---|---|---|---|---|---|---|---|---|---|
| | | | | hits | % | | | hits | % |
| **CHINA** | Hangzhou | PRJEB24527 | 97 | 39 | 40 | PRJNA375935 | 211 | 125 | 59 |
| | Hangzhou | PRJEB29103 | 168 | 71 | 42 | PRJNA356102 | 168 | 106 | 63 |
| | Hangzhou | PRJEB26158 | 97 | 40 | 41 | PRJNA353560 | 97 | 54 | 56 |
| | Hangzhou | | | | | PRJEB6337 | 312 | 137 | 44 |
| | Hangzhou | | | | | PRJNA505228 | 50 | 27 | 54 |
| | Shenzen | PRJEB30046 | 370 | 135 | 36 | PRJNA422434 | 370 | 163 | 44 |
| | Shenzen | PRJEB26908 | 19 | 2 | 11 | PRJEB12669 | 20 | 7 | 35 |
| | Guangzhou | | | | | PRJEB18755 | 124 | 40 | 32 |
| | Guangzhou | | | | | PRJEB15371 | 122 | 58 | 48 |
| | Beijin | | | | | PRJNA401977 | 145 | 101 | 70 |
| | Tangshan | | | | | PRJEB13870 | 193 | 62 | 32 |
| | Hong-Kong | PRJEB24748 | 128 | 23 | 18 | PRJNA557323 | 564 | 356 | 63 |
| | Hunan | | | | | PRJNA349463 | 40 | 8 | 20 |
| | Many places* | PRJEB26167 | 150 | 32 | 21 | PRJNA356225 | 150 | 67 | 45 |
| | Inner Mongolia | | | | | PRJNA328899 | 47 | 5 | 11 |
| **MONGOLIA** | Ulan Bator/ Khentti / TUV province | | | | | PRJNA328899 | 63 | 2 | 3 |
| **JAPAN** | Tokyo | PRJEB26092 | 254 | 50 | 20 | PRJDB3601 | 255 | 133 | 52 |
| | Kanazawa | | | | | PRJNA517801 | 68 | 10 | 15 |
| | Tokyo | | | | | PRJDB7378 | 50 | 47 | 94 |
| | Tokyo | PRJDB4176 | 643 | 170 | 26 | PRJDB4176 | 645 | 548 | 85 |
| **KOREA** | Seoul | PRJNA678426 | 90 | 23 | 26 | PRJNA678426 | 106 | 96 | 91 |
| | NA | | | | | PRJEB17896 | 27 | 24 | 89 |
| **MALAYSIA** | Kuala Lumpur | | | | | PRJNA797994 | 56 | 9 | 16 |
| | Orang Asli | | | | | PRJNA797994 | 351 | 3 | 1 |
| **INDIA** | Bhopal | PRJEB33179 | 53 | 0 | 0 | PRJNA397112 | 53 | 0 | 0 |
| | Kasaragod | PRJEB33179 | 57 | 0 | 0 | PRJNA397112 | 57 | 0 | 0 |
| **BANGLADESH** | Dhaka | PRJEB22359 | 7 | 0 | 0 | | | | |
| **USA** | Lincoln | PRJEB26490 | 63 | 1 | 1.6 | PRJNA324129 | 87 | 1 | 1 |
| | Seattle | PRJEB22365 | 22 | 0 | 0 | | | | |
| | Saint Louis | PRJEB24849 | 387 | 0 | 0 | | | | |
| | HMP | PRJEB22283 | 717 | 5 | 0.7 | | | | |
| | Cheyenne tribe | PRJEB22552 | 37 | 0 | 0 | | | | |
| **PERU+USA** | | PRJEB24847 | 36+11 | 0 | 0 | PRJNA268964 | 36 | 0 | 0 |
| **MADAGASCAR** | | PREJB33766 | 112 | 0 | 0 | PRJNA485056 | 112 | 0 | 0 |
| **TANZANIA + ITALY** | | PREJB22391 | 27+11 | 0 | 0 | PRJNA278393 | 27+11 | 0 | 0 |

**Table S4:** List of the reconstructed PUL-PorB found in the assemblied human gut metagenomes of 591 individuals from China, Japan, Korea or USA (note that there is 764 hits because some individuals had multiple hits). Only full length genes were reported in the list. The genes were classified in Group I , II, III or

| Study | Location | Biosample | GH29 (Bacple_01702) | GH16 (Bacple_01703) | SusC (Bacple_01704) | SusD (Bacple_01705) | GH2 (Bacple_01706) |
|---|---|---|---|---|---|---|---|
| PRJEB24527 | Hangzhou (China) | SAMN06348158 | GI | GI | GI | GI | GI |
| PRJEB24527 | Hangzhou (China) | SAMN06348180 | | GI | GI | GI | GI |
| PRJEB24527 | Hangzhou (China) | SAMN06348180 | GI | | | | |
| PRJEB24527 | Hangzhou (China) | SAMN06348127 | GIIIrec | GIIIrec | GI | | |
| PRJEB24527 | Hangzhou (China) | SAMN06348139 | GII | GII | GII | GII | GII |
| PRJEB24527 | Hangzhou (China) | SAMN06348137 | GII | GII | GII | GII | GII |
| PRJEB24527 | Hangzhou (China) | SAMN06348159 | | GII | GII | | |
| PRJEB24527 | Hangzhou (China) | SAMN06348159 | GII | | | | |
| PRJEB24527 | Hangzhou (China) | SAMN06348194 | GI | GI | GI | | |
| PRJEB24527 | Hangzhou (China) | SAMN06348194 | | | | | GI |
| PRJEB24527 | Hangzhou (China) | SAMN06348136 | GIIIrec | GIII | GI | GI | GI |
| PRJEB24527 | Hangzhou (China) | SAMN06348110 | | | | | GII |
| PRJEB24527 | Hangzhou (China) | SAMN06348110 | GII | GII | GII | GII | |
| PRJEB24527 | Hangzhou (China) | SAMN06348142 | | GI | | | |
| PRJEB24527 | Hangzhou (China) | SAMN06348196 | GI | | GI | GI | GI |
| PRJEB24527 | Hangzhou (China) | SAMN06348117 | GI | | | | |
| PRJEB24527 | Hangzhou (China) | SAMN06348117 | | GI | GI | GI | GI |
| PRJEB24527 | Hangzhou (China) | SAMN06348105 | GIIIrec | GIII | GI | GI | GI |
| PRJEB24527 | Hangzhou (China) | SAMN06348134 | GI | GI | GI | GI | GI |
| PRJEB24527 | Hangzhou (China) | SAMN06348145 | GII | GII | GII | | |
| PRJEB24527 | Hangzhou (China) | SAMN06348167 | GII | GII | GII | GII | |
| PRJEB24527 | Hangzhou (China) | SAMN06348115 | GI | GI | GI | GI | GI |
| PRJEB24527 | Hangzhou (China) | SAMN06348140 | | GI | | | |
| PRJEB24527 | Hangzhou (China) | SAMN06348168 | GIIIrec | GIIIrec | GI | GI | GI |
| PRJEB24527 | Hangzhou (China) | SAMN06348186 | GII | GII | GII | GII | GII |
| PRJEB24527 | Hangzhou (China) | SAMN06348183 | | GI | | | |
| PRJEB24527 | Hangzhou (China) | SAMN06348193 | GI | | GI | GI | GI |
| PRJEB24527 | Hangzhou (China) | SAMN06348190 | GI | GI | GI | GI | GI |
| PRJEB24527 | Hangzhou (China) | SAMN06348123 | GII | GII | GII | GII | GII |
| PRJEB24527 | Hangzhou (China) | SAMN06348149 | | | | | GI |
| PRJEB24527 | Hangzhou (China) | SAMN06348149 | GI | GI | GI | | |
| PRJEB24527 | Hangzhou (China) | SAMN06348164 | GI | | | | |
| PRJEB24527 | Hangzhou (China) | SAMN06348164 | | | | | GI |
| PRJEB24527 | Hangzhou (China) | SAMN06348164 | | GI | GI | | |
| PRJEB24527 | Hangzhou (China) | SAMN06348124 | GI | GI | | | |
| PRJEB24527 | Hangzhou (China) | SAMN06348124 | | | | GI | |
| PRJEB24527 | Hangzhou (China) | SAMN06348192 | GII | GII | GII | | |
| PRJEB24527 | Hangzhou (China) | SAMN06348176 | GII | | | | |
| PRJEB24527 | Hangzhou (China) | SAMN06348176 | | GI | GI | | |
| PRJEB24527 | Hangzhou (China) | SAMN06348185 | GI | | | | |
| PRJEB24527 | Hangzhou (China) | SAMN06348143 | | | | | GI |
| PRJEB24527 | Hangzhou (China) | SAMN06348156 | GII | GII | GII | GII | GII |
| PRJEB24527 | Hangzhou (China) | SAMN06348175 | GII | GII | GII | | |
| PRJEB24527 | Hangzhou (China) | SAMN06348175 | | | | | GII |
| PRJEB24527 | Hangzhou (China) | SAMN06348152 | | GI | GI | | |
| PRJEB24527 | Hangzhou (China) | SAMN06348174 | | | GI | GI | GI |
| PRJEB24527 | Hangzhou (China) | SAMN06348150 | GIIIrec | GIII | GI | GI | GI |
| PRJEB24527 | Hangzhou (China) | SAMN06348113 | GI | GI | GI | | |
| PRJEB24527 | Hangzhou (China) | SAMN06348166 | | | | | GI |
| PRJEB24527 | Hangzhou (China) | SAMN06348166 | GI | | | | |
| PRJEB24527 | Hangzhou (China) | SAMN06348166 | | GI | GI | | |

GI
GII
GIII
GIIIrec
Outgroup

| Study | Location | Biosample | GH29 (Bacple_01702) | GH16 (Bacple_01703) | SusC (Bacple_01704) | SusD (Bacple_01705) | GH2 (Bacple_01706) |
|---|---|---|---|---|---|---|---|
| PRJEB24527 | Hangzhou (China) | SAMN06348146 | | | | | |
| PRJEB26158 | Hangzhou (China) | SAMN06016365 | | | | | |
| PRJEB26158 | Hangzhou (China) | SAMN06016335 | | | | | |
| PRJEB26158 | Hangzhou (China) | SAMN06016335 | | | | | |
| PRJEB26158 | Hangzhou (China) | SAMN06016357 | | | | | |
| PRJEB26158 | Hangzhou (China) | SAMN06016417 | | | | | |
| PRJEB26158 | Hangzhou (China) | SAMN06016417 | | | | | |
| PRJEB26158 | Hangzhou (China) | SAMN06016417 | | | | | |
| PRJEB26158 | Hangzhou (China) | SAMN06016394 | | | | | |
| PRJEB26158 | Hangzhou (China) | SAMN06016376 | | | | | |
| PRJEB26158 | Hangzhou (China) | SAMN06016349 | | | | | |
| PRJEB26158 | Hangzhou (China) | SAMN06016375 | | | | | |
| PRJEB26158 | Hangzhou (China) | SAMN06016368 | | | | | |
| PRJEB26158 | Hangzhou (China) | SAMN06016400 | | | | | |
| PRJEB26158 | Hangzhou (China) | SAMN06016403 | | | | | |
| PRJEB26158 | Hangzhou (China) | SAMN06016395 | | | | | |
| PRJEB26158 | Hangzhou (China) | SAMN06016332 | | | | | |
| PRJEB26158 | Hangzhou (China) | SAMN06016333 | | | | | |
| PRJEB26158 | Hangzhou (China) | SAMN06016407 | | | | | |
| PRJEB26158 | Hangzhou (China) | SAMN06016369 | | | | | |
| PRJEB26158 | Hangzhou (China) | SAMN06016356 | | | | | |
| PRJEB26158 | Hangzhou (China) | SAMN06016341 | | | | | |
| PRJEB26158 | Hangzhou (China) | SAMN06016327 | | | | | |
| PRJEB26158 | Hangzhou (China) | SAMN06016354 | | | | | |
| PRJEB26158 | Hangzhou (China) | SAMN06016409 | | | | | |
| PRJEB26158 | Hangzhou (China) | SAMN06016409 | | | | | |
| PRJEB26158 | Hangzhou (China) | SAMN06016383 | | | | | |
| PRJEB26158 | Hangzhou (China) | SAMN06016383 | | | | | |
| PRJEB26158 | Hangzhou (China) | SAMN06016384 | | | | | |
| PRJEB26158 | Hangzhou (China) | SAMN06016384 | | | | | |
| PRJEB26158 | Hangzhou (China) | SAMN06016415 | | | | | |
| PRJEB26158 | Hangzhou (China) | SAMN06016420 | | | | | |
| PRJEB26158 | Hangzhou (China) | SAMN06016343 | | | | | |
| PRJEB26158 | Hangzhou (China) | SAMN06016411 | | | | | |
| PRJEB26158 | Hangzhou (China) | SAMN06016355 | | | | | |
| PRJEB26158 | Hangzhou (China) | SAMN06016355 | | | | | |
| PRJEB26158 | Hangzhou (China) | SAMN06016416 | | | | | |
| PRJEB26158 | Hangzhou (China) | SAMN06016416 | | | | | |
| PRJEB26158 | Hangzhou (China) | SAMN06016351 | | | | | |
| PRJEB26158 | Hangzhou (China) | SAMN06016377 | | | | | |
| PRJEB26158 | Hangzhou (China) | SAMN06016377 | | | | | |
| PRJEB26158 | Hangzhou (China) | SAMN06016406 | | | | | |
| PRJEB26158 | Hangzhou (China) | SAMN06016412 | | | | | |
| PRJEB26158 | Hangzhou (China) | SAMN06016412 | | | | | |
| PRJEB26158 | Hangzhou (China) | SAMN06016352 | | | | | |
| PRJEB26158 | Hangzhou (China) | SAMN06016389 | | | | | |
| PRJEB26158 | Hangzhou (China) | SAMN06016360 | | | | | |
| PRJEB26158 | Hangzhou (China) | SAMN06016391 | | | | | |
| PRJEB26158 | Hangzhou (China) | SAMN06016391 | | | | | |
| PRJEB26158 | Hangzhou (China) | SAMN06016362 | | | | | |
| PRJEB26158 | Hangzhou (China) | SAMN06016340 | | | | | |
| PRJEB26158 | Hangzhou (China) | SAMN06016364 | | | | | |
| PRJEB29103 | Hangzhou (China) | SAMN06111977 | | | | | |
| PRJEB29103 | Hangzhou (China) | SAMN06111977 | | | | | |
| PRJEB29103 | Hangzhou (China) | SAMN06112032 | | | | | |
| PRJEB29103 | Hangzhou (China) | SAMN06112032 | | | | | |
| PRJEB29103 | Hangzhou (China) | SAMN06112020 | | | | | |

| Study | Location | Biosample |
| --- | --- | --- |
| PRJEB29103 | Hangzhou (China) | SAMN06112008 |
| PRJEB29103 | Hangzhou (China) | SAMN06112008 |
| PRJEB29103 | Hangzhou (China) | SAMN06112024 |
| PRJEB29103 | Hangzhou (China) | SAMN06111979 |
| PRJEB29103 | Hangzhou (China) | SAMN06112084 |
| PRJEB29103 | Hangzhou (China) | SAMN06112084 |
| PRJEB29103 | Hangzhou (China) | SAMN06112031 |
| PRJEB29103 | Hangzhou (China) | SAMN06112031 |
| PRJEB29103 | Hangzhou (China) | SAMN06112010 |
| PRJEB29103 | Hangzhou (China) | SAMN06112010 |
| PRJEB29103 | Hangzhou (China) | SAMN06112010 |
| PRJEB29103 | Hangzhou (China) | SAMN06111975 |
| PRJEB29103 | Hangzhou (China) | SAMN06111975 |
| PRJEB29103 | Hangzhou (China) | SAMN06112021 |
| PRJEB29103 | Hangzhou (China) | SAMN06112021 |
| PRJEB29103 | Hangzhou (China) | SAMN06112021 |
| PRJEB29103 | Hangzhou (China) | SAMN06111969 |
| PRJEB29103 | Hangzhou (China) | SAMN06111969 |
| PRJEB29103 | Hangzhou (China) | SAMN06111989 |
| PRJEB29103 | Hangzhou (China) | SAMN06111989 |
| PRJEB29103 | Hangzhou (China) | SAMN06112051 |
| PRJEB29103 | Hangzhou (China) | SAMN06111957 |
| PRJEB29103 | Hangzhou (China) | SAMN06705192 |
| PRJEB29103 | Hangzhou (China) | SAMN06111983 |
| PRJEB29103 | Hangzhou (China) | SAMN06111992 |
| PRJEB29103 | Hangzhou (China) | SAMN06111992 |
| PRJEB29103 | Hangzhou (China) | SAMN06111961 |
| PRJEB29103 | Hangzhou (China) | SAMN06112053 |
| PRJEB29103 | Hangzhou (China) | SAMN06111984 |
| PRJEB29103 | Hangzhou (China) | SAMN06111995 |
| PRJEB29103 | Hangzhou (China) | SAMN06111995 |
| PRJEB29103 | Hangzhou (China) | SAMN06112015 |
| PRJEB29103 | Hangzhou (China) | SAMN06111970 |
| PRJEB29103 | Hangzhou (China) | SAMN06111970 |
| PRJEB29103 | Hangzhou (China) | SAMN06111970 |
| PRJEB29103 | Hangzhou (China) | SAMN06111999 |
| PRJEB29103 | Hangzhou (China) | SAMN06111999 |
| PRJEB29103 | Hangzhou (China) | SAMN06111999 |
| PRJEB29103 | Hangzhou (China) | SAMN06111931 |
| PRJEB29103 | Hangzhou (China) | SAMN06112046 |
| PRJEB29103 | Hangzhou (China) | SAMN06112027 |
| PRJEB29103 | Hangzhou (China) | SAMN06112034 |
| PRJEB29103 | Hangzhou (China) | SAMN06112034 |
| PRJEB29103 | Hangzhou (China) | SAMN06112034 |
| PRJEB29103 | Hangzhou (China) | SAMN06112000 |
| PRJEB29103 | Hangzhou (China) | SAMN06112044 |
| PRJEB29103 | Hangzhou (China) | SAMN06112044 |
| PRJEB29103 | Hangzhou (China) | SAMN06112044 |
| PRJEB29103 | Hangzhou (China) | SAMN06112057 |
| PRJEB29103 | Hangzhou (China) | SAMN06111937 |
| PRJEB29103 | Hangzhou (China) | SAMN06111937 |
| PRJEB29103 | Hangzhou (China) | SAMN06111948 |
| PRJEB29103 | Hangzhou (China) | SAMN06111948 |
| PRJEB29103 | Hangzhou (China) | SAMN06112019 |
| PRJEB29103 | Hangzhou (China) | SAMN06112019 |
| PRJEB29103 | Hangzhou (China) | SAMN06111946 |
| PRJEB29103 | Hangzhou (China) | SAMN06111946 |

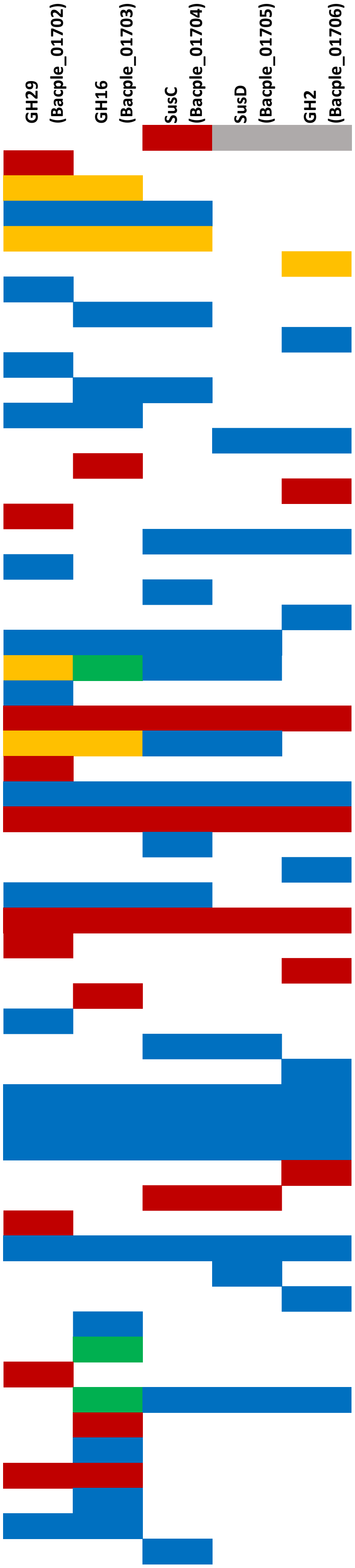

| Study | Location | Biosample |
|---|---|---|
| PRJEB29103 | Hangzhou (China) | SAMN06112061 |
| PRJEB29103 | Hangzhou (China) | SAMN06112061 |
| PRJEB29103 | Hangzhou (China) | SAMN06111988 |
| PRJEB29103 | Hangzhou (China) | SAMN06111990 |
| PRJEB29103 | Hangzhou (China) | SAMN06111991 |
| PRJEB29103 | Hangzhou (China) | SAMN06111996 |
| PRJEB29103 | Hangzhou (China) | SAMN06112052 |
| PRJEB29103 | Hangzhou (China) | SAMN06112042 |
| PRJEB29103 | Hangzhou (China) | SAMN06111962 |
| PRJEB29103 | Hangzhou (China) | SAMN06111962 |
| PRJEB29103 | Hangzhou (China) | SAMN06111963 |
| PRJEB29103 | Hangzhou (China) | SAMN06112005 |
| PRJEB29103 | Hangzhou (China) | SAMN06112050 |
| PRJEB29103 | Hangzhou (China) | SAMN06112050 |
| PRJEB29103 | Hangzhou (China) | SAMN06112002 |
| PRJEB29103 | Hangzhou (China) | SAMN06111997 |
| PRJEB29103 | Hangzhou (China) | SAMN06111934 |
| PRJEB29103 | Hangzhou (China) | SAMN06111950 |
| PRJEB29103 | Hangzhou (China) | SAMN06111943 |
| PRJEB29103 | Hangzhou (China) | SAMN06111943 |
| PRJEB29103 | Hangzhou (China) | SAMN06112094 |
| PRJEB29103 | Hangzhou (China) | SAMN06112058 |
| PRJEB29103 | Hangzhou (China) | SAMN06112040 |
| PRJEB29103 | Hangzhou (China) | SAMN06112040 |
| PRJEB29103 | Hangzhou (China) | SAMN06112001 |
| PRJEB29103 | Hangzhou (China) | SAMN06112001 |
| PRJEB29103 | Hangzhou (China) | SAMN06112017 |
| PRJEB29103 | Hangzhou (China) | SAMN06112069 |
| PRJEB29103 | Hangzhou (China) | SAMN06111971 |
| PRJEB29103 | Hangzhou (China) | SAMN06111953 |
| PRJEB29103 | Hangzhou (China) | SAMN06111945 |
| PRJEB29103 | Hangzhou (China) | SAMN06111929 |
| PRJEB29103 | Hangzhou (China) | SAMN06112083 |
| PRJEB29103 | Hangzhou (China) | SAMN06111980 |
| PRJEB29103 | Hangzhou (China) | SAMN06112060 |
| PRJEB29103 | Hangzhou (China) | SAMN06112060 |
| PRJEB29103 | Hangzhou (China) | SAMN06111981 |
| PRJEB29103 | Hangzhou (China) | SAMN06111967 |
| PRJEB29103 | Hangzhou (China) | SAMN06112070 |
| PRJEB29103 | Hangzhou (China) | SAMN06112070 |
| PRJEB29103 | Hangzhou (China) | SAMN06111986 |
| PRJEB29103 | Hangzhou (China) | SAMN06112073 |
| PRJEB29103 | Hangzhou (China) | SAMN06112012 |
| PRJEB29103 | Hangzhou (China) | SAMN06112012 |
| PRJEB29103 | Hangzhou (China) | SAMN06112036 |
| PRJEB24748 | Hong Kong (China) | SAMEA3541511 |
| PRJEB24748 | Hong Kong (China) | SAMEA3541481 |
| PRJEB24748 | Hong Kong (China) | SAMEA3541543 |
| PRJEB24748 | Hong Kong (China) | SAMEA3541539 |
| PRJEB24748 | Hong Kong (China) | SAMEA3541586 |
| PRJEB24748 | Hong Kong (China) | SAMEA3541586 |
| PRJEB24748 | Hong Kong (China) | SAMEA3541592 |
| PRJEB24748 | Hong Kong (China) | SAMEA3541577 |
| PRJEB24748 | Hong Kong (China) | SAMEA3541522 |
| PRJEB24748 | Hong Kong (China) | SAMEA3541551 |
| PRJEB24748 | Hong Kong (China) | SAMEA3541559 |
| PRJEB24748 | Hong Kong (China) | SAMEA3541570 |

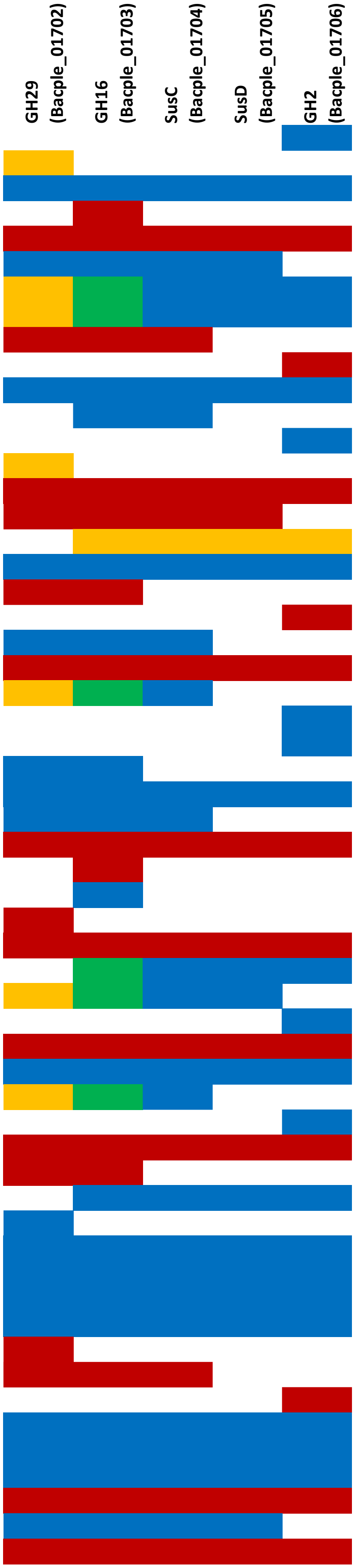

| Study | Location | Biosample |
|---|---|---|
| PRJEB24748 | Hong Kong (China) | SAMEA3541556 |
| PRJEB24748 | Hong Kong (China) | SAMEA3541516 |
| PRJEB24748 | Hong Kong (China) | SAMEA3541553 |
| PRJEB24748 | Hong Kong (China) | SAMEA3541580 |
| PRJEB24748 | Hong Kong (China) | SAMEA3541519 |
| PRJEB24748 | Hong Kong (China) | SAMEA3541493 |
| PRJEB24748 | Hong Kong (China) | SAMEA3541531 |
| PRJEB24748 | Hong Kong (China) | SAMEA3541554 |
| PRJEB24748 | Hong Kong (China) | SAMEA3541585 |
| PRJEB24748 | Hong Kong (China) | SAMEA3541585 |
| PRJEB24748 | Hong Kong (China) | SAMEA3541472 |
| PRJEB24748 | Hong Kong (China) | SAMEA3541557 |
| PRJEB24748 | Hong Kong (China) | SAMEA3541557 |
| PRJEB24748 | Hong Kong (China) | SAMEA3541524 |
| PRJEB24748 | Hong Kong (China) | SAMEA3541524 |
| PRJEB26908 | Shenzen (China) | SAMN02841183 |
| PRJEB26908 | Shenzen (China) | SAMEA4031715 |
| PRJEB30046 | Shenzen (China) | SAMN00791943 |
| PRJEB30046 | Shenzen (China) | SAMN00791936 |
| PRJEB30046 | Shenzen (China) | SAMN00791921 |
| PRJEB30046 | Shenzen (China) | SAMN00791910 |
| PRJEB30046 | Shenzen (China) | SAMN00791920 |
| PRJEB30046 | Shenzen (China) | SAMN00791914 |
| PRJEB30046 | Shenzen (China) | SAMN00791914 |
| PRJEB30046 | Shenzen (China) | SAMN00791914 |
| PRJEB30046 | Shenzen (China) | SAMN00791915 |
| PRJEB30046 | Shenzen (China) | SAMN00791915 |
| PRJEB30046 | Shenzen (China) | SAMN00791927 |
| PRJEB30046 | Shenzen (China) | SAMN00791927 |
| PRJEB30046 | Shenzen (China) | SAMN00791913 |
| PRJEB30046 | Shenzen (China) | SAMN00791937 |
| PRJEB30046 | Shenzen (China) | SAMN00791934 |
| PRJEB30046 | Shenzen (China) | SAMN00791919 |
| PRJEB30046 | Shenzen (China) | SAMN00791918 |
| PRJEB30046 | Shenzen (China) | SAMN00791911 |
| PRJEB30046 | Shenzen (China) | SAMN00791911 |
| PRJEB30046 | Shenzen (China) | SAMN00791911 |
| PRJEB30046 | Shenzen (China) | SAMN00791904 |
| PRJEB30046 | Shenzen (China) | SAMN00791923 |
| PRJEB30046 | Shenzen (China) | SAMN00791907 |
| PRJEB30046 | Shenzen (China) | SAMN00791931 |
| PRJEB30046 | Shenzen (China) | SAMN00791912 |
| PRJEB30046 | Shenzen (China) | SAMN00715199 |
| PRJEB30046 | Shenzen (China) | SAMN00791909 |
| PRJEB30046 | Shenzen (China) | SAMN00715191 |
| PRJEB30046 | Shenzen (China) | SAMN00715184 |
| PRJEB30046 | Shenzen (China) | SAMN00715161 |
| PRJEB30046 | Shenzen (China) | SAMN00715215 |
| PRJEB30046 | Shenzen (China) | SAMN00715158 |
| PRJEB30046 | Shenzen (China) | SAMN00715187 |
| PRJEB30046 | Shenzen (China) | SAMN00715170 |
| PRJEB30046 | Shenzen (China) | SAMN00715242 |
| PRJEB30046 | Shenzen (China) | SAMN00715163 |
| PRJEB30046 | Shenzen (China) | SAMN00715163 |
| PRJEB30046 | Shenzen (China) | SAMN00715203 |
| PRJEB30046 | Shenzen (China) | SAMN00715203 |
| PRJEB30046 | Shenzen (China) | SAMN00715233 |

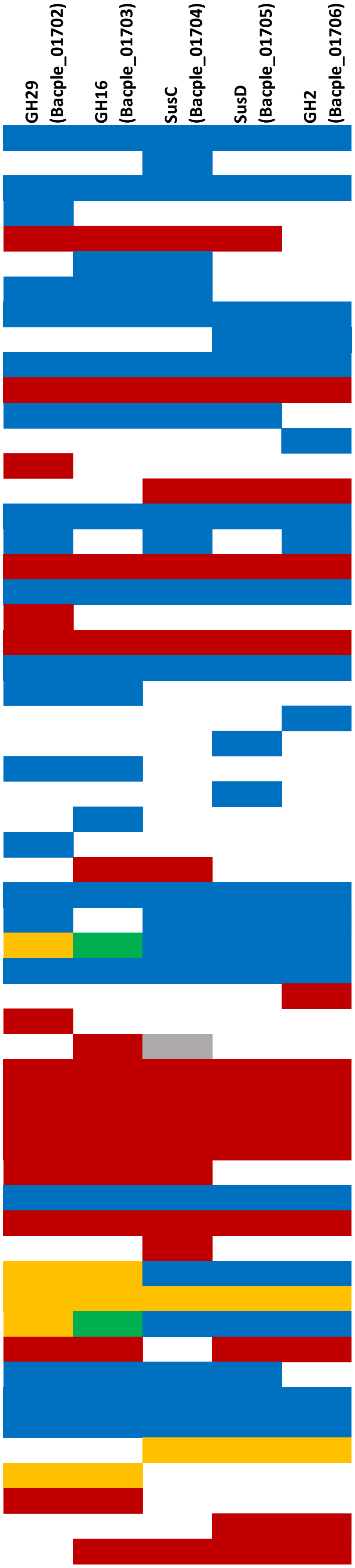

| Study | Location | Biosample |
|---|---|---|
| PRJEB30046 | Shenzen (China) | SAMN00715233 |
| PRJEB30046 | Shenzen (China) | SAMN00715237 |
| PRJEB30046 | Shenzen (China) | SAMN00715217 |
| PRJEB30046 | Shenzen (China) | SAMN00715217 |
| PRJEB30046 | Shenzen (China) | SAMN00715192 |
| PRJEB30046 | Shenzen (China) | SAMN00715225 |
| PRJEB30046 | Shenzen (China) | SAMN00715225 |
| PRJEB30046 | Shenzen (China) | SAMN00715196 |
| PRJEB30046 | Shenzen (China) | SAMN00715231 |
| PRJEB30046 | Shenzen (China) | SAMN00715168 |
| PRJEB30046 | Shenzen (China) | SAMN00715168 |
| PRJEB30046 | Shenzen (China) | SAMN00715239 |
| PRJEB30046 | Shenzen (China) | SAMN00715238 |
| PRJEB30046 | Shenzen (China) | SAMN00715174 |
| PRJEB30046 | Shenzen (China) | SAMN00715197 |
| PRJEB30046 | Shenzen (China) | SAMN00715224 |
| PRJEB30046 | Shenzen (China) | SAMN00715171 |
| PRJEB30046 | Shenzen (China) | SAMN00715241 |
| PRJEB30046 | Shenzen (China) | SAMN00715176 |
| PRJEB30046 | Shenzen (China) | SAMN00715173 |
| PRJEB30046 | Shenzen (China) | SAMN00715220 |
| PRJEB30046 | Shenzen (China) | SAMN00715234 |
| PRJEB30046 | Shenzen (China) | SAMN00715219 |
| PRJEB30046 | Shenzen (China) | SAMN00715156 |
| PRJEB30046 | Shenzen (China) | SAMN00715156 |
| PRJEB30046 | Shenzen (China) | SAMN00715148 |
| PRJEB30046 | Shenzen (China) | SAMN00715137 |
| PRJEB30046 | Shenzen (China) | SAMN00715136 |
| PRJEB30046 | Shenzen (China) | SAMN00715145 |
| PRJEB30046 | Shenzen (China) | SAMN00715139 |
| PRJEB30046 | Shenzen (China) | SAMN00715143 |
| PRJEB30046 | Shenzen (China) | SAMN00715143 |
| PRJEB30046 | Shenzen (China) | SAMN00715134 |
| PRJEB30046 | Shenzen (China) | SAMN00792025 |
| PRJEB30046 | Shenzen (China) | SAMN00792038 |
| PRJEB30046 | Shenzen (China) | SAMN00792008 |
| PRJEB30046 | Shenzen (China) | SAMN00791959 |
| PRJEB30046 | Shenzen (China) | SAMN00791969 |
| PRJEB30046 | Shenzen (China) | SAMN00792036 |
| PRJEB30046 | Shenzen (China) | SAMN00791998 |
| PRJEB30046 | Shenzen (China) | SAMN00792026 |
| PRJEB30046 | Shenzen (China) | SAMN00791946 |
| PRJEB30046 | Shenzen (China) | SAMN00792016 |
| PRJEB30046 | Shenzen (China) | SAMN00792011 |
| PRJEB30046 | Shenzen (China) | SAMN00791961 |
| PRJEB30046 | Shenzen (China) | SAMN00792029 |
| PRJEB30046 | Shenzen (China) | SAMN00792043 |
| PRJEB30046 | Shenzen (China) | SAMN00791953 |
| PRJEB30046 | Shenzen (China) | SAMN00792017 |
| PRJEB30046 | Shenzen (China) | SAMN00792035 |
| PRJEB30046 | Shenzen (China) | SAMN00792007 |
| PRJEB30046 | Shenzen (China) | SAMN00792042 |
| PRJEB30046 | Shenzen (China) | SAMN00792031 |
| PRJEB30046 | Shenzen (China) | SAMN00792031 |
| PRJEB30046 | Shenzen (China) | SAMN00792022 |
| PRJEB30046 | Shenzen (China) | SAMN00791990 |
| PRJEB30046 | Shenzen (China) | SAMN00791984 |

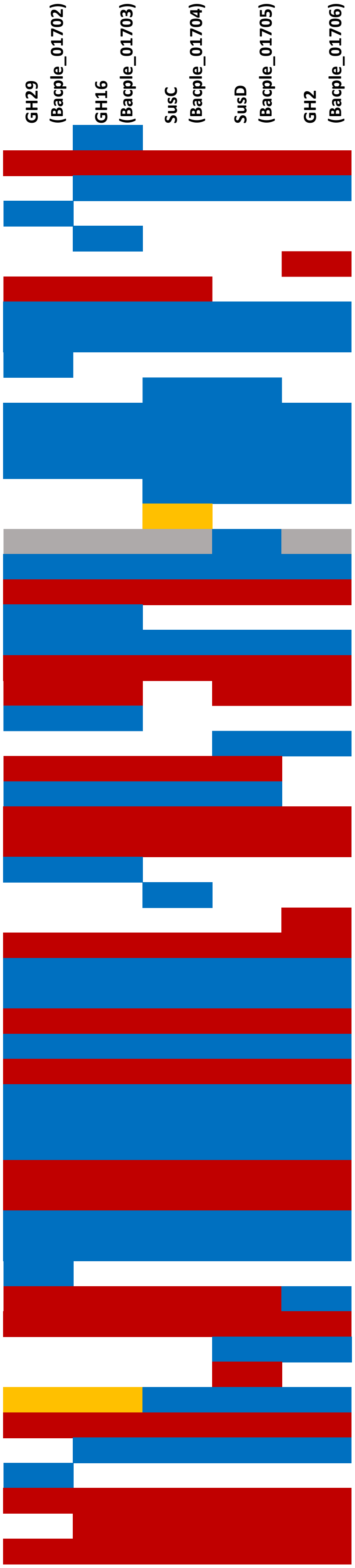

| Study | Location | Biosample |
|---|---|---|
| PRJEB30046 | Shenzen (China) | SAMN00792028 |
| PRJEB30046 | Shenzen (China) | SAMN00791967 |
| PRJEB30046 | Shenzen (China) | SAMN00791968 |
| PRJEB30046 | Shenzen (China) | SAMN00792020 |
| PRJEB30046 | Shenzen (China) | SAMN00792039 |
| PRJEB30046 | Shenzen (China) | SAMN00791952 |
| PRJEB30046 | Shenzen (China) | SAMN00792027 |
| PRJEB30046 | Shenzen (China) | SAMN00792027 |
| PRJEB30046 | Shenzen (China) | SAMN00792037 |
| PRJEB30046 | Shenzen (China) | SAMN00715258 |
| PRJEB30046 | Shenzen (China) | SAMN00715271 |
| PRJEB30046 | Shenzen (China) | SAMN00715275 |
| PRJEB30046 | Shenzen (China) | SAMN00715273 |
| PRJEB30046 | Shenzen (China) | SAMN00715259 |
| PRJEB30046 | Shenzen (China) | SAMN00715256 |
| PRJEB30046 | Shenzen (China) | SAMN00715266 |
| PRJEB30046 | Shenzen (China) | SAMN00715264 |
| PRJEB30046 | Shenzen (China) | SAMN00715264 |
| PRJEB30046 | Shenzen (China) | SAMN00715272 |
| PRJEB30046 | Shenzen (China) | SAMN00715269 |
| PRJEB30046 | Shenzen (China) | SAMN00715255 |
| PRJEB30046 | Shenzen (China) | SAMN00715257 |
| PRJEB30046 | Shenzen (China) | SAMN00792081 |
| PRJEB30046 | Shenzen (China) | SAMN00792091 |
| PRJEB30046 | Shenzen (China) | SAMN00792045 |
| PRJEB30046 | Shenzen (China) | SAMN00792045 |
| PRJEB30046 | Shenzen (China) | SAMN00792098 |
| PRJEB30046 | Shenzen (China) | SAMN00792111 |
| PRJEB30046 | Shenzen (China) | SAMN00792069 |
| PRJEB30046 | Shenzen (China) | SAMN00792069 |
| PRJEB30046 | Shenzen (China) | SAMN00792092 |
| PRJEB30046 | Shenzen (China) | SAMN00792092 |
| PRJEB30046 | Shenzen (China) | SAMN00792085 |
| PRJEB30046 | Shenzen (China) | SAMN00792085 |
| PRJEB30046 | Shenzen (China) | SAMN00792115 |
| PRJEB30046 | Shenzen (China) | SAMN00792062 |
| PRJEB30046 | Shenzen (China) | SAMN00792107 |
| PRJEB30046 | Shenzen (China) | SAMN00792073 |
| PRJEB30046 | Shenzen (China) | SAMN00792112 |
| PRJEB30046 | Shenzen (China) | SAMN00792061 |
| PRJEB30046 | Shenzen (China) | SAMN00792048 |
| PRJEB30046 | Shenzen (China) | SAMN00792099 |
| PRJEB30046 | Shenzen (China) | SAMN00792104 |
| PRJEB30046 | Shenzen (China) | SAMN00792058 |
| PRJEB30046 | Shenzen (China) | SAMN00792080 |
| PRJEB30046 | Shenzen (China) | SAMN00792080 |
| PRJEB30046 | Shenzen (China) | SAMN00792084 |
| PRJEB30046 | Shenzen (China) | SAMN00792089 |
| PRJEB30046 | Shenzen (China) | SAMN00792089 |
| PRJEB30046 | Shenzen (China) | SAMN00792093 |
| PRJEB30046 | Shenzen (China) | SAMN00792056 |
| PRJEB30046 | Shenzen (China) | SAMN00792079 |
| PRJEB30046 | Shenzen (China) | SAMN00792047 |
| PRJEB30046 | Shenzen (China) | SAMN00792102 |
| PRJEB30046 | Shenzen (China) | SAMN00792070 |
| PRJEB30046 | Shenzen (China) | SAMN00792053 |
| PRJEB30046 | Shenzen (China) | SAMN00792057 |

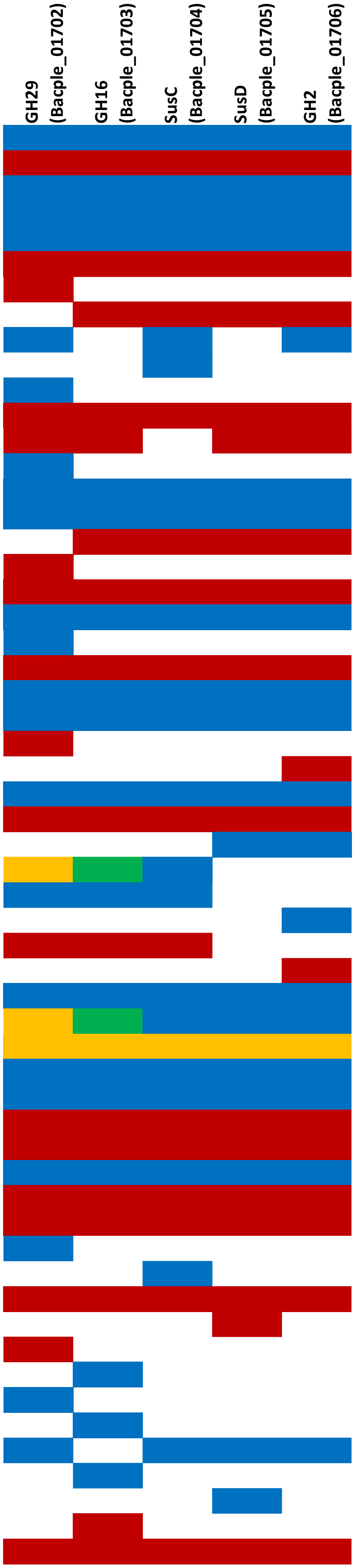

| Study | Location | Biosample |
|---|---|---|
| PRJEB30046 | Shenzen (China) | SAMN00792055 |
| PRJEB30046 | Shenzen (China) | SAMN00792067 |
| PRJEB30046 | Shenzen (China) | SAMN00993245 |
| PRJEB30046 | Shenzen (China) | SAMN00993245 |
| PRJEB30046 | Shenzen (China) | SAMN00993243 |
| PRJEB26092 | Tokyo (Japan) | SAMD00036237 |
| PRJEB26092 | Tokyo (Japan) | SAMD00036296 |
| PRJEB26092 | Tokyo (Japan) | SAMD00036296 |
| PRJEB26092 | Tokyo (Japan) | SAMD00036241 |
| PRJEB26092 | Tokyo (Japan) | SAMD00036241 |
| PRJEB26092 | Tokyo (Japan) | SAMD00036237 |
| PRJEB26092 | Tokyo (Japan) | SAMD00036298 |
| PRJEB26092 | Tokyo (Japan) | SAMD00036284 |
| PRJEB26092 | Tokyo (Japan) | SAMD00036244 |
| PRJEB26092 | Tokyo (Japan) | SAMD00036252 |
| PRJEB26092 | Tokyo (Japan) | SAMD00036252 |
| PRJEB26092 | Tokyo (Japan) | SAMD00036252 |
| PRJEB26092 | Tokyo (Japan) | SAMD00036244 |
| PRJEB26092 | Tokyo (Japan) | SAMD00036244 |
| PRJEB26092 | Tokyo (Japan) | SAMD00036252 |
| PRJEB26092 | Tokyo (Japan) | SAMD00036242 |
| PRJEB26092 | Tokyo (Japan) | SAMD00036242 |
| PRJEB26092 | Tokyo (Japan) | SAMD00036238 |
| PRJEB26092 | Tokyo (Japan) | SAMD00036230 |
| PRJEB26092 | Tokyo (Japan) | SAMD00036296 |
| PRJEB26092 | Tokyo (Japan) | SAMD00036296 |
| PRJEB26092 | Tokyo (Japan) | SAMD00036297 |
| PRJEB26092 | Tokyo (Japan) | SAMD00036297 |
| PRJEB26092 | Tokyo (Japan) | SAMD00036305 |
| PRJEB26092 | Tokyo (Japan) | SAMD00036320 |
| PRJEB26092 | Tokyo (Japan) | SAMD00036344 |
| PRJEB26092 | Tokyo (Japan) | SAMD00036337 |
| PRJEB26092 | Tokyo (Japan) | SAMD00036308 |
| PRJEB26092 | Tokyo (Japan) | SAMD00036308 |
| PRJEB26092 | Tokyo (Japan) | SAMD00036328 |
| PRJEB26092 | Tokyo (Japan) | SAMD00036320 |
| PRJEB26092 | Tokyo (Japan) | SAMD00036204 |
| PRJEB26092 | Tokyo (Japan) | SAMD00036202 |
| PRJEB26092 | Tokyo (Japan) | SAMD00036197 |
| PRJEB26092 | Tokyo (Japan) | SAMD00036197 |
| PRJEB26092 | Tokyo (Japan) | SAMD00036213 |
| PRJEB26092 | Tokyo (Japan) | SAMD00036204 |
| PRJEB26092 | Tokyo (Japan) | SAMD00036192 |
| PRJEB26092 | Tokyo (Japan) | SAMD00036210 |
| PRJEB26092 | Tokyo (Japan) | SAMD00036195 |
| PRJEB26092 | Tokyo (Japan) | SAMD00036195 |
| PRJEB26092 | Tokyo (Japan) | SAMD00036215 |
| PRJEB26092 | Tokyo (Japan) | SAMD00036197 |
| PRJEB26092 | Tokyo (Japan) | SAMD00036215 |
| PRJEB26092 | Tokyo (Japan) | SAMD00036197 |
| PRJEB26092 | Tokyo (Japan) | SAMD00036197 |
| PRJEB26092 | Tokyo (Japan) | SAMD00036441 |
| PRJEB26092 | Tokyo (Japan) | SAMD00036203 |
| PRJEB26092 | Tokyo (Japan) | SAMD00036203 |
| PRJEB26092 | Tokyo (Japan) | SAMD00036214 |
| PRJEB26092 | Tokyo (Japan) | SAMD00036217 |
| PRJEB26092 | Tokyo (Japan) | SAMD00036217 |

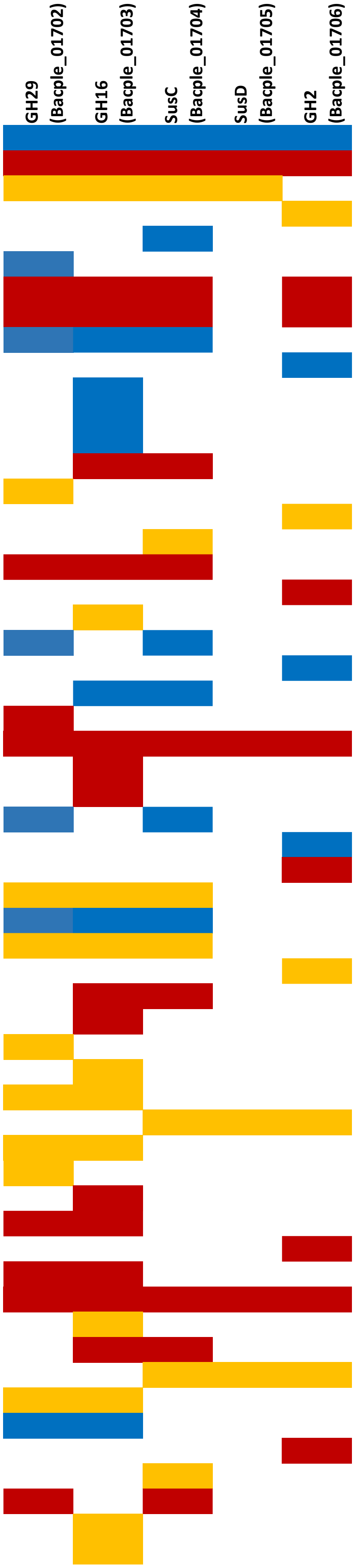

| Study | Location | Biosample | GH29 (Bacple_01702) | GH16 (Bacple_01703) | SusC (Bacple_01704) | SusD (Bacple_01705) | GH2 (Bacple_01706) |
|---|---|---|---|---|---|---|---|
| PRJEB26092 | Tokyo (Japan) | SAMD00036215 | | | | | |
| PRJEB26092 | Tokyo (Japan) | SAMD00036192 | | | | | |
| PRJEB26092 | Tokyo (Japan) | SAMD00036210 | | | | | |
| PRJEB26092 | Tokyo (Japan) | SAMD00036215 | | | | | |
| PRJEB26092 | Tokyo (Japan) | SAMD00036192 | | | | | |
| PRJEB26092 | Tokyo (Japan) | SAMD00036441 | | | | | |
| PRJEB26092 | Tokyo (Japan) | SAMD00036441 | | | | | |
| PRJEB26092 | Tokyo (Japan) | SAMD00036436 | | | | | |
| PRJEB26092 | Tokyo (Japan) | SAMD00036436 | | | | | |
| PRJEB26092 | Tokyo (Japan) | SAMD00036214 | | | | | |
| PRJEB26092 | Tokyo (Japan) | SAMD00036352 | | | | | |
| PRJEB26092 | Tokyo (Japan) | SAMD00036359 | | | | | |
| PRJEB26092 | Tokyo (Japan) | SAMD00036354 | | | | | |
| PRJEB26092 | Tokyo (Japan) | SAMD00036354 | | | | | |
| PRJEB26092 | Tokyo (Japan) | SAMD00036367 | | | | | |
| PRJEB26092 | Tokyo (Japan) | SAMD00036359 | | | | | |
| PRJEB26092 | Tokyo (Japan) | SAMD00036350 | | | | | |
| PRJEB26092 | Tokyo (Japan) | SAMD00036354 | | | | | |
| PRJEB26092 | Tokyo (Japan) | SAMD00036349 | | | | | |
| PRJEB26092 | Tokyo (Japan) | SAMD00036354 | | | | | |
| PRJEB26092 | Tokyo (Japan) | SAMD00036354 | | | | | |
| PRJEB26092 | Tokyo (Japan) | SAMD00036409 | | | | | |
| PRJEB26092 | Tokyo (Japan) | SAMD00036409 | | | | | |
| PRJEB26092 | Tokyo (Japan) | SAMD00036419 | | | | | |
| PRJEB26092 | Tokyo (Japan) | SAMD00036419 | | | | | |
| PRJEB26092 | Tokyo (Japan) | SAMD00036349 | | | | | |
| PRJEB26092 | Tokyo (Japan) | SAMD00036349 | | | | | |
| PRJEB26092 | Tokyo (Japan) | SAMD00036388 | | | | | |
| PRJEB26092 | Tokyo (Japan) | SAMD00036412 | | | | | |
| PRJEB26092 | Tokyo (Japan) | SAMD00036404 | | | | | |
| PRJEB26092 | Tokyo (Japan) | SAMD00036404 | | | | | |
| PRJEB26092 | Tokyo (Japan) | SAMD00036404 | | | | | |
| PRJEB26092 | Tokyo (Japan) | SAMD00036404 | | | | | |
| PRJEB26092 | Tokyo (Japan) | SAMD00036400 | | | | | |
| PRJEB26092 | Tokyo (Japan) | SAMD00036392 | | | | | |
| PRJEB26092 | Tokyo (Japan) | SAMD00036388 | | | | | |
| PRJEB26092 | Tokyo (Japan) | SAMD00036388 | | | | | |
| PRJEB26092 | Tokyo (Japan) | SAMD00036388 | | | | | |
| PRJEB26092 | Tokyo (Japan) | SAMD00036413 | | | | | |
| PRJEB26092 | Tokyo (Japan) | SAMD00036405 | | | | | |
| PRJEB26092 | Tokyo (Japan) | SAMD00036405 | | | | | |
| PRJEB26092 | Tokyo (Japan) | SAMD00036379 | | | | | |
| PRJEB26092 | Tokyo (Japan) | SAMD00036398 | | | | | |
| PRJEB26092 | Tokyo (Japan) | SAMD00036432 | | | | | |
| PRJEB26092 | Tokyo (Japan) | SAMD00036377 | | | | | |
| PRJEB26092 | Tokyo (Japan) | SAMD00036388 | | | | | |
| PRJEB26092 | Tokyo (Japan) | SAMD00036386 | | | | | |
| PRJEB26092 | Tokyo (Japan) | SAMD00036400 | | | | | |
| PRJEB26092 | Tokyo (Japan) | SAMD00036386 | | | | | |
| PRJEB26092 | Tokyo (Japan) | SAMD00036379 | | | | | |
| PRJEB26092 | Tokyo (Japan) | SAMD00036377 | | | | | |
| PRJEB26092 | Tokyo (Japan) | SAMD00036404 | | | | | |
| PRJEB26092 | Tokyo (Japan) | SAMD00036432 | | | | | |
| PRJDB4176 | Tokyo (Japan) | SAMD00114727 | | | | | |
| PRJDB4176 | Tokyo (Japan) | SAMD00114731 | | | | | |
| PRJDB4176 | Tokyo (Japan) | SAMD00114734 | | | | | |
| PRJDB4176 | Tokyo (Japan) | SAMD00114737 | | | | | |

| Study | Location | Biosample |
|---|---|---|
| PRJDB4176 | Tokyo (Japan) | SAMD00114760 |
| PRJDB4176 | Tokyo (Japan) | SAMD00114798 |
| PRJDB4176 | Tokyo (Japan) | SAMD00114805 |
| PRJDB4176 | Tokyo (Japan) | SAMD00114805 |
| PRJDB4176 | Tokyo (Japan) | SAMD00114805 |
| PRJDB4176 | Tokyo (Japan) | SAMD00114825 |
| PRJDB4176 | Tokyo (Japan) | SAMD00114825 |
| PRJDB4176 | Tokyo (Japan) | SAMD00114829 |
| PRJDB4176 | Tokyo (Japan) | SAMD00114834 |
| PRJDB4176 | Tokyo (Japan) | SAMD00114834 |
| PRJDB4176 | Tokyo (Japan) | SAMD00114865 |
| PRJDB4176 | Tokyo (Japan) | SAMD00114871 |
| PRJDB4176 | Tokyo (Japan) | SAMD00114892 |
| PRJDB4176 | Tokyo (Japan) | SAMD00114893 |
| PRJDB4176 | Tokyo (Japan) | SAMD00114895 |
| PRJDB4176 | Tokyo (Japan) | SAMD00114953 |
| PRJDB4176 | Tokyo (Japan) | SAMD00114954 |
| PRJDB4176 | Tokyo (Japan) | SAMD00114966 |
| PRJDB4176 | Tokyo (Japan) | SAMD00114977 |
| PRJDB4176 | Tokyo (Japan) | SAMD00115001 |
| PRJDB4176 | Tokyo (Japan) | SAMD00115023 |
| PRJDB4176 | Tokyo (Japan) | SAMD00115023 |
| PRJDB4176 | Tokyo (Japan) | SAMD00115023 |
| PRJDB4176 | Tokyo (Japan) | SAMD00154995 |
| PRJDB4176 | Tokyo (Japan) | SAMD00164695 |
| PRJDB4176 | Tokyo (Japan) | SAMD00164713 |
| PRJDB4176 | Tokyo (Japan) | SAMD00164721 |
| PRJDB4176 | Tokyo (Japan) | SAMD00164755 |
| PRJDB4176 | Tokyo (Japan) | SAMD00164756 |
| PRJDB4176 | Tokyo (Japan) | SAMD00164765 |
| PRJDB4176 | Tokyo (Japan) | SAMD00164767 |
| PRJDB4176 | Tokyo (Japan) | SAMD00164769 |
| PRJDB4176 | Tokyo (Japan) | SAMD00164778 |
| PRJDB4176 | Tokyo (Japan) | SAMD00164780 |
| PRJDB4176 | Tokyo (Japan) | SAMD00164781 |
| PRJDB4176 | Tokyo (Japan) | SAMD00164805 |
| PRJDB4176 | Tokyo (Japan) | SAMD00164808 |
| PRJDB4176 | Tokyo (Japan) | SAMD00164817 |
| PRJDB4176 | Tokyo (Japan) | SAMD00164817 |
| PRJDB4176 | Tokyo (Japan) | SAMD00164817 |
| PRJDB4176 | Tokyo (Japan) | SAMD00164818 |
| PRJDB4176 | Tokyo (Japan) | SAMD00164819 |
| PRJDB4176 | Tokyo (Japan) | SAMD00164820 |
| PRJDB4176 | Tokyo (Japan) | SAMD00164824 |
| PRJDB4176 | Tokyo (Japan) | SAMD00164829 |
| PRJDB4176 | Tokyo (Japan) | SAMD00164830 |
| PRJDB4176 | Tokyo (Japan) | SAMD00164832 |
| PRJDB4176 | Tokyo (Japan) | SAMD00164833 |
| PRJDB4176 | Tokyo (Japan) | SAMD00164833 |
| PRJDB4176 | Tokyo (Japan) | SAMD00164834 |
| PRJDB4176 | Tokyo (Japan) | SAMD00164835 |
| PRJDB4176 | Tokyo (Japan) | SAMD00164841 |
| PRJDB4176 | Tokyo (Japan) | SAMD00164841 |
| PRJDB4176 | Tokyo (Japan) | SAMD00164841 |
| PRJDB4176 | Tokyo (Japan) | SAMD00164849 |
| PRJDB4176 | Tokyo (Japan) | SAMD00164853 |
| PRJDB4176 | Tokyo (Japan) | SAMD00164856 |

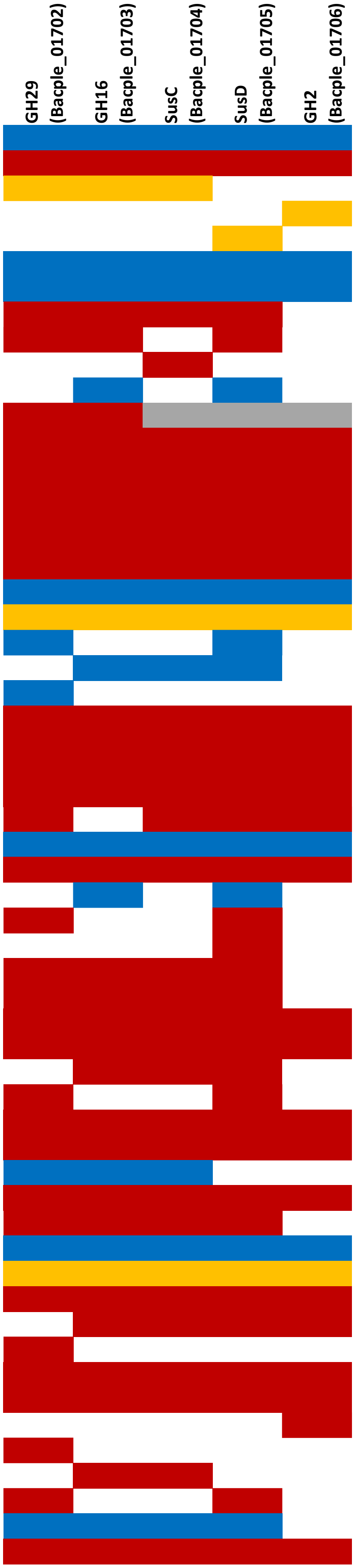

| Study | Location | Biosample | GH29 (Bacple_01702) | GH16 (Bacple_01703) | SusC (Bacple_01704) | SusD (Bacple_01705) | GH2 (Bacple_01706) |
|---|---|---|---|---|---|---|---|
| PRJDB4176 | Tokyo (Japan) | SAMD00164860 | | | | | |
| PRJDB4176 | Tokyo (Japan) | SAMD00164860 | | | | | |
| PRJDB4176 | Tokyo (Japan) | SAMD00164863 | | | | | |
| PRJDB4176 | Tokyo (Japan) | SAMD00164867 | | | | | |
| PRJDB4176 | Tokyo (Japan) | SAMD00164867 | | | | | |
| PRJDB4176 | Tokyo (Japan) | SAMD00164869 | | | | | |
| PRJDB4176 | Tokyo (Japan) | SAMD00164872 | | | | | |
| PRJDB4176 | Tokyo (Japan) | SAMD00164874 | | | | | |
| PRJDB4176 | Tokyo (Japan) | SAMD00164888 | | | | | |
| PRJDB4176 | Tokyo (Japan) | SAMD00164889 | | | | | |
| PRJDB4176 | Tokyo (Japan) | SAMD00164894 | | | | | |
| PRJDB4176 | Tokyo (Japan) | SAMD00164895 | | | | | |
| PRJDB4176 | Tokyo (Japan) | SAMD00164895 | | | | | |
| PRJDB4176 | Tokyo (Japan) | SAMD00164897 | | | | | |
| PRJDB4176 | Tokyo (Japan) | SAMD00164898 | | | | | |
| PRJDB4176 | Tokyo (Japan) | SAMD00164900 | | | | | |
| PRJDB4176 | Tokyo (Japan) | SAMD00164901 | | | | | |
| PRJDB4176 | Tokyo (Japan) | SAMD00164915 | | | | | |
| PRJDB4176 | Tokyo (Japan) | SAMD00164916 | | | | | |
| PRJDB4176 | Tokyo (Japan) | SAMD00164921 | | | | | |
| PRJDB4176 | Tokyo (Japan) | SAMD00164922 | | | | | |
| PRJDB4176 | Tokyo (Japan) | SAMD00164924 | | | | | |
| PRJDB4176 | Tokyo (Japan) | SAMD00164925 | | | | | |
| PRJDB4176 | Tokyo (Japan) | SAMD00164928 | | | | | |
| PRJDB4176 | Tokyo (Japan) | SAMD00164942 | | | | | |
| PRJDB4176 | Tokyo (Japan) | SAMD00164945 | | | | | |
| PRJDB4176 | Tokyo (Japan) | SAMD00164946 | | | | | |
| PRJDB4176 | Tokyo (Japan) | SAMD00164948 | | | | | |
| PRJDB4176 | Tokyo (Japan) | SAMD00164950 | | | | | |
| PRJDB4176 | Tokyo (Japan) | SAMD00164954 | | | | | |
| PRJDB4176 | Tokyo (Japan) | SAMD00164967 | | | | | |
| PRJDB4176 | Tokyo (Japan) | SAMD00164967 | | | | | |
| PRJDB4176 | Tokyo (Japan) | SAMD00164976 | | | | | |
| PRJDB4176 | Tokyo (Japan) | SAMD00164977 | | | | | |
| PRJDB4176 | Tokyo (Japan) | SAMD00164984 | | | | | |
| PRJDB4176 | Tokyo (Japan) | SAMD00164987 | | | | | |
| PRJDB4176 | Tokyo (Japan) | SAMD00164991 | | | | | |
| PRJDB4176 | Tokyo (Japan) | SAMD00164995 | | | | | |
| PRJDB4176 | Tokyo (Japan) | SAMD00164995 | | | | | |
| PRJDB4176 | Tokyo (Japan) | SAMD00164998 | | | | | |
| PRJDB4176 | Tokyo (Japan) | SAMD00164999 | | | | | |
| PRJDB4176 | Tokyo (Japan) | SAMD00165000 | | | | | |
| PRJDB4176 | Tokyo (Japan) | SAMD00165002 | | | | | |
| PRJDB4176 | Tokyo (Japan) | SAMD00165004 | | | | | |
| PRJDB4176 | Tokyo (Japan) | SAMD00165005 | | | | | |
| PRJDB4176 | Tokyo (Japan) | SAMD00165015 | | | | | |
| PRJDB4176 | Tokyo (Japan) | SAMD00165032 | | | | | |
| PRJDB4176 | Tokyo (Japan) | SAMD00114718 | | | | | |
| PRJDB4176 | Tokyo (Japan) | SAMD00114719 | | | | | |
| PRJDB4176 | Tokyo (Japan) | SAMD00114719 | | | | | |
| PRJDB4176 | Tokyo (Japan) | SAMD00114721 | | | | | |
| PRJDB4176 | Tokyo (Japan) | SAMD00114729 | | | | | |
| PRJDB4176 | Tokyo (Japan) | SAMD00114738 | | | | | |
| PRJDB4176 | Tokyo (Japan) | SAMD00114740 | | | | | |
| PRJDB4176 | Tokyo (Japan) | SAMD00114742 | | | | | |
| PRJDB4176 | Tokyo (Japan) | SAMD00114756 | | | | | |
| PRJDB4176 | Tokyo (Japan) | SAMD00114758 | | | | | |

| Study | Location | Biosample |
|---|---|---|
| PRJDB4176 | Tokyo (Japan) | SAMD00114768 |
| PRJDB4176 | Tokyo (Japan) | SAMD00114771 |
| PRJDB4176 | Tokyo (Japan) | SAMD00114779 |
| PRJDB4176 | Tokyo (Japan) | SAMD00114788 |
| PRJDB4176 | Tokyo (Japan) | SAMD00114799 |
| PRJDB4176 | Tokyo (Japan) | SAMD00114799 |
| PRJDB4176 | Tokyo (Japan) | SAMD00114800 |
| PRJDB4176 | Tokyo (Japan) | SAMD00114802 |
| PRJDB4176 | Tokyo (Japan) | SAMD00114804 |
| PRJDB4176 | Tokyo (Japan) | SAMD00114806 |
| PRJDB4176 | Tokyo (Japan) | SAMD00114806 |
| PRJDB4176 | Tokyo (Japan) | SAMD00114807 |
| PRJDB4176 | Tokyo (Japan) | SAMD00114810 |
| PRJDB4176 | Tokyo (Japan) | SAMD00114816 |
| PRJDB4176 | Tokyo (Japan) | SAMD00114819 |
| PRJDB4176 | Tokyo (Japan) | SAMD00114824 |
| PRJDB4176 | Tokyo (Japan) | SAMD00114828 |
| PRJDB4176 | Tokyo (Japan) | SAMD00114833 |
| PRJDB4176 | Tokyo (Japan) | SAMD00114847 |
| PRJDB4176 | Tokyo (Japan) | SAMD00114851 |
| PRJDB4176 | Tokyo (Japan) | SAMD00114853 |
| PRJDB4176 | Tokyo (Japan) | SAMD00114853 |
| PRJDB4176 | Tokyo (Japan) | SAMD00114856 |
| PRJDB4176 | Tokyo (Japan) | SAMD00114870 |
| PRJDB4176 | Tokyo (Japan) | SAMD00114872 |
| PRJDB4176 | Tokyo (Japan) | SAMD00114872 |
| PRJDB4176 | Tokyo (Japan) | SAMD00114877 |
| PRJDB4176 | Tokyo (Japan) | SAMD00114878 |
| PRJDB4176 | Tokyo (Japan) | SAMD00114885 |
| PRJDB4176 | Tokyo (Japan) | SAMD00114885 |
| PRJDB4176 | Tokyo (Japan) | SAMD00114888 |
| PRJDB4176 | Tokyo (Japan) | SAMD00114891 |
| PRJDB4176 | Tokyo (Japan) | SAMD00114896 |
| PRJDB4176 | Tokyo (Japan) | SAMD00114897 |
| PRJDB4176 | Tokyo (Japan) | SAMD00114903 |
| PRJDB4176 | Tokyo (Japan) | SAMD00114904 |
| PRJDB4176 | Tokyo (Japan) | SAMD00114909 |
| PRJDB4176 | Tokyo (Japan) | SAMD00114913 |
| PRJDB4176 | Tokyo (Japan) | SAMD00114919 |
| PRJDB4176 | Tokyo (Japan) | SAMD00114922 |
| PRJDB4176 | Tokyo (Japan) | SAMD00114927 |
| PRJDB4176 | Tokyo (Japan) | SAMD00114927 |
| PRJDB4176 | Tokyo (Japan) | SAMD00114928 |
| PRJDB4176 | Tokyo (Japan) | SAMD00114928 |
| PRJDB4176 | Tokyo (Japan) | SAMD00114939 |
| PRJDB4176 | Tokyo (Japan) | SAMD00114940 |
| PRJDB4176 | Tokyo (Japan) | SAMD00114947 |
| PRJDB4176 | Tokyo (Japan) | SAMD00114961 |
| PRJDB4176 | Tokyo (Japan) | SAMD00114962 |
| PRJDB4176 | Tokyo (Japan) | SAMD00114968 |
| PRJDB4176 | Tokyo (Japan) | SAMD00114981 |
| PRJDB4176 | Tokyo (Japan) | SAMD00114984 |
| PRJDB4176 | Tokyo (Japan) | SAMD00114987 |
| PRJDB4176 | Tokyo (Japan) | SAMD00114989 |
| PRJDB4176 | Tokyo (Japan) | SAMD00114990 |
| PRJDB4176 | Tokyo (Japan) | SAMD00114993 |
| PRJDB4176 | Tokyo (Japan) | SAMD00114999 |

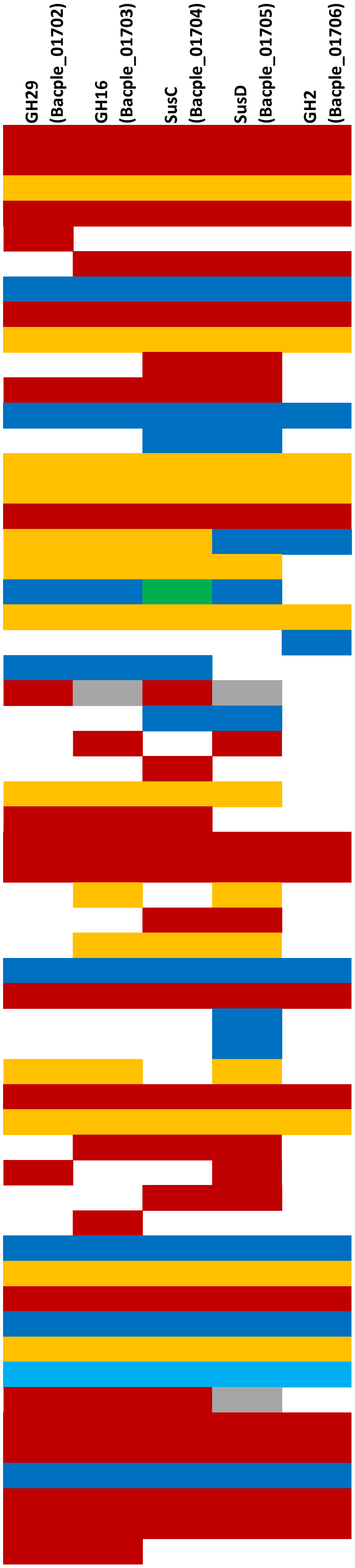

| Study | Location | Biosample |
|---|---|---|
| PRJDB4176 | Tokyo (Japan) | SAMD00114999 |
| PRJDB4176 | Tokyo (Japan) | SAMD00115004 |
| PRJDB4176 | Tokyo (Japan) | SAMD00115005 |
| PRJDB4176 | Tokyo (Japan) | SAMD00115005 |
| PRJDB4176 | Tokyo (Japan) | SAMD00115006 |
| PRJDB4176 | Tokyo (Japan) | SAMD00115006 |
| PRJDB4176 | Tokyo (Japan) | SAMD00115007 |
| PRJDB4176 | Tokyo (Japan) | SAMD00115010 |
| PRJDB4176 | Tokyo (Japan) | SAMD00115015 |
| PRJDB4176 | Tokyo (Japan) | SAMD00115015 |
| PRJDB4176 | Tokyo (Japan) | SAMD00115018 |
| PRJDB4176 | Tokyo (Japan) | SAMD00115024 |
| PRJDB4176 | Tokyo (Japan) | SAMD00115027 |
| PRJDB4176 | Tokyo (Japan) | SAMD00115036 |
| PRJDB4176 | Tokyo (Japan) | SAMD00115045 |
| PRJDB4176 | Tokyo (Japan) | SAMD00115048 |
| PRJDB4176 | Tokyo (Japan) | SAMD00115051 |
| PRJDB4176 | Tokyo (Japan) | SAMD00115053 |
| PRJDB4176 | Tokyo (Japan) | SAMD00115059 |
| PRJDB4176 | Tokyo (Japan) | SAMD00115063 |
| PRJDB4176 | Tokyo (Japan) | SAMD00115068 |
| PRJDB4176 | Tokyo (Japan) | SAMD00115074 |
| PRJDB4176 | Tokyo (Japan) | SAMD00115089 |
| PRJNA678426 | Korea | SAMN16796161 |
| PRJNA678426 | Korea | SAMN16796164 |
| PRJNA678426 | Korea | SAMN16796151 |
| PRJNA678426 | Korea | SAMN16796151 |
| PRJNA678426 | Korea | SAMN16796157 |
| PRJNA678426 | Korea | SAMN16796157 |
| PRJNA678426 | Korea | SAMN16796165 |
| PRJNA678426 | Korea | SAMN16796170 |
| PRJNA678426 | Korea | SAMN16796172 |
| PRJNA678426 | Korea | SAMN16796174 |
| PRJNA678426 | Korea | SAMN16796176 |
| PRJNA678426 | Korea | SAMN16796179 |
| PRJNA678426 | Korea | SAMN16796183 |
| PRJNA678426 | Korea | SAMN16796184 |
| PRJNA678426 | Korea | SAMN16796188 |
| PRJNA678426 | Korea | SAMN16796194 |
| PRJNA678426 | Korea | SAMN16796195 |
| PRJNA678426 | Korea | SAMN16796203 |
| PRJNA678426 | Korea | SAMN16796204 |
| PRJNA678426 | Korea | SAMN16796211 |
| PRJNA678426 | Korea | SAMN16796213 |
| PRJNA678426 | Korea | SAMN16796215 |
| PRJNA678426 | Korea | SAMN16796217 |
| PRJNA678426 | Korea | SAMN16796221 |
| PRJNA678426 | Korea | SAMN16796236 |
| PRJEB26167 | Anhui (China) | SAMN06115447 |
| PRJEB26167 | Anhui (China) | SAMN06115484 |
| PRJEB26167 | Anhui (China) | SAMN06115505 |
| PRJEB26167 | Chongqing (China) | SAMN06115457 |
| PRJEB26167 | Chongqing (China) | SAMN06115459 |
| PRJEB26167 | Chongqing (China) | SAMN06115468 |
| PRJEB26167 | Chongqing (China) | SAMN06115485 |
| PRJEB26167 | Chongqing (China) | SAMN06115492 |
| PRJEB26167 | Fujian (China) | SAMN06115453 |

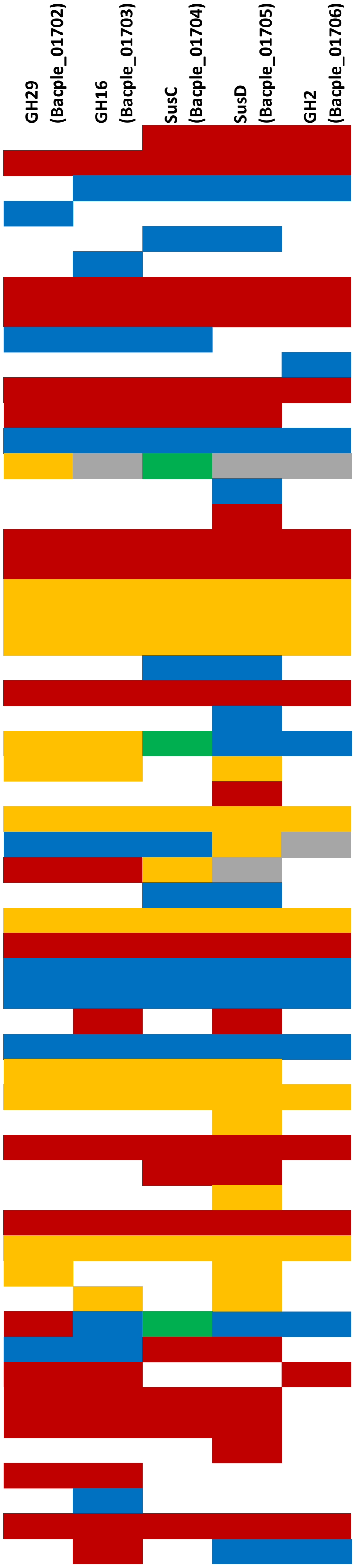

| Study | Location | Biosample | GH29 (Bacple_01702) | GH16 (Bacple_01703) | SusC (Bacple_01704) | SusD (Bacple_01705) | GH2 (Bacple_01706) |
|---|---|---|---|---|---|---|---|
| PRJEB26167 | Gansu (China) | SAMN06115528 | | | | | |
| PRJEB26167 | Guizhou (China) | SAMN06115411 | | | | | |
| PRJEB26167 | Guizhou (China) | SAMN06115416 | | | | | |
| PRJEB26167 | Guizhou (China) | SAMN06115421 | | | | | |
| PRJEB26167 | Guizhou (China) | SAMN06115482 | | | | | |
| PRJEB26167 | Henan (China) | SAMN06115427 | | | | | |
| PRJEB26167 | Henan (China) | SAMN06115433 | | | | | |
| PRJEB26167 | Henan (China) | SAMN06115441 | | | | | |
| PRJEB26167 | Henan (China) | SAMN06115443 | | | | | |
| PRJEB26167 | Henan (China) | SAMN06115455 | | | | | |
| PRJEB26167 | Henan (China) | SAMN06115470 | | | | | |
| PRJEB26167 | Hubei (China) | SAMN06115440 | | | | | |
| PRJEB26167 | Hubei (China) | SAMN06115444 | | | | | |
| PRJEB26167 | Hubei (China) | SAMN06115501 | | | | | |
| PRJEB26167 | Jiangsu (China) | SAMN06115452 | | | | | |
| PRJEB26167 | Jiangsu (China) | SAMN06115474 | | | | | |
| PRJEB26167 | Ningxia (China) | SAMN06115515 | | | | | |
| PRJEB26167 | Shanxi (China) | SAMN06115514 | | | | | |
| PRJEB26167 | Shanxi (China) | SAMN06115539 | | | | | |
| PRJEB26167 | Sichuan (China) | SAMN06115478 | | | | | |
| PRJEB26167 | Sichuan (China) | SAMN06115483 | | | | | |
| PRJEB26167 | Sichuan (China) | SAMN06115510 | | | | | |
| PRJEB26167 | Sichuan (China) | SAMN06115530 | | | | | |
| PRJEB22283 | HMP (USA) | SAMN00034905 | | | | | |
| PRJEB22283 | HMP (USA) | SAMN00037012 | | | | | |
| PRJEB22283 | HMP (USA) | SAMN00038172 | | | | | |
| PRJEB22283 | HMP (USA) | SAMN00045549 | | | | | |
| PRJEB22283 | HMP (USA) | SAMN00045843 | | | | | |
| PRJEB26490 | Lincoln (USA) | SAMN05194702 | | | | | |

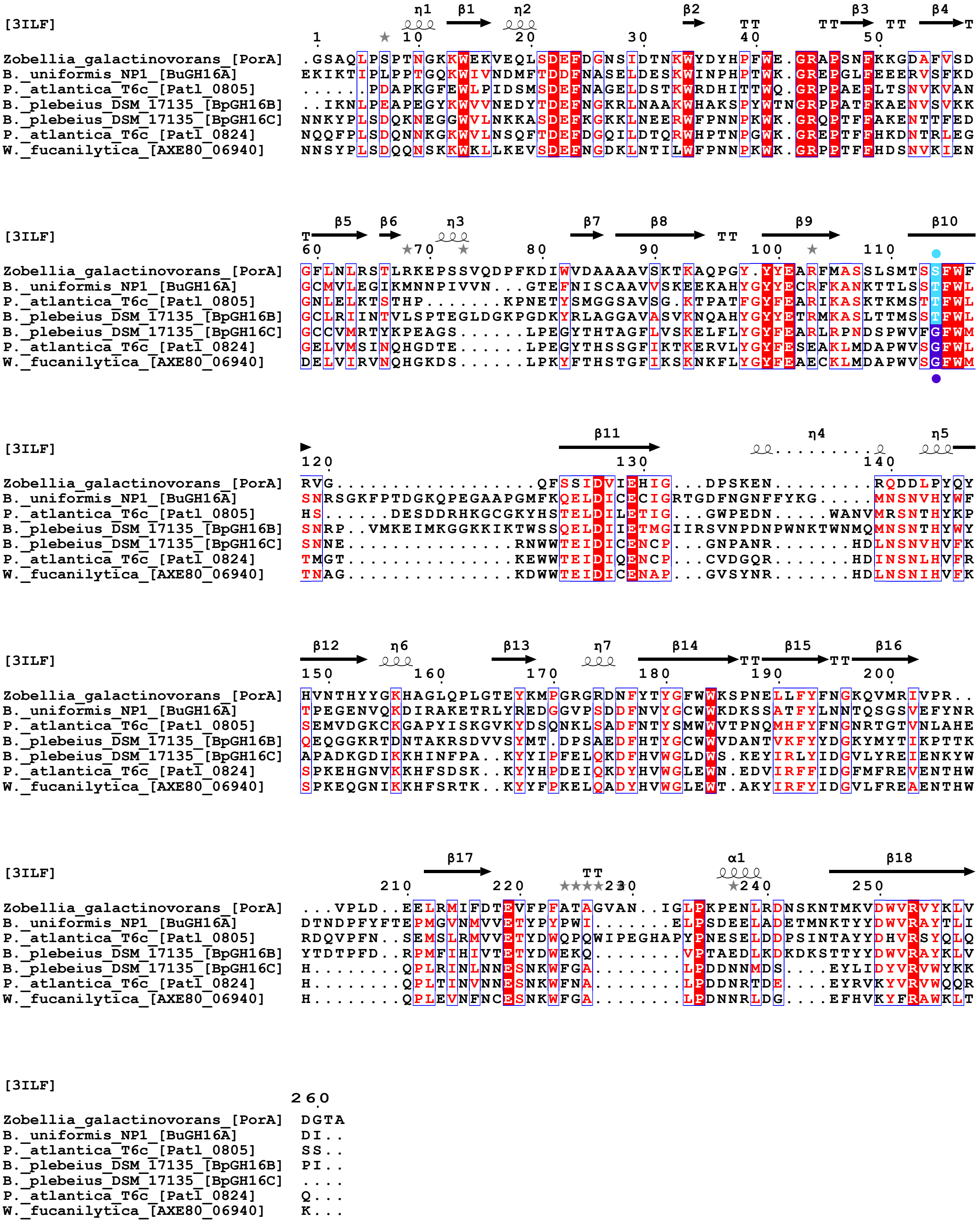


**Figure S1:** Structural alignment of the *B. plebeius* DSM 17135 BpGH16C 6-O- methyl-β-porphyranase compared with the other characterized 6-O-methyl-β-porphyranases (*P. atlantica* T6c, Patl_0824; *W. fucanilytica*, AXE80_06940) and characterized β-porphyranases (*B. uniformis* NP1, BuGH16A; *P. atlantica* T6c, Patl_0805; *B. plebeius* DSM 17135, BpGH16B). The blue circle indicates the serine/threonine amino acids observed in the active site of the β-porphyranases which are replaced by a glycine amino acid in 6-O-methyl-β-porphyranases allowing to accommodate the methyl group.

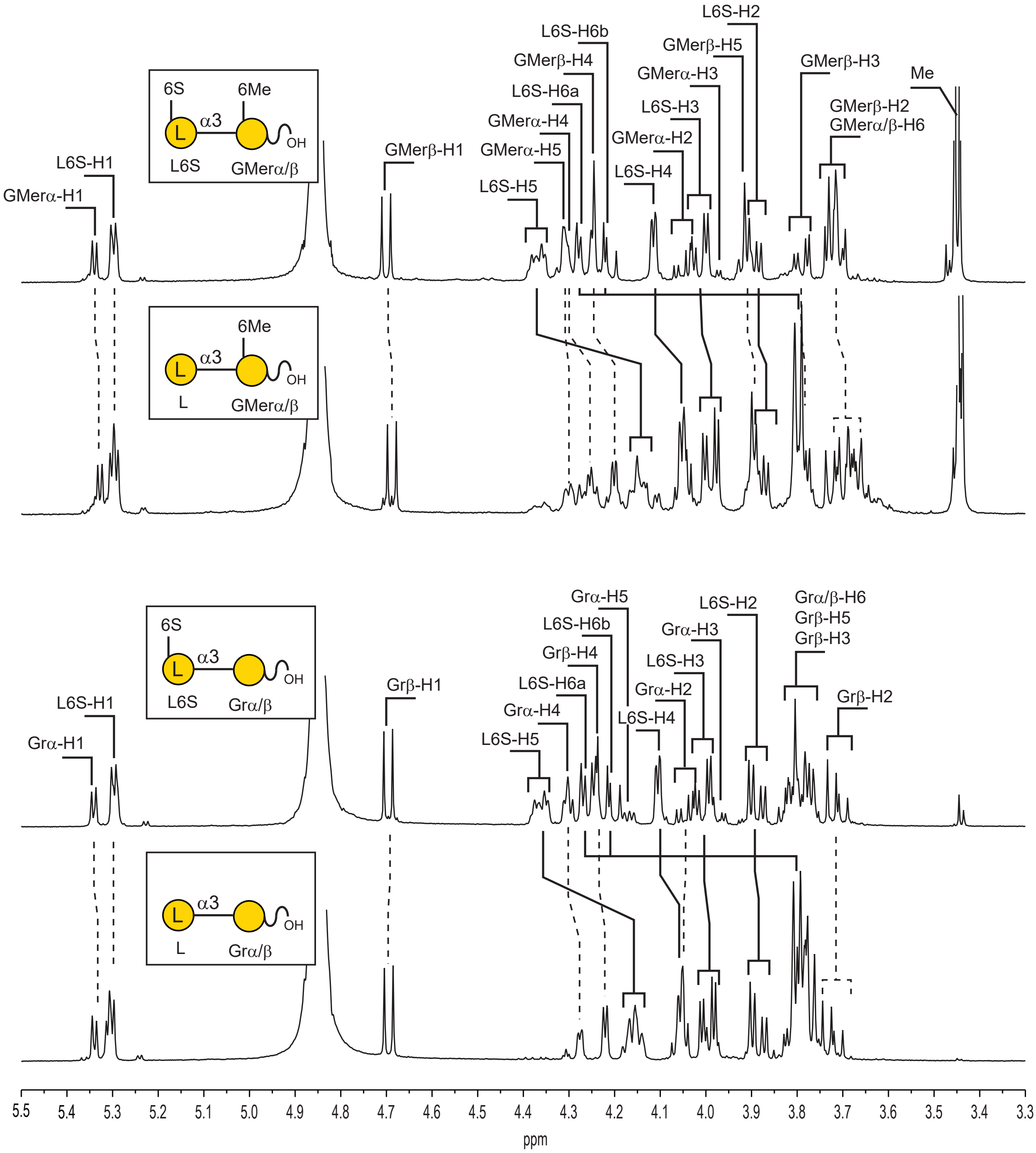


**Figure S2**: [1]H NMR recorded before and after incubation with the sulfatase BpS1_11 on purified disaccharides end-products of the BpGH16C. The occurrence of the methyl group located at the position 6 of the D-galactose didn't hinder the removal of the sulfate ester group of the L-galactose residue located at the non-reducing end.

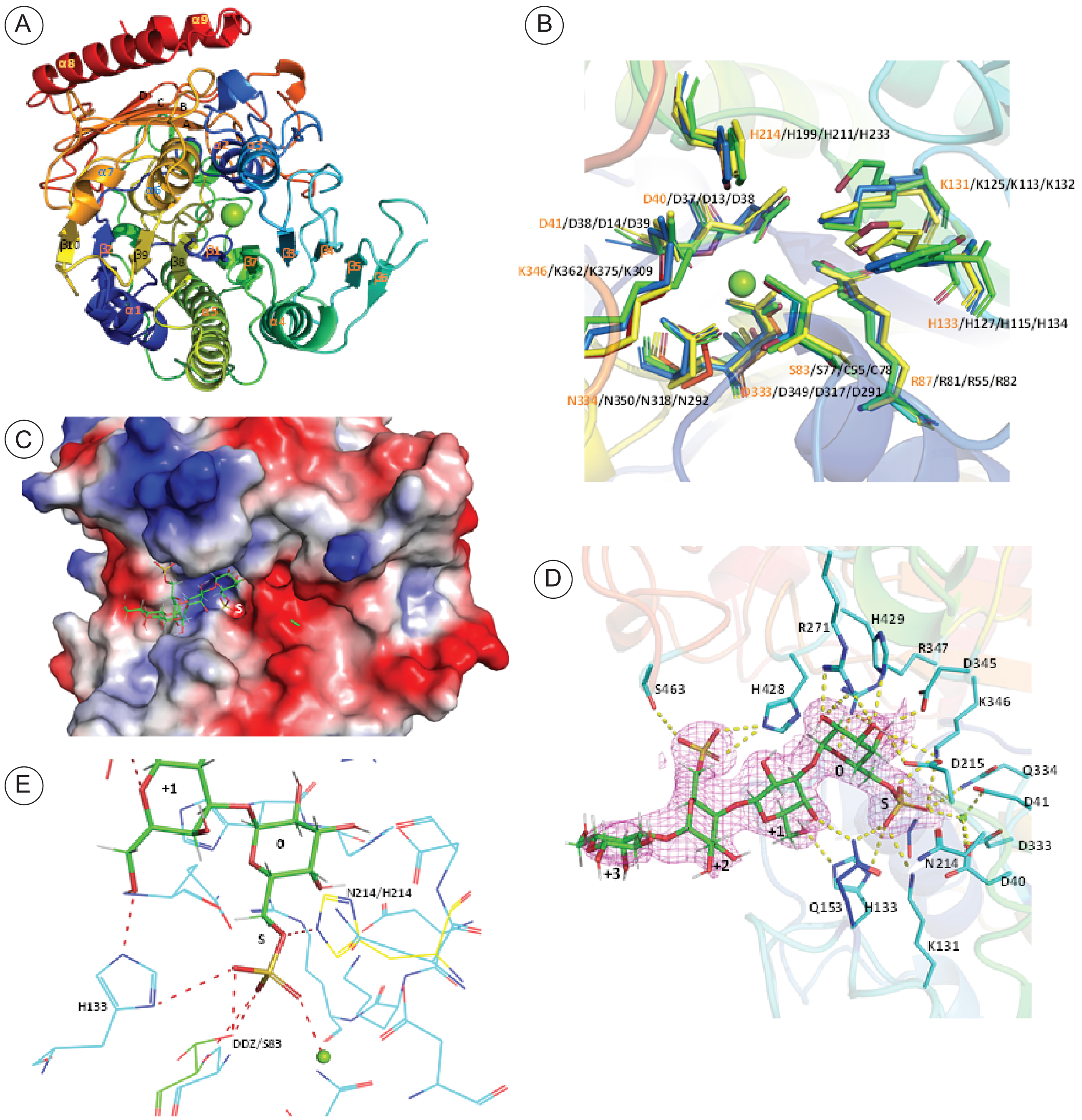


**Figure S3: A)** Structural details of Bacple_01701. Schematic representation of Bacple_01701 shown in cartoon representation and colored from N to C terminus. Calcium ion is shown as green sphere. **B)** Conserved active site residues from S1 family sulfatases. The residues are shown in stick representation in the order Bacple_01701/ 5G2V/1HDH/6BIA. **C)** Surface representation of H214 mutant with the tetrasaccharide, showing pocket like architecture of active site. The 6S sulfate subsite is labelled as S. **D)** Complex structure of H214N mutant. The electron density (2fo-fc) map for the sugar contoured at 1.5σ. Sugars numbered from 0 to +3 and 6S sulfate subsite labelled as S. Residues at H bonding distance to the substrate are shown in stick representation. **E)** Structural comparison of H214 mutant with 1HDH. The residues involved in catalysis S83 (overlays with DDZ from 1HDH), H133, H214 (this is the mutated residue) are shown in stick representation.

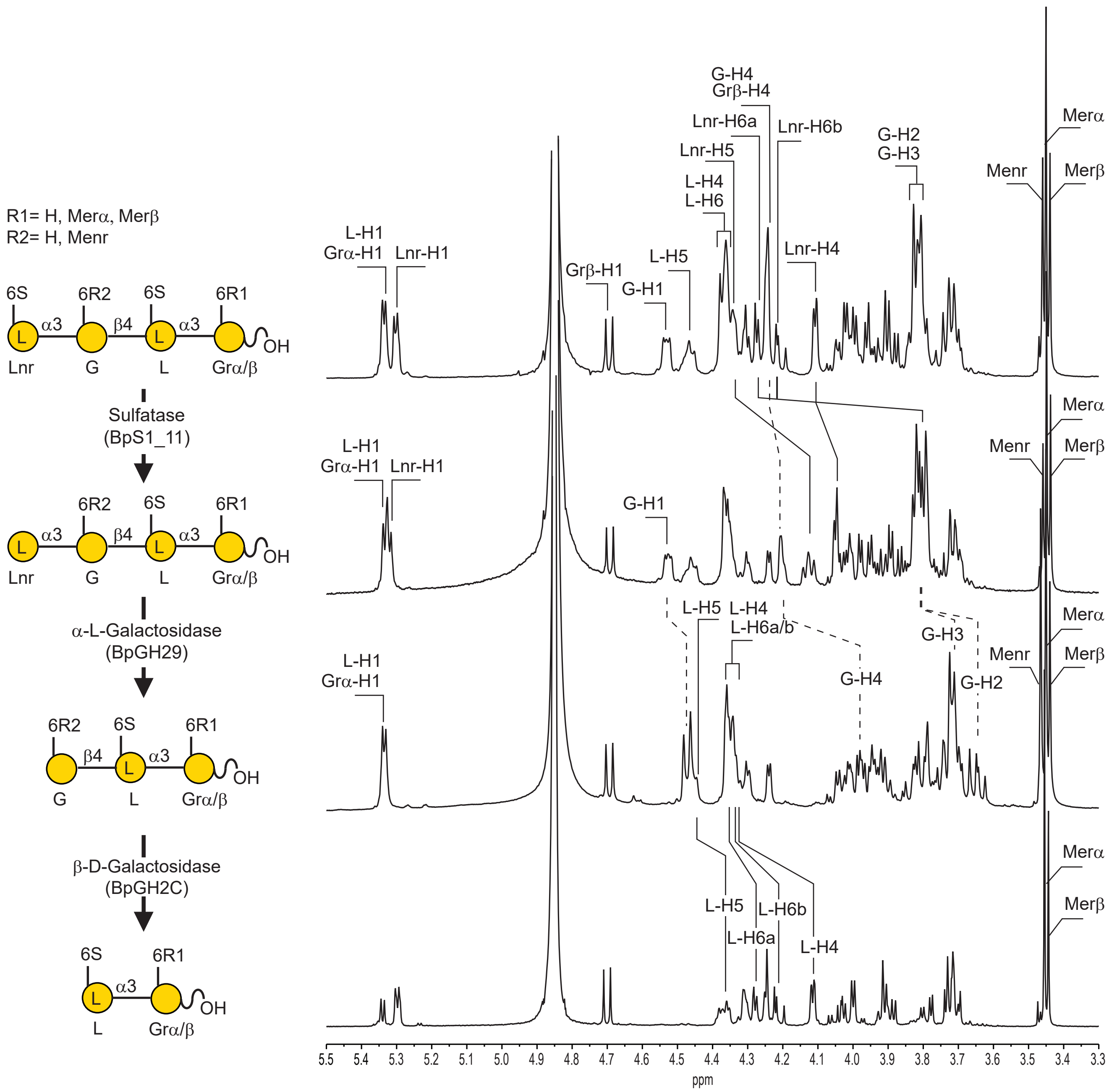


**Figure S4:** [1]H NMR recorded after sequential incubation of the methylated tetrasaccharide end-products of the 6-O-methyl-β-porphyranase BpGH16C with the sulfatase BpS1_11 followed by the β-L-galactosidase BpGH29 and the β-D-galactosidase BpGH2C.

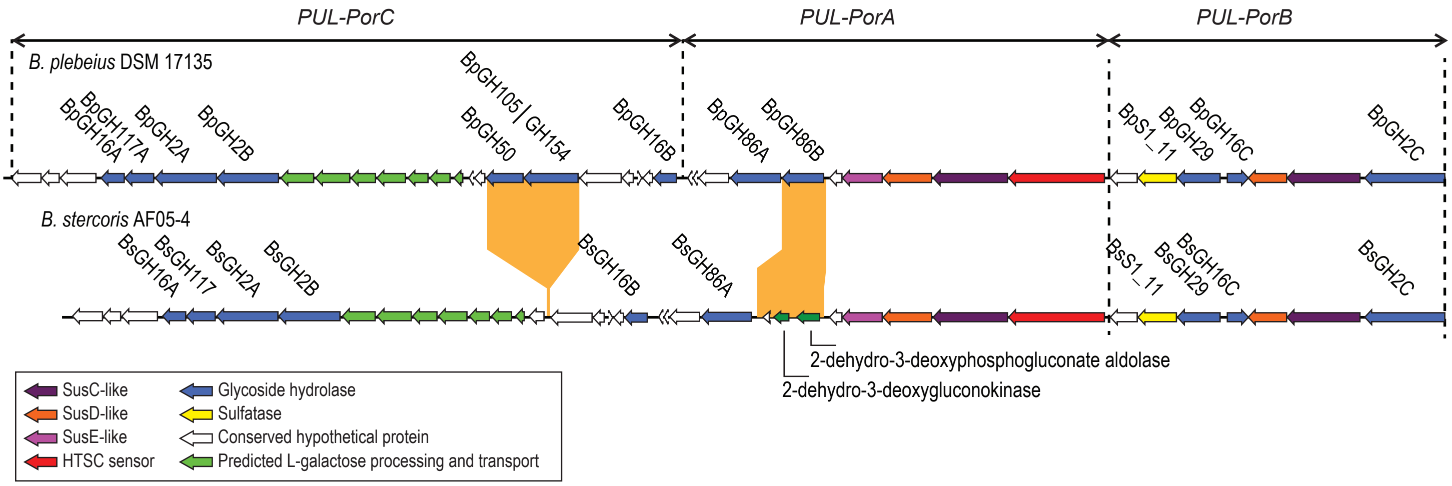


**Figure S5:** Comparison of the two main gene organizations of the porphyran PUL observed in the set of isolated bacteria investigated highlighting deletion and recombination events.

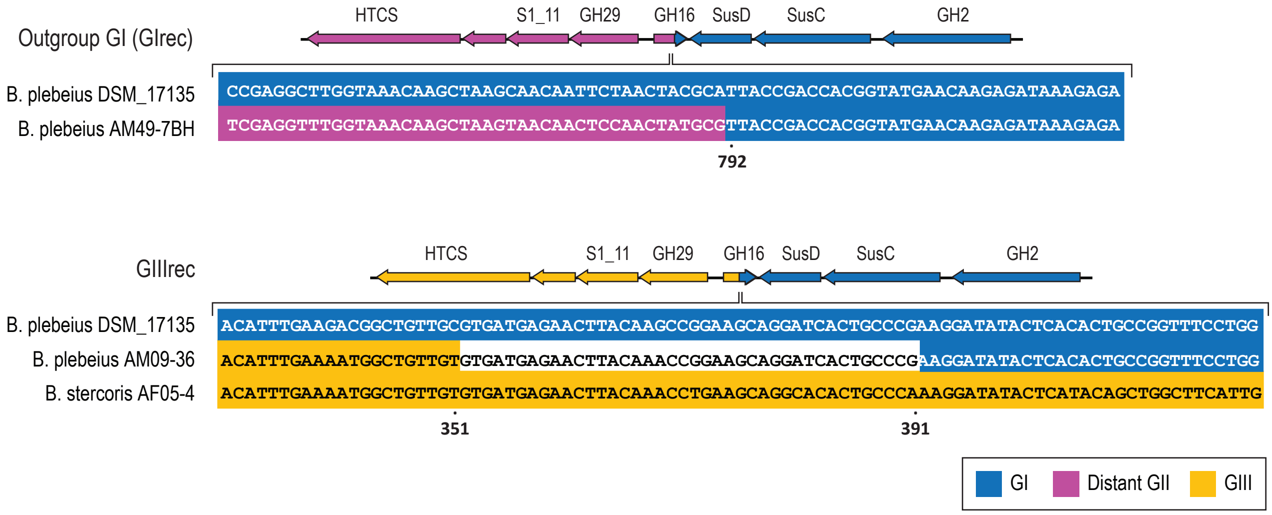


**Figure S6:** Recombination sites observed in the GH16 (Bacple_01703) of the *PUL-PorB*. The Outgroup GI (GIrec) was obtained by the recombination of one gene grouped in GI with one gene distantly related to the GII group (top). Recombination of genes of the groups GI/GIII produced GIIIrec (bottom).

```
               *  ******** *****************    ********  ********
GI       339-  AGACGGCTGTTGCGTGATGAGAACTTACAAGCCGGAAGCAGGATCACTGCCCG -392
GII            GAATGGCTGTTGTGTGATGAGAACTTACAAACCGGAAGCAGGCACACTGCCCG
GIIIrec        AAATGGCTGTTGTGTGATGAGAACTTACAAACCGGAAGCAGGATCACTGCCCG
GIII           AAATGGCTGTTGTGTGATGAGAACTTACAAACCTGAAGCAGGCACACTGCCCA
```

**Figure S7:** Sequences of 54 nucleotides characteristic of the different groups of PUL-PorB used to Blastn SRA dataset.

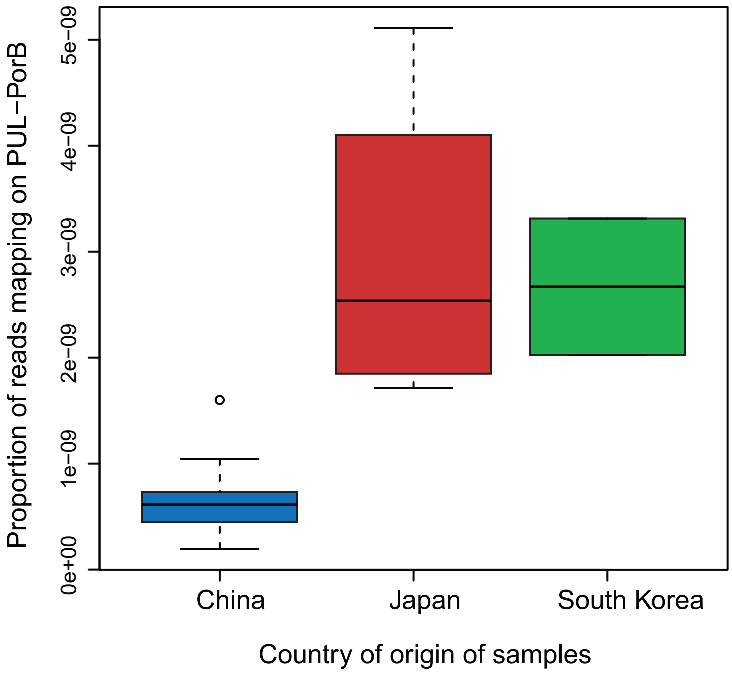


**Figure S8:** Relative abundance of short reads mapping to the 54-nucleotides probes specific to the PUL-PorB in positive individuals across Chinese, Japanese and Koreans projects.

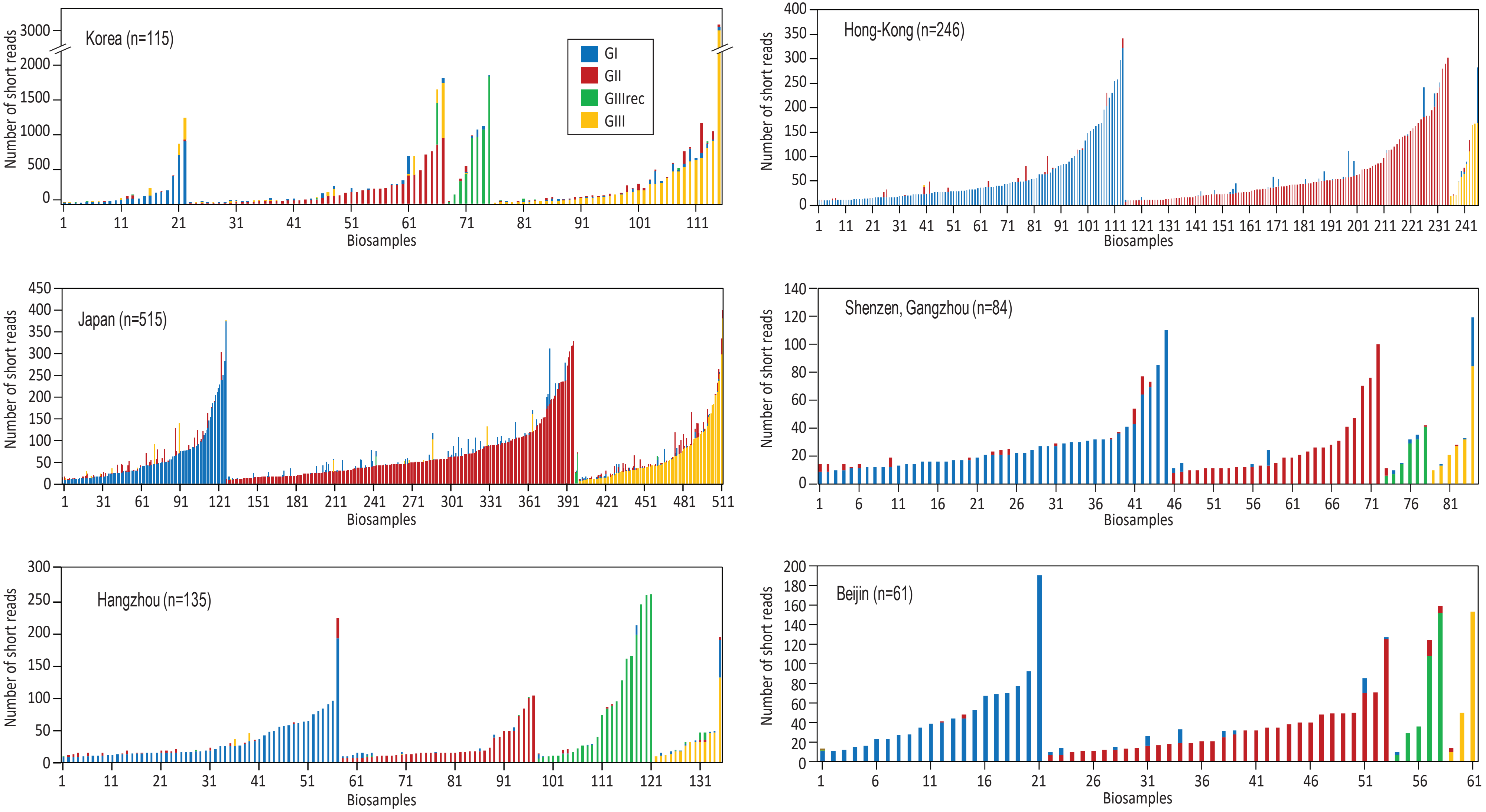


**Figure S9:** The 54 nucleotides sequence characteristic of different group of porphyran PUL were Blastn against the short read sequencing (SRA) of metagnomic data recorded on Japanese, Korean and Chinese populations. The number of detected short reads are indicated for each individuals and were grouped as a function of the geographic location of the sampling. Only individuals for which at least 10 reads were recorded are shown.
